# Phase-specific signatures of wound fibroblasts and matrix patterns define cancer-associated fibroblast subtypes

**DOI:** 10.1101/2022.11.18.516967

**Authors:** Mateusz S. Wietecha, David Lauenstein, Michael Cangkrama, Sybille Seiler, Juyoung Jin, Andreas Goppelt, Manfred Claassen, Mitchell P. Levesque, Reinhard Dummer, Sabine Werner

**Affiliations:** Institute of Molecular Health Sciences, Department of Biology, ETH Zürich, Otto-Stern-Weg 7, 8093 Zürich, Switzerland; Ottobock SE & Co. KGaA; 37115 Duderstadt, Germany; Department of Medicine, Department of Computer Science, Eberhard Karls University, 72076 Tübingen, Germany; Department of Dermatology, University of Zürich Hospital, 8952 Schlieren, Switzerland; College of Dentistry, University of Illinois at Chicago, 801 S. Paulina St., Chicago, IL, 60612, United States of America

**Keywords:** Cancer, cancer-associated fibroblast, collagen, elastin, wound healing

## Abstract

Healing wounds and cancers present remarkable cellular and molecular parallels, but the specific roles of the healing phases are largely unknown. We developed a bioinformatics pipeline to identify genes and pathways that define distinct phases across the time course of healing. Their comparison to cancer transcriptomes revealed that a resolution-phase wound signature is associated with increased severity in skin cancer and enriches for extracellular matrix-related pathways. Comparisons of transcriptomes of early- and late-phase wound fibroblasts vs skin cancer-associated fibroblasts (CAFs) identified an “early-wound” CAF subtype, which localizes to the inner tumor stroma and expresses collagen-related genes that are controlled by the RUNX2 transcription factor. A “late-wound” CAF subtype localizes to the outer tumor stroma and expresses elastin-related genes. Matrix imaging of primary melanoma tissue microarrays validated these matrix signatures and identified collagen- vs elastin-rich niches within the tumor microenvironment, whose spatial organization predicts survival and recurrence. These results identify wound-regulated genes and matrix patterns with prognostic potential in skin cancer.

## INTRODUCTION

Wound healing is a highly complex process that progresses in three distinct and partially overlapping phases – inflammation, proliferation and resolution. Efficient wound healing requires the coordinated actions of specific cell populations and their products [1, 2]. Major spatio-temporal transformations of healing tissue have been described for each phase of repair at the histological level. These changes, which occur in a well-controlled fashion and result in the restoration of tissue homeostasis, are disturbed in non-healing wounds, leading to damage rather than repair [3]. The chronic inflammation in such wounds is also a risk factor for cancer development [4]. Cancer can encapsulate the entire wound healing process run amok, and has been described as a wound that fails to heal [4–6]. Similarities between healing and tumorigenesis include immune and stromal cell-mediated processes such as inflammation, angiogenesis and extracellular matrix (ECM) formation. These processes resolve naturally in acute wounds, but become chronic in tumors, likely due to the persistent production of cytokines and growth factors [4, 5].

Since wound healing is a universal coordinated response, it has been proposed as a marker for describing states or stages of cancer progression [5]. Molecular profiling of fibroblasts, which were exposed to serum to mimic a wound healing response, has demonstrated a high similarity to the expression signatures of some bulk cancer tissue [7]. Indeed, various genes that are regulated by skin wounding are also differentially expressed in malignant tumors, particularly within the tumor microenvironment (TME), and some of them predict survival [4, 5, 8]. However, comparisons of wounds to cancers thus far have not been comprehensive, since the temporal nature of wound healing has not been taken into account. Previous studies have focused on the inflammatory and proliferative phases of wound healing, which feature cellular activation toward proliferative and migratory phenotypes [7–9]. A largely ignored comparison has been to the late phase of repair, which comprises active remodeling of the provisional wound ECM into a mature scar by (myo)fibroblasts [10]. A recent meta-analysis of fibroblasts derived from perturbed states in mice and humans, including wound healing and cancer formation, revealed strong similarities between certain ECM-associated clusters of cancer-associated fibroblasts (CAFs) and wound fibroblasts from early-stage wounds [11]. However, direct comparisons of cancer-derived and wound-derived stromal cells from across the healing spectrum are missing.

Here we present a systematic characterization of the entire wound healing response and unveil gene expression profiles specific to each healing phase. We used these phase-specific genes to make systems-level comparisons to tumors, and derived phase-predictive wound healing genes with prognostic potential in human cancers, with a focus on skin cancer. These genes enrich for ECM pathways, such as collagen and elastin formation, suggesting that wound fibroblast/CAF signatures may predict cancer outcome. Our results identify signatures of early- and late-phase wound fibroblasts in different CAF subtypes. These wound-related CAF subtypes exist in spatially distinct niches within the tumor stroma and produce divergent repertoires of collagen and elastin fibers organized in unique ECM architectures, which have prognostic potential. Our results pave the way for the development of innovative wound healing and matrix-based diagnostic tools to predict cancer outcome.

## RESULTS

### Identification of phase-specific wound healing genes

To identify molecular parallels between malignant tumors and skin wounds at different stages, it is important to identify genes that are specifically regulated in individual phases of healing. A meta-analysis of microarray-based wound healing profiles had identified genes, which changed their expression over the time-course of healing in mice [12], but many of them were up-regulated in multiple phases (Fig. S1A) and could not be used as reliable phase markers. Therefore, we developed a bioinformatics pipeline to identify wound phase-specific gene signatures by re-analysis of two independent datasets of mouse excisional wounds at different stages [13, 14] (Fig. 1A; Table S1). There were clear correlations between time points from both studies at the whole transcriptome level (Fig. 1B). We binned the samples according to their correlations to intact skin or inflammatory, proliferative, or resolution phases of healing, and – using the new Biopeak R package (see Experimental Procedures) – extracted genes, which showed specific and robust up-regulation in only one phase (Fig. 1A). We identified 1211 such phase-specific genes (Fig. 1C; Table S2), which showed strong within-phase correlation and between-phase anti-correlation (Fig. 1D). Their comparison to previously identified healing genes showed phase-congruent overlap in the inflammatory and resolution phases (Fig. S1B). These phase-specific genes were highly enriched for pathways characteristic for each phase of healing, such as reactive oxygen species- (ROS) and cytokine-mediated pathways in the inflammatory phase; cell proliferation, migration and growth factor signaling in the inflammation-to-proliferation phases; and ECM-related pathways in the proliferative-to-resolution phases (Figs. 1E, S1C and S1D; Table S2). Specifically, “assembly of collagen fibrils” was enriched in the proliferative and resolution phases, whereas “degradation of the ECM”, “ECM proteoglycans”, “integrin cell surface interactions” and “molecules associated with elastic fibers” were more enriched in the resolution phase of healing (Fig. 1E).

**Fig. 1.**
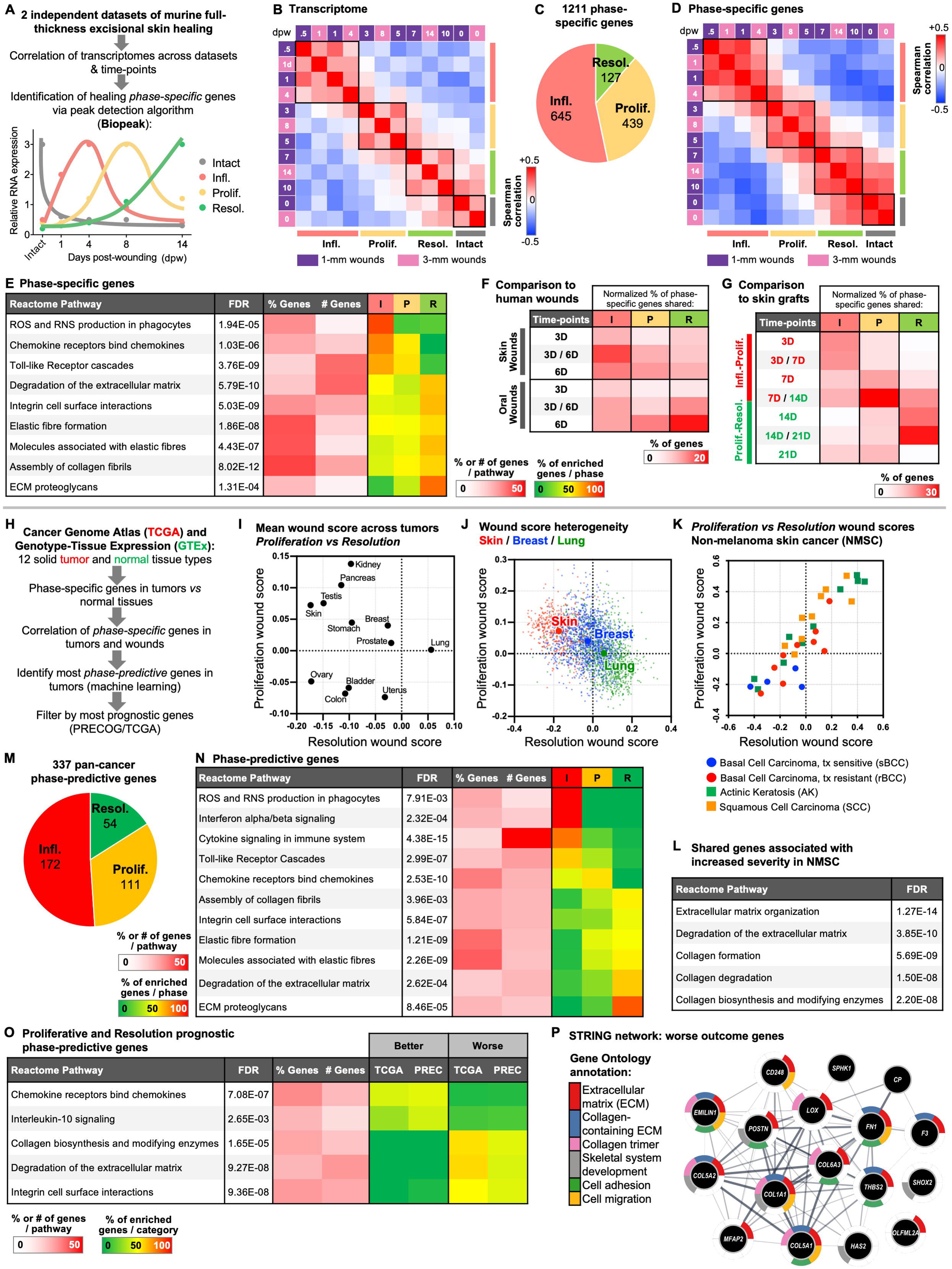
Identification of wound healing phase-specific genes and tumor phase-predictive genes. (A) Bioinformatics workflow for the identification of healing phase-specific genes by peak detection. (B) Spearman correlation matrix from the shared transcriptome of samples from across the healing spectrum. (C) Breakdown of the healing phase-specific genes into the inflammatory (I), proliferative (P) and resolution (R) phases according to peak detection. (D) Spearman correlation matrix using only the phase-specific genes of samples from across the healing spectrum. (E) Functional enrichment analysis of healing phase-specific genes, showing selected Reactome pathways, statistical significance, the percentage (%) and number (#) of genes enriched per pathway, and the percentage of enriched genes per healing phase. (F) Comparison of healing phase-specific genes to human skin and oral wound healing datasets, showing an input-normalized percentage (%) of shared phase-specific genes with time-resolved human gene sets. (G) Comparison of healing phase-specific genes to human skin graft datasets, showing an input-normalized percentage (%) of shared phase-specific genes with time-resolved human gene sets, separated into the inflammatory-proliferative and proliferative-resolution healing phases. (H) Bioinformatics workflow for the identification of healing phase-predictive genes expressed in cancer tissue with potential prognostic relevance based on analysis of TCGA/GTEx and PRECOG databases. (I) Plot of mean proliferative and resolution phase wound scores across solid tumor types, showing high inter-tumor type heterogeneity. (J) Plot of individual proliferative and resolution phase wound scores in skin (melanoma), breast and lung tumor samples, showing high intra-tumor type heterogeneity. Large circles indicate mean wound score for each tumor type. (K) Plot of individual proliferative and resolution wound scores in non-melanoma skin cancer (NMSC), including squamous cell carcinoma (SCC) and basal cell carcinoma (BCC), and SCC precursor lesions (Actinic Keratosis (AK)) [18, 19]. (L) Functional enrichment analysis of shared NMSC genes associated with increased severity. (M) Breakdown of tumor phase-predictive genes into the inflammatory (I), proliferative (P) and resolution (R) phases using machine learning algorithm. (N) Comparison of pan-cancer prognostic proliferative and resolution phase-predictive genes according to the TCGA and PRECOG databases. (O) Functional enrichment analysis of pan-cancer prognostic proliferative and resolution phase-predictive genes, showing selected Reactome pathways, statistical significance, the percentage (%) and number (#) of genes enriched per pathway, and the percentage of enriched genes per database-specific patient outcome category. (P) Functionally annotated StringDB gene-gene interaction network of the shared phase-predictive genes associated with worse outcome. Infl, I: Inflammatory; Prolif, P: Proliferative; Resol, R: Resolution; dpw: days post-wounding; FDR: false discovery rate; TCGA: The Cancer Genome Atlas; PREC: PRECOG.

To validate our findings in the human setting, we compared the phase-specific genes to datasets of paired oral and skin wounds from healthy adults at 3 and 6 days post-injury [15] (Fig. S1E; Table S1). Skin wounds at both time-points were still mostly in the inflammatory phase, whereas the oral wounds had already transitioned to the resolution phase at 6 days post-injury (Fig. 1F; Table S3), which is consistent with their more rapid healing [13]. Since there are currently no published datasets of human excisional skin wounds that include the full time-course of healing, we used transcriptome data from split-thickness skin grafts sampled at early or late time-points [16] (Table S1). We derived the genes, which were up-regulated after human skin grafting in a time-resolved fashion (Fig. S1F; Table S3), and compared them to the healing phase-specific genes. There was a strong overlap of inflammatory and resolution phase-specific genes with early-stage (3-day and/or 7-day) and late-stage (14-day and/or 21-day) grafts, respectively, and proliferative phase-specific genes were enriched in the transition of early-stage to late-stage (7-day and 14-day) grafts (Fig. 1G). Examples of shared phase-specific genes include inflammatory mediators and early collagen regulators in the inflammatory phase; late-inflammatory mediators and collagen production regulators in the proliferative phase; and fibrillar collagens, fibrosis-associated glycoproteins, and elastin-associated genes in the resolution phase (Fig. S1G). These results highlight the time-resolved molecular similarities between wound healing in mice and humans.

### Phase-specific wound healing genes may predict cancer prognosis

To compare our healing phase-specific genes with gene expression signatures in human cancers, we developed a bioinformatics pipeline using normalized transcriptomes of tumors from “The Cancer Genome Atlas” (TCGA) and respective healthy tissue samples from “Genotype-Tissue Expression” (GTEx) (Fig. 1H). We used GTEx since the tumor-adjacent samples in the TCGA are significantly different at the transcriptome level compared to tissue samples from healthy individuals [17]. Gene expression in 12 tumor types was systematically correlated to phase-specific wound healing genes using their human orthologues. We defined phase-specific wound scores as Spearman correlations of a tumor to the inflammatory, proliferative, or resolution phase of wounds, a robust approach especially for skin samples (see Experimental Procedures). The distribution of wound scores between and within tumor types showed a high degree of inter- and intra-tumor type heterogeneity, including in skin melanoma (Figs. 1I and 1J). We also tested for the presence of wound score signatures in non-melanoma skin cancer (NMSC) datasets, including cutaneous squamous cell carcinoma (cSCC), their actinic keratosis (AK) precursor lesions [18] and basal cell carcinoma (BCC) [19] (Table S1), which revealed heterogeneity in proliferative and resolution-phase wound scores in these lesions (Fig. 1K). However, when we compared the phase-specific genes with respect to skin cancer severity by identifying the genes upregulated in SCC compared to AK and those upregulated in treatment-resistant *vs* treatment-sensitive BCC, we found that overexpression of the wound-regulated genes involved in collagen formation correlates with tumor severity in both comparisons (Fig. 1L). The relevant overexpressed genes include *COL1A1* (encoding collagen α1(I)) and *PLOD2* (encoding procollagen-lysine, 2-oxoglutarate 5-dioxygenase), as well as *FN1* (encoding fibronectin), *TNC* (encoding tenascin C), and *INHBA,* which encodes the INHBA subunit of activin A (Fig. S1H). The latter is a potent regulator of the expression of collagens and other ECM genes in skin fibroblasts [20].

We then used machine learning to identify the most phase-predictive genes in TCGA tumors, *i.e.* the wound phase-specific genes that are up-regulated in tumors and most strongly contribute to an increase in their phase-specific wound scores (see Experimental Procedures). After aggregating the best-scoring features across datasets, we identified 337 phase-predictive genes, which spanned the whole spectrum of healing (Fig. 1M; Table S4). We re-calculated the TCGA tumor wound scores using only the phase-predictive genes, which resulted in shifting of the wound scores and an increase in intra-tumor type heterogeneity, including in skin melanoma (Fig. S1I). This heterogeneity enabled us to stratify the patient samples according to their respective wound scores and to test whether the wound scores themselves are prognostic in individual tumor types. Indeed, the resolution phase wound scores were statistically predictive of worse outcome in primary and metastatic melanoma (Fig. S1J).

These findings suggest that wound-associated gene expression signatures are important contributors to tumor severity, as was first shown by the analysis of the fibroblast core serum response (CSR) signature [7]. We thus compared the CSR signature with the phase-specific and phase-predictive genes, which showed moderate overlap of the CSR with inflammatory and proliferative, but not resolution, phase-specific genes (Fig. S1K). Therefore, the healing phase in which a gene is expressed in a specific cancer type is of key relevance to its prognostic potential.

The phase-predictive genes enriched for wound-associated biological processes that are also shared by tumors, including ROS- and cytokine-mediated pathways in the inflammatory phase, growth factor signaling and ECM formation in the proliferative phase, and ECM formation/remodeling in the resolution phase-predictive gene sets (Fig. 1N; Table S4). To test for clinical significance of individual wound-associated genes, we analyzed the phase-predictive genes using the PRECOG and TCGA databases across all solid tumors [21] (Table S5). We focused on proliferative and resolution phase-predictive genes, since the inflammatory phase genes were highly enriched in inflammation-associated pathways that had already been the focus of previous studies [7, 9]. The proliferative and resolution phase-predictive genes were concordant with respect to their predicted clinical cancer outcome in both databases (Fig. S1L). Functional enrichment analysis of these phase-predictive and prognostic genes showed that those correlating with better prognosis were enriched for immune-related pathways, whereas those correlating with worse prognosis were enriched for ECM-related pathways (Fig. 1O). This result was confirmed when we focused the prognostic analysis on skin melanoma, with genes enriching for “collagen formation” or “elastic fibre formation” predicting worse or better prognosis, respectively (Fig. S1M, Table S5).

In order to visualize the known connections between phase-predictive genes that were correlated with worse clinical outcome, we generated a STRING network that incorporated their predicted protein-protein interactions and showed their shared pathway enrichments. These genes were highly connected and commonly enriched for the GO terms “ECM”, “collagen-containing ECM”, “collagen trimer”, “skeletal system development”, “cell adhesion” and “cell migration” (Fig. 1P). Functional enrichment for cell types showed that these genes are highly enriched in curated fibroblast signatures (Fig. S2A), strongly suggesting that the prognostic wound-associated genes in tumors are mainly expressed by CAFs. Therefore, we focused the following analyses on skin wound fibroblasts and CAFs.

### Identification of wound fibroblast phase-specific genes

Since we showed that time-resolved wound signatures could act as markers for whole tumor states, it is possible that genes that are overexpressed in wound fibroblasts at different phases of healing could act as markers for CAF subtypes, which are highly heterogeneous within and across tumor types [22, 23]. Therefore, we first correlated the inflammatory, proliferative and resolution phase wound genes with the gene expression signatures of wound fibroblasts from different phases of healing by re-analyzing the transcriptomes of wound fibroblasts from early- and late-stage large excisional mouse wounds [24] (Table S1). Their correlation with the expression profiles of whole wounds identified two main clusters of fibroblast signatures, one correlating with the early (inflammatory-to-proliferative) phases and one correlating with the late (proliferative-to-resolution) phase of healing (Fig. 2A). The fibroblast time points did not only cluster with each other, but also with their corresponding phases in whole wounds, pointing to a strong contribution of this cell type to expression signatures in bulk-sequenced wound tissues.

**Fig. 2.**
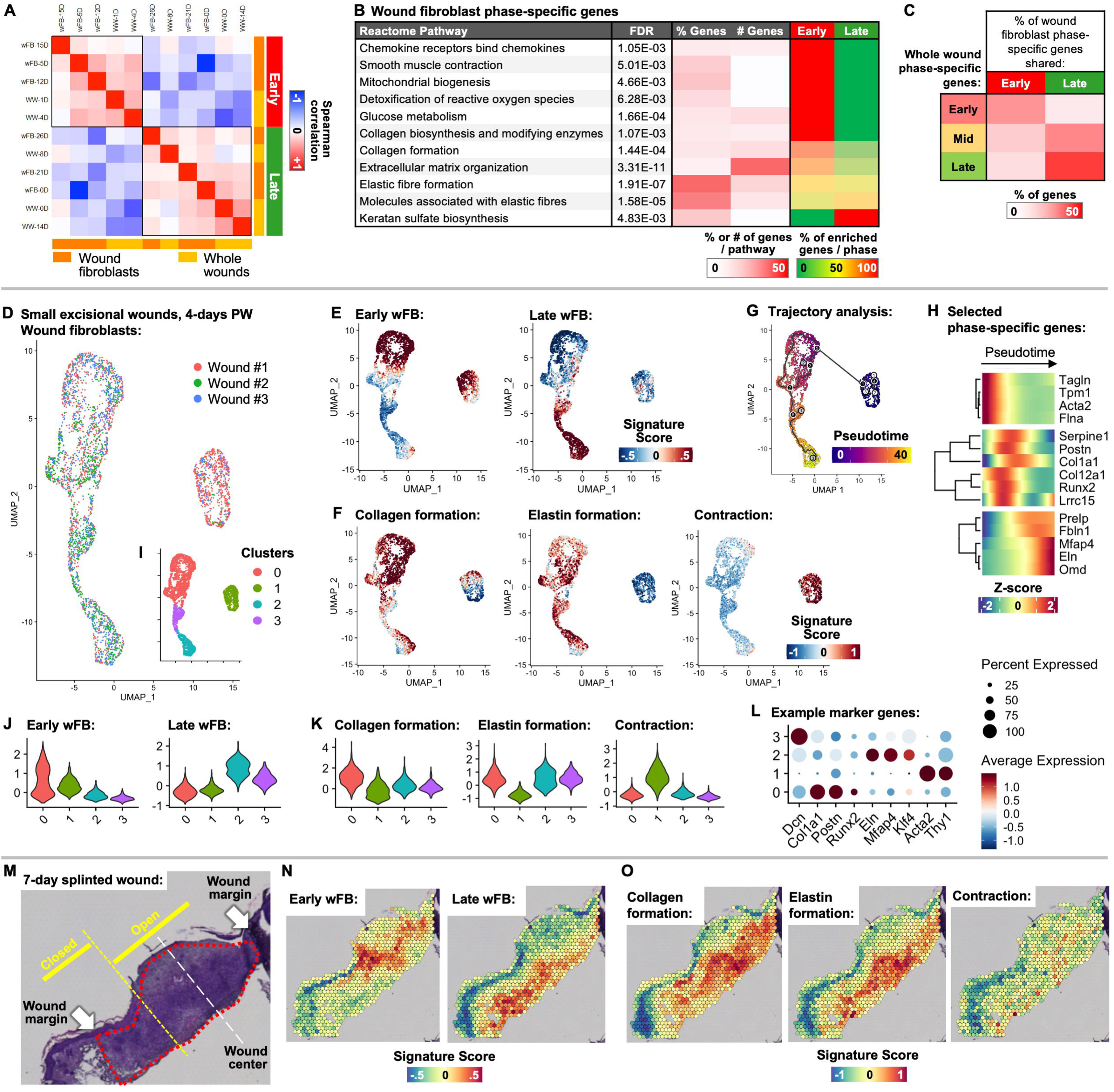
Identification of wound fibroblast phase-specific genes. (A) Spearman correlation matrix from the shared transcriptome of whole mouse wound (WW) and wound fibroblast (wFB) samples from across the healing spectrum. (B) Functional enrichment analysis of wFB phase-specific genes, showing selected Reactome pathways, statistical significance, the percentage (%) and number (#) of genes enriched per pathway, and the percentage of enriched genes per healing phase. (C) Comparison of wFB phase-specific genes to WW phase-specific genes, showing a percentage (%) of shared genes. (D-G) UMAP plots showing mouse wFB cells from small excisional wounds at 4-days post-wounding (PW) colored by dataset of origin (D), distribution of Early (left) and Late (right) wFB phase-specific gene signature scores (E), distribution of selected Reactome pathway gene signature scores (F) or Monocle-based trajectory path and distribution of pseudotime values (G). (H) Clustered heatmap showing normalized expression of selected wFB phase-specific genes along the trajectory pseudotime. (I) UMAP plot showing mouse fibroblast cells in (D) colored by transcriptional cluster number. (J) Violin plots showing the distribution of early and late wFB phase-specific gene signature scores in wFB clusters identified in (I). (K) Violin plots showing the distribution of selected Reactome pathway gene signature scores in wFB clusters identified in (I). (L) Dot plot showing the average normalized expression (node color) and percent expression (node size) of selected wFB phase-specific genes in wFB clusters identified in (I). (M) Diagram of the 7-day PW splinted excisional wound section analyzed via spatial transcriptomics, indicating locations of major wound-associated morphological features. (N) Feature plot showing the spatial distribution of early (left) and late (right) wFB phase-specific gene signature scores overlaying the wound section shown in (M). (O) Feature plot showing the spatial distribution of selected Reactome pathway gene signature scores overlaying the wound section shown in (M).

We binned the wound fibroblast samples according to their correlations into “early” (overexpressed in the inflammatory-to-proliferative phase wounds) and “late” categories (overexpressed in proliferative-to-resolution phase wounds), and – using the Biopeak R package (see Experimental Procedures) – extracted genes, which showed specific and robust up-regulation in one of the temporal categories (Table S6). Pathways enriched in the early wound fibroblasts include inflammation, contraction, glycolysis, and collagen biosynthesis; ECM organization and collagen formation were enriched in both early and late wound fibroblasts; and keratan sulfate biosynthesis and elastin formation were more enriched in the late group (Fig. 2B; Table S6). Network Reactome pathway analysis identified the genes from the diverse ECM-related pathways that are enriched in the early and late wound fibroblast signatures. In the early group were collagen-associated *Plod2*, dual-associated *Loxl2* (encoding lysyl oxidase like 2) and elastin-associated *Fbln2* (fibulin-2), while the late group included *Eln* (elastin) itself, the elastin-associated *Mfap4* (microfibril-associated glycoprotein 4 precursor) and *Fbln5* (Fig. S2B). High numbers of the early fibroblast genes were shared with genes expressed during the inflammatory and proliferative phase in whole wounds, while in the late fibroblasts most genes were shared with genes expressed in proliferative and resolution phase whole wounds (Fig. 2C). This analysis showed that fibroblasts from early- and late-phase wounds express gene signatures encoding divergent ECM repertoires.

To investigate whether this early *vs* late wound fibroblast heterogeneity exists within and across wounds, we made use of available single cell RNA sequencing (scRNA)-seq datasets of murine wound fibroblasts (Table S1). We first re-analyzed cells from day-4 small mouse excisional wounds [25] (Fig. 2D) and calculated the early and late wound fibroblast signature scores across this dataset. We found that about half of the cells enriched for the early or late signatures, with a gradient of early-to-late signature scores from the top to bottom of the UMAP plot (Fig. 2E). Signature scores of major Reactome pathways enriched in bulk-sorted early *v*s late wound fibroblasts showed a gradient of predicted phenotypes in the scRNA-seq dataset, with “collagen formation”, “elastin formation” and “smooth muscle contraction” enriching on the top, bottom and right sides of the UMAP plot, respectively (Fig. 2F). We then performed trajectory analyses to investigate potential differentiation pathways in wound fibroblasts. Monocle and Cytotrace analyses predicted a path from the area of cells with high early wound fibroblast signature scores to the area of cells with high late wound fibroblast signatures scores (Figs. 2G and S2D). Plotting of selected wound fibroblast phase-specific gene expression along the Monocle pseudotime revealed a cluster of genes associated with contraction (e.g. *Acta2* (encoding alpha smooth muscle actin (α-SMA)) and *Tpm1* (tropomyosin 1), leading to genes associated with collagen formation (e.g. *Col1a1*, *Postn* (periostin) and *Lrrc15* (leucine rich repeat containing 15)) and finally to genes associated with elastin formation (e.g. *Eln*, *Mfap4* and *Fbln1*) (Fig. 2H). Interestingly, the pseudotime expression of the gene encoding Runx2 (RUNX family transcription factor 2) was strongly associated with the early wound fibroblast and collagen formation cluster of genes (Fig. 2H). We repeated this analysis on integrated scRNA-seq datasets of fibroblasts from early- and late-phase large excisional wounds [26, 27] and obtained similar results with regard to the distribution of early and late wFB signature scores and especially the association of the late wFB enriched cells with those with high “elastin formation” pathway scores (Figs. S2E-S2I).

Cluster-based analysis of the small excisional wound scRNA-seq dataset identified four transcriptionally-distinct clusters (Fig. 2I). The early wound fibroblast signature was especially enriched in clusters #0 and #1 (Fig. 2J), and these clusters also enriched for collagen formation and contraction, respectively (Fig. 2K). The late wound fibroblast signature was enriched in clusters #2 and #3 (Fig. 2J) and also enriched for elastin formation (Fig. 2K). Early wound fibroblast and collagen formation genes, e.g. *Col1a1* and *Postn,* were strong markers for cluster #0 along with *Runx2*, whereas elastin formation genes (*Eln* and *Mfap4*) were strong markers for cluster #2 along with the gene encoding the transcription factor Klf4 (Fig. 2L, Table S7). Cluster #1 was marked by high expression of contraction associated genes, such as *Acta2*, while Cluster #3 was characterized by high expression of genes associated with fibroblasts from unwounded skin, e.g. *Dcn* (decorin) (Fig. 2L).

Finally, we re-analyzed spatial transcriptomics data of splinted excisional mouse wounds [28], focusing on a 7-day wound that had clearly defined margins and spatially distinct open and closed wound areas, with a gradient of healing within a single section (Fig. 2M). The early wound fibroblast signature was defined to the early granulation tissue immediately surrounding the open part of the wound, whereas the late wound fibroblast signature was enriched in deeper areas of the granulation tissue where the wound is already covered by neo-epidermis (Fig. 2N). The collagen formation and contraction pathway signatures were high in the early fibroblast-enriched granulation tissue area, while the elastin formation pathway signature was high in the late fibroblast-enriched healed wound areas (Fig. 2O). These data collectively show that fibroblasts take on early- *vs* late-phase wound signatures within and between wounds that are spatially defined and encode for divergent collagen *vs* elastin ECM repertoires.

### Wound phase-specific genes are enriched in CAF subgroups of mouse and human skin cancers

We next compared the transcriptomes of early and late wound fibroblasts with those of bulk-sequenced CAFs derived from a variety of solid tumors (Table S1). The batch-corrected data from all studies were merged into one matrix, filtered for phase-specific wound fibroblast signature genes, and the CAFs were compared with early and late wound fibroblasts (see Experimental Procedures). This analysis identified two groups of CAFs corresponding to early and late phase wound fibroblasts (Fig. 3A). There was some, but not exclusive tissue-specific clustering, indicating minimal batch effects based on organ or study of origin. One exception were the fibroblasts from murine human papilloma virus (HPV8)-induced papillomas (see below), which clustered together, and highly correlated to the earliest wound fibroblast signature.

**Fig. 3.**
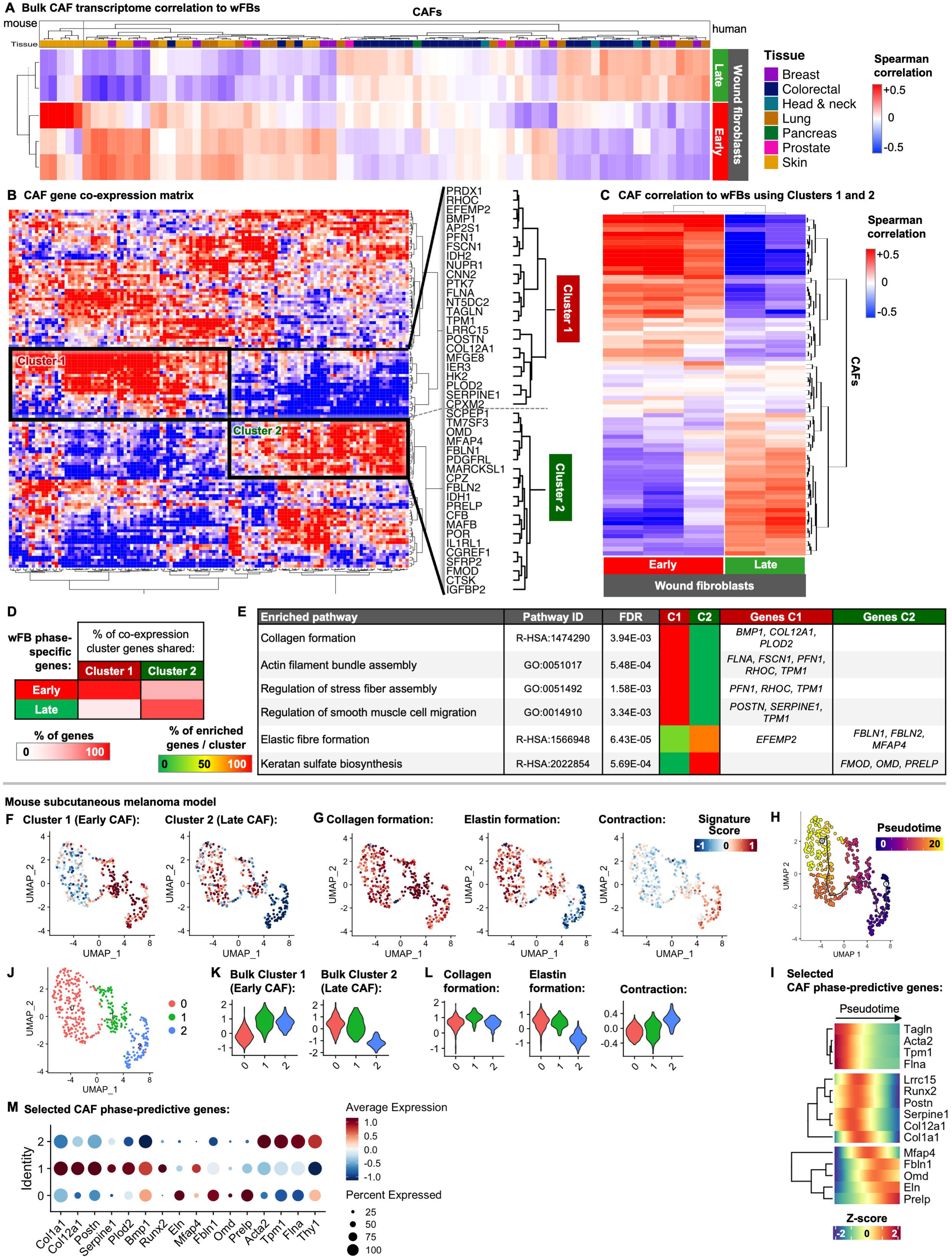
Comparison of phase-resolved wound fibroblasts with CAFs. (A) Spearman correlation matrix of wFBs from across the healing spectrum (rows) and CAFs derived from several bulk datasets comprising seven tissue types (columns) using wFB phase-specific genes (datasets are listed in Table S1). (B) Gene-gene Spearman correlation matrix of pan-CAF datasets, identifying two mutually-exclusive gene clusters, termed co-expression clusters 1 and 2. (C) Detailed Spearman correlation matrix of wound fibroblasts (wFBs) from across the healing spectrum (rows) and CAFs derived from several bulk datasets comprising seven tissue types (columns) using only genes in co-expression clusters 1 and 2. (D) Comparison of co-expression gene clusters 1 and 2 to wFB phase-specific gene sets, showing percentage (%) of shared genes. (E) Functional enrichment analysis of co-expression cluster genes, termed CAF phase-predictive genes, showing selected Gene Ontology (GO) and Reactome pathways, statistical significance, the percentage of enriched genes per cluster, and lists of enriched genes per cluster. (F-H) UMAP plots showing distribution of early (left) and late (right) CAF phase-predictive gene signature scores in experimental mouse subcutaneous melanoma model CAFs (F), (selected Reactome pathway gene signature scores (G) or Monocle-based trajectory path and distribution of pseudotime values (H). (I) Clustered heatmap showing normalized expression of selected CAF phase-predictive genes along the trajectory pseudotime. (J) UMAP plot showing CAFs from an experimental mouse subcutaneous melanoma model colored by transcriptional cluster number. (K) Violin plots showing the distribution of early (left) and late (right) CAF phase-predictive gene signature scores in CAF clusters identified in (J). (L) Violin plots showing the distribution of selected Reactome pathway gene signature scores in CAF clusters identified in (J). (M) Dot plot showing the average normalized expression (node color) and percent expression (node size) of selected CAF phase-predictive genes in CAF clusters identified in (J). C#: cluster number; FDR: false discovery rate.

To identify the genes that contribute most to classifying CAFs into either early- or late-phase wound fibroblasts, we investigated gene-gene co-expression patterns across the meta-CAF dataset and identified two mutually-exclusive gene signature clusters (Fig. 3A). Cluster 1 contained 24 genes, including *POSTN*, *PLOD2*, *SERPINE1, TPM1* and *LRRC15*, while Cluster 2 contained 20 genes, including *MFAP4*, *FBLN1*, *PRELP* (proline and arginine rich end leucine rich repeat protein) and *OMD* (osteomodulin) (Table S8). We then used these genes to re-classify the CAFs, which identified two distinct clusters highly corresponding to early and late wound fibroblast signatures (Fig. 3C). Comparing these cluster genes with the wound fibroblast phase-specific genes confirmed the association of Cluster 1 with the early and Cluster 2 with the late wound fibroblast signature (Fig. 3D). Functional enrichment analyses of these CAF phase-predictive genes showed exclusive enrichment for “collagen formation,” “stress fiber assembly” and “migration” in Cluster 1, and for “elastic fiber formation” and “keratan sulfate biosynthesis” in Cluster 2 (Fig. 3E; Table S8).

To investigate whether CAF heterogeneity exists within and across experimental tumors, we re-analyzed published scRNA-seq data of CAFs from mouse subcutaneous melanomas [29] (Table S1). We calculated the early and late CAF signature scores across this dataset, showing that about half of the cells enriched for the early or late CAF signatures, with a gradient of early-to-late signature scores from the right to left of the UMAP plot (Fig. 3F). Signature scores of major Reactome pathways enriched in bulk-sorted early *vs* late CAFs showed a gradient of predicted phenotypes in the scRNA-seq dataset, with “collagen formation”, “elastin formation” and “smooth muscle contraction” enriching at the center, left and right of the UMAP plot, respectively (Fig. 3G). We then performed trajectory analyses to investigate potential differentiation pathways of the experimental CAFs. Monocle analysis predicted a path from the area of cells with high early CAF signature scores to the area of cells with high late CAF signatures scores (Fig. 3H). Plotting of selected CAF phase-predictive gene expression along the pseudotime revealed a cluster of genes associated with contraction, leading to genes associated with collagen formation and finally to genes associated with elastin formation (Fig. 3I). Interestingly, the pseudotime expression of *Runx2* was also associated with the early wound fibroblast and collagen formation cluster of genes in CAFs, which is consistent with the wound healing data (Fig. 2H).

Cluster-based analysis of the subcutaneous melanoma scRNA-seq dataset identified three transcriptionally distinct clusters (Fig. 3J). The early CAF signature was especially enriched in clusters #1 and #2 (Fig. 3K), which also enriched for collagen formation or contraction, respectively (Fig. 3L). The late CAF signature was enriched in clusters #0 and #1, which also enriched for elastin formation (Figs. 3K and 3L). Differential expression analyses showed that early CAF and collagen formation genes were strong markers for cluster #1 along with *Runx2*, whereas elastin formation genes were strong markers for cluster #0 (Fig. 3M; Table S7). Cluster #2 was marked by high expression of contraction-associated genes.

These results suggest the existence of two CAF subtypes encoding divergent ECM repertoires: an “early wound”-like CAF subtype that promotes contraction and collagen fiber formation, and a “late wound”-like CAF subtype that promotes elastic fiber formation.

### Meta-analysis of single-cell sequenced human CAFs from skin tumors

To investigate whether CAF heterogeneity exists within and across human skin tumors, we undertook a comprehensive meta-analysis of available scRNA-seq datasets (Table S1). We re-processed scRNA-seq data of CAFs from BCC [30], skin SCC [31], and skin melanoma [32]. We calculated the early and late CAF signature scores across these datasets, showing that about half of the cells enriched for the early or late CAF signatures in all three types of tumors (Figs. 4A, 4B, and S3A). Signature scores of major Reactome pathways enriched in bulk-sorted early *vs* late CAFs showed gradients of predicted phenotypes in the scRNA-seq datasets, with collagen formation and contraction enriching in the areas of high early CAF scores, and elastin formation enriched in the areas of high late CAF scores in BCC, SCC and melanoma (Figs. 4C, 4D, and S3B). We then performed trajectory analyses on the BCC and SCC datasets to investigate potential differentiation pathways in CAFs. In both tumor types, Monocle predicted paths from the area of cells with high early CAF signature scores to the area of cells with high late CAF signatures scores (Figs. 4E and 4F). Plotting of selected CAF phase-predictive gene expression along the pseudotime revealed a cluster of genes associated with fibroblast contraction, leading to genes associated with collagen formation and finally to genes associated with elastin formation (Figs. 4G and 4H in BCC and SCC, respectively). Again, the pseudotime expression of *RUNX2* was associated with the early wound fibroblast and collagen formation clusters of genes in both BCC and SCC.

**Fig. 4.**
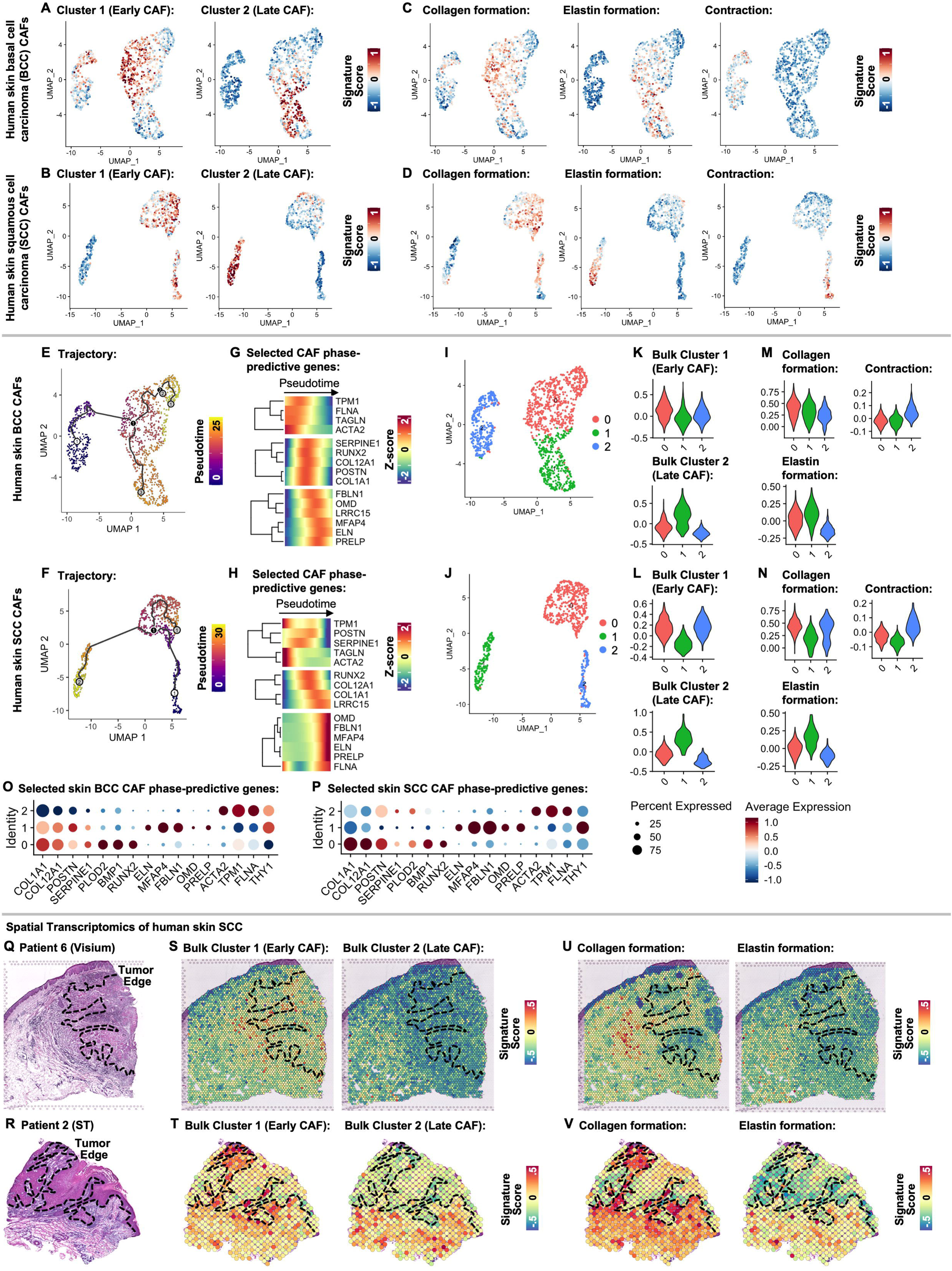
Early and late CAFs in human skin cancers. (A, B) UMAP plots showing distribution of early (left) and late (right) CAF phase-predictive gene signature scores in BCC-derived CAFs (A) or skin SCC-derived CAFs (B). (C, D) UMAP plots showing distribution of selected Reactome pathway gene signature scores in BCC CAFs (C) or SCC CAFs (D). (E, F) UMAP plots showing Monocle-based trajectory path and distribution of pseudotime values in BCC CAFs (E) or SCC CAFs (F). (G, H) Clustered heatmaps showing normalized expression of selected CAF phase-predictive genes along the trajectory pseudotime in BCC CAFs (G) or SCC CAFs (H). (I, J) UMAP plots showing BCC CAFs (I) or SCC CAFs (J) colored by transcriptional cluster number. (K, L) Violin plots showing the distribution of early (left) and late (right) CAF phase-predictive gene signature scores in BCC or SCC CAF clusters identified in (I) or in (J), respectively. (M, N) Violin plots showing the distribution of selected Reactome pathway gene signature scores in BCC or SCC CAF clusters identified in (I) or in (J), respectively. (O, P) Dot plot showing the average normalized expression (node color) and percent expression (node size) of selected CAF phase-predictive genes in BCC CAF clusters identified in (I) or in SCC CAF clusters identified in (J), respectively. (Q, R) Diagram of SCC section analyzed via Visium (Q) or early Spatial Transcriptomics (R); dashed lines indicate tumor margin. Scale bars = 500 μm. (S, T) Feature plots showing the spatial distribution of early (left) and late (right) CAF phase-predictive gene signature scores overlaying the SCC section shown in (Q) or (R), respectively. (U, V) Feature plots showing the spatial distribution of selected Reactome pathway gene signature scores overlaying the SCC section shown in (Q) or in (R), respectively.

Cluster-based analyses of the SCC, BCC and melanoma scRNA-seq datasets identified three transcriptionally-distinct clusters (Figs. 4I, 4J and Fig S3C). In BCC and SCC, the early CAF signature was enriched in clusters #0 and #2 (Figs. 4K and 4L), which also enriched for collagen formation and contraction, respectively (Figs. 4M and 4N). In contrast, the late CAF signature was enriched in cluster #1 in BCC and SCC (Figs. 4K and 4L), which also enriched for elastin formation (Figs. 4M and 4N). Differential expression analyses showed that early CAF and collagen formation genes, such as *COL1A1* and *POSTN,* were strong markers for cluster #0 along with *RUNX2* especially in SCC, whereas elastin formation genes such as *ELN* and *MFAP4* were strong markers for cluster #1 in BCC and SCC (Figs. 4O and 4P; Table S9). Cluster #2 was marked by high expression of contraction associated genes. Similar results were obtained in the melanoma dataset: cluster #2 enriched for early CAF and collagen formation signatures, while cluster #0 enriched for late CAF and elastin formation signatures (Figs. S3D and S3E).

Analyses using scRNA-seq datasets from breast, lung, colorectal, head and neck SCC, and ovarian cancers showed remarkable similarities in early *vs* late CAF signature distributions to those in skin tumors (Figs. S4A-S4E). Integration of all CAF datasets (Fig. S4F) confirmed the strong polarization of early *vs* late CAF signatures and their associated pathway enrichments in collagen *vs* elastin formation (Fig. S4G), respectively, along with their respective marker gene expression, across different tumor types (Figs. S4H-S4J; Table S9).

In addition to the existence of several CAF subtypes, their localization within the tumor is of likely importance. Therefore, we re-analyzed spatial transcriptomics (ST) data of skin SCC [31], which included tumor sections from several patients. We used two ST platforms and calculated the early and late CAF signature scores within the spatial voxels. In sections from patients 6 and 2, which had large stromal areas beyond the tumor margins (Figs. 4Q and 4R), the early CAF signatures were high in close vicinity to the tumor margins, whereas the late CAF signatures were high in stroma further distant to the tumor cells (Fig. 4S and 4T). In both patients, the collagen *vs* elastin formation signatures were high in the early *vs.* late CAF-enriched tumor stromal areas (Figs. 4U and 4V). The co-localization of early CAF and collagen signatures was also observed in sections from additional SCC patients (Figs. S5A-S5C).

These data collectively show that within and between skin tumors, fibroblasts take on early- *vs* late-phase wound signatures that are spatially defined and encode divergent collagen *vs* elastin ECM repertoires.

### Direct similarities between early wound fibroblasts and CAFs

To directly compare early-stage wound and tumor fibroblasts in animals of the same genetic background, we FACS-sorted fibroblasts from normal skin and early-proliferative (5-day) excisional skin wounds of wild-type mice and from skin papillomas that spontaneously formed on the back of mice expressing the oncogenes of HPV8 in keratinocytes [33], using antibodies against the pan-fibroblast marker platelet-derived growth factor receptor alpha (*Pdgfra*; *CD140a*) [20] (Fig. S6A for gating strategy; Fig. S6B for papilloma histology; Table S1). RNA-seq showed that the sorted cells shared a majority of the most highly-expressed transcripts, which are known fibroblast markers (Fig. S6C), and enriched for stromal cell signatures and the fibroblast lineage (Fig. S6D). Principal component analysis (PCA) showed notable separation between the wound-, cancer- and normal skin-derived fibroblast groups (Fig. S6E).

A comparison of the significantly up-regulated genes in 5-day wound- and tumor-derived fibroblasts showed that over 25% of these genes were shared between the two conditions (Fig. 5A; Table S7). The shared genes were enriched for pathways related to inflammation, angiogenesis and ECM-related processes, with especially high enrichment for collagen formation and remodeling (Fig. 5B; Table S10).

**Fig. 5.**
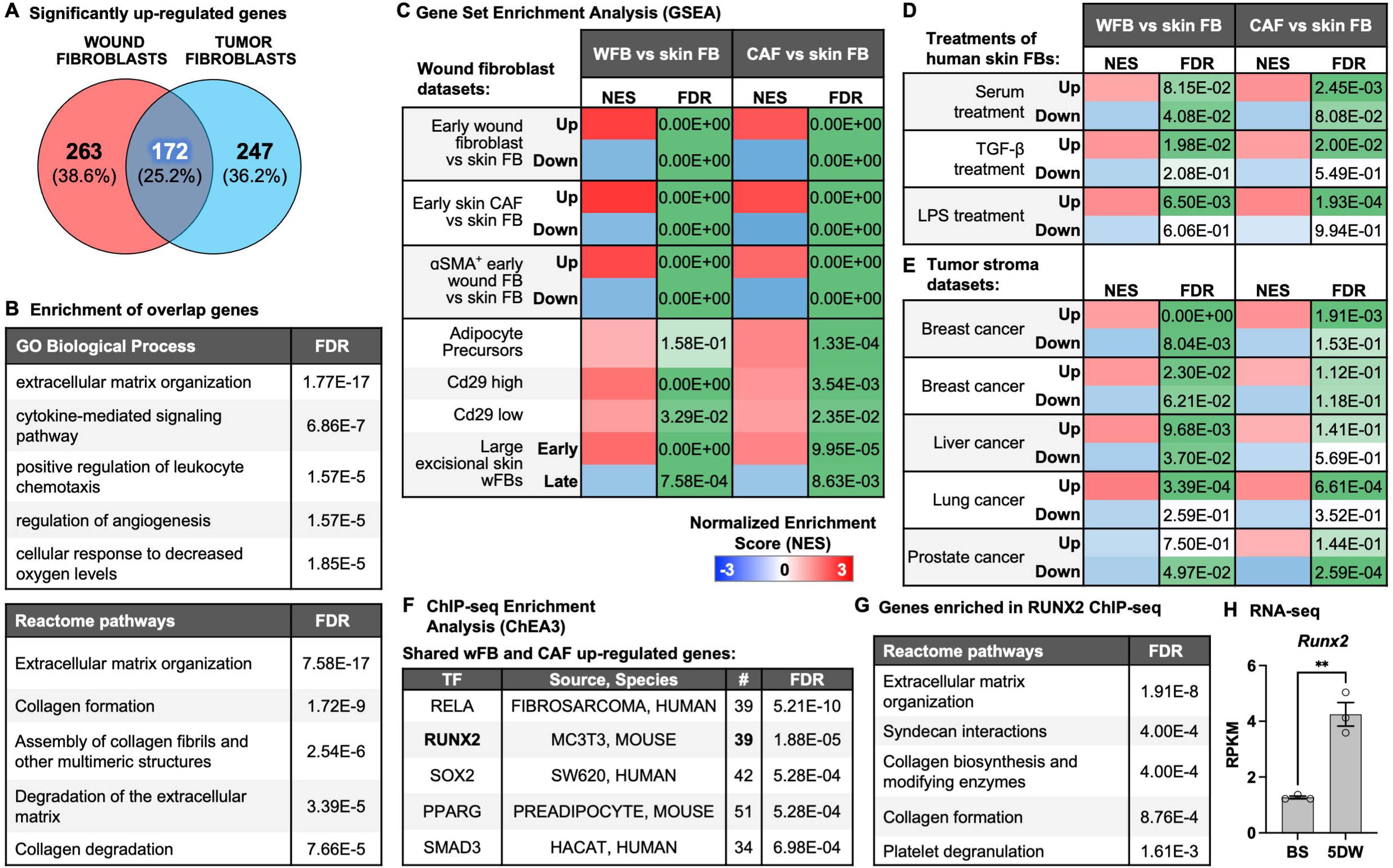
Direct comparisons of early wound fibroblasts and CAFs. (A) Comparison of the significantly up-regulated genes in experimental early wound fibroblasts (wFBs) and skin papilloma-derived CAFs. (B) Functional enrichment analysis of the shared up-regulated genes according to Reactome pathways. (C) Gene set enrichment analysis (GSEA) comparing early wFB and CAF up-regulated genes with published datasets of wFBs, showing normalized enrichment scores (NES) and statistical significance. (D, E) GSEA comparing early wFB and CAF up-regulated genes with published datasets of human skin FBs treated with serum, TGF-β or lipopolysaccharide (LPS) (D) or with published datasets of tumor stroma from across several cancer types (E). (F) ChIP-seq Enrichment Analysis (ChEA3) of shared wound fibroblast (wFB) and skin CAF up-regulated genes, showing number of enriched genes per transcription factor (TF); FDR: false discovery rate. (G) Functional enrichment analysis according to Reactome pathways of shared wFB and skin CAF up-regulated genes, which were also predicted to be regulated by RUNX2 by ChEA3 (F). (H) Runx2 mRNA abundance measured by RNA-seq in fibroblasts derived from intact back skin (BS) and 5-day wounds (5DW) (n=3).

We next performed gene set enrichment analyses (GSEA) to compare our wound- and tumor-derived fibroblasts with other published wound-and cancer-related datasets (Table S1), and found high positive co-enrichment of both wound- and tumor-derived fibroblasts for genes that are overexpressed in early wound (myo)-fibroblasts, including most clusters identified in a scRNA-seq analysis of early wound fibroblasts (Figs. 5C and S6F; Table S11). Leading edge analyses revealed significant numbers of co-enriched genes in at least three datasets (Fig. S6G), many of which were known myofibroblast and CAF markers (Figs. S6G and S6H, bolded genes). Comparison of early wound- and tumor-derived fibroblasts with *in vitro* fibroblast datasets showed dual enrichment of up-regulated genes in serum-, transforming growth factor (TGF)-β- and lipopolysaccharide-treated skin fibroblasts (Fig. 5D). Their comparison with up-regulated genes in tumor stroma also showed high co-enrichment, especially in breast, liver and lung cancer (Fig. 5E). These results suggest that the gene expression signatures of early wound- and tumor-derived fibroblasts are a consequence of the complex stimulatory microenvironments present in both wounds and tumors, which include inflammatory mediators and growth factors [7].

To identify the potential regulators of the genes strongly upregulated in bulk-sorted mouse early wound- and tumor-derived fibroblasts, we performed ChIP-seq Enrichment Analysis (ChEA) and IPA Upstream Regulator analysis on this gene set. Interestingly, RELA (NF-κB subunit) and RUNX2 were identified as the top candidate regulators of the early wound fibroblast/CAF signature (Figs. 5F and S7A). Because the role of RELA in the regulation of CAF markers genes had previously been described [34], we focused our further studies on RUNX2. *RUNX2* itself was a consistent marker for the early, collagen-rich wound fibroblast and CAF subtype in mice (Figs. 2H, 2L, 3I and 3M) and in humans (Figs. 4G, 4H, 4O, and 4P). This was confirmed by pathway analysis of the shared upregulated genes in wound- and tumor-derived fibroblasts that were found to be regulated by RUNX2 in ChEA, which showed strong enrichment for collagen formation related pathways (Fig. 5G). Finally, the RNA-seq of sorted early wound fibroblasts showed significant up-regulation of *Runx2* expression compared to fibroblasts from uninjured skin (Fig. 5H), further pointing to its potential role as a regulator of the early wound fibroblast.

### RUNX2 as a regulator of the early wound CAF subtype

Immunostaining of early excisional skin wounds of Pdgfra-eGFP mice, which express enhanced green fluorescence protein (eGFP) in fibroblast nuclei [35], showed that Runx2 is indeed expressed in wound bed fibroblasts, but not in fibroblasts of uninjured skin (Fig. S7B). This was confirmed by intracellular flow cytometry analysis, which demonstrated a significant increase in the percentage of fibroblasts with high Runx2 expression at days 5 and 7 post-wounding *vs* uninjured skin (Figs. 6A and S7C-S7F for gating strategy). Runx2-positive fibroblasts were also strongly increased in back skin papillomas of HPV8-transgenic mice in comparison to normal skin (Figs. 6B, S7C, S7D, and S7G for gating strategy). Importantly, RUNX2 localized to the stroma of skin BCC, SCC and malignant melanoma according to data from the Human Protein Atlas (Figs. 6C, 6D, 6E, respectively).

**Fig. 6.**
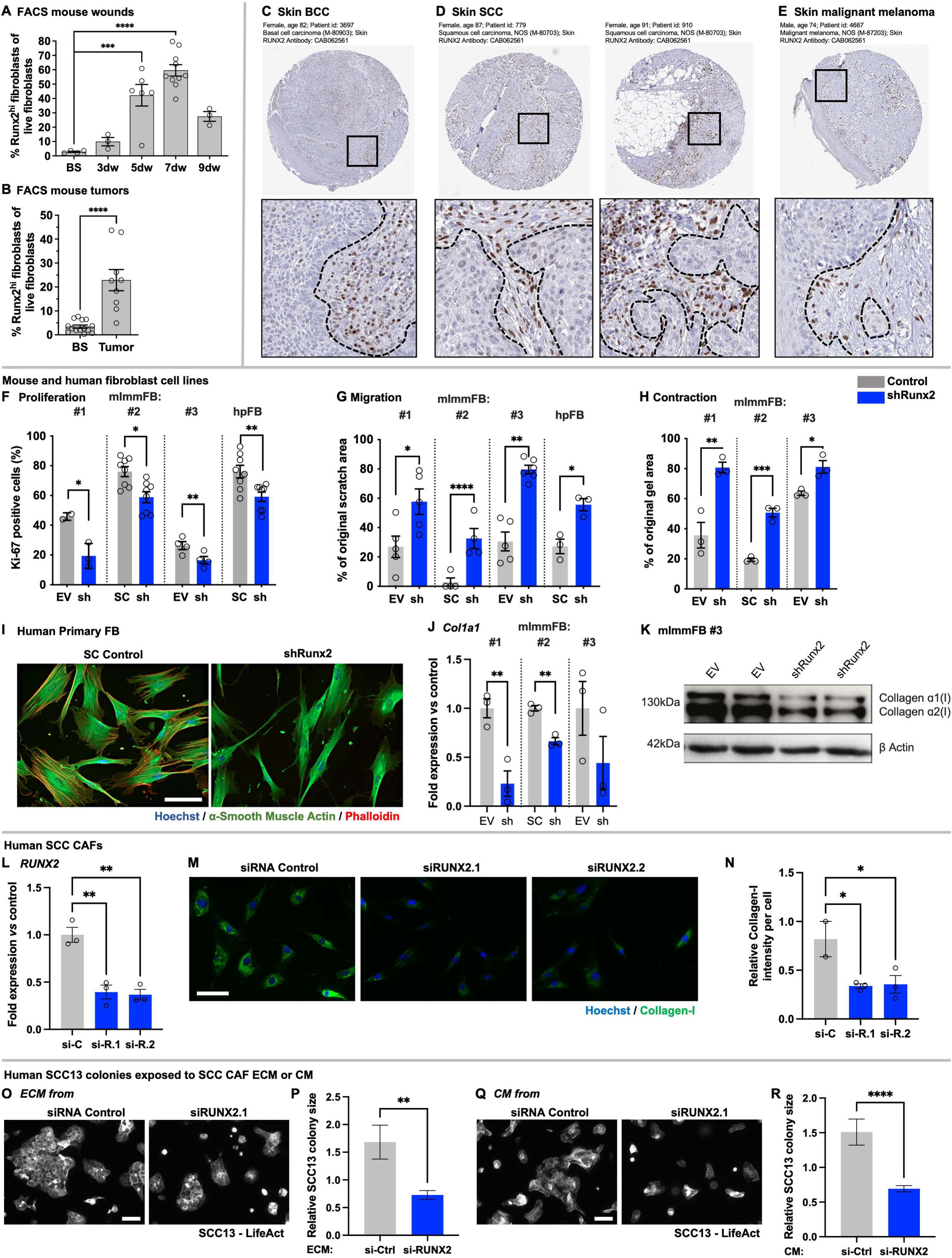
RUNX2 as a regulator of the early CAF subtype. (A) Intracellular flow cytometric analysis for Runx2-highly positive (Runx2^hi^) fibroblasts as a percentage of total live fibroblasts in non-injured back skin (BS) and 3-, 5-, 7-, and 9-day wounds (n=3-10). (B) Intracellular flow cytometric analysis for Runx2^hi^ fibroblasts as a percentage of total live fibroblasts in BS and skin tumors (n=9-14). (C-E) Photomicrographs from Human Protein Atlas showing RUNX2 staining in C) BCC, D) skin SCC, and E) skin malignant melanoma; insets show positive stained cells in respective tumor stroma; black dotted lines indicate tumor/stroma margin. (F) Ki-67 immunofluorescence quantification in mouse immortalized fibroblasts (mImmFB) or human primary fibroblasts (hpFB) transduced with lentiviruses expressing RUNX2 shRNA (shRUNX2) or viruses carrying an empty vector (EV; control fibroblasts) (n=2-9). (G) Quantification of scratched area after 24 h of cell migration in cultures of shRUNX2 or EV mImmFB and hpFB (n=3-6). (H) Quantification of gel area after 20 h in collagen gels with shRUNX2 or EV mImmFB (n=3). (I) Representative immunofluorescence stainings for α-SMA (green) and phalloidin fluorescence (red) in shRUNX2 or EV hpFB; nuclei are counterstained with Hoechst (blue). (J) qRT-PCR analysis of RNA samples from shRUNX2 *vs* EV mImmFB for *Col1a1* relative to *Rps29* (n=3). Expression in one control sample was set to 1. (K) Western blot analysis showing abundance of collagen α1(I) and α2(I) in shRunx2 and EV immFB, cell line #3; β-actin was used as loading control (n=2). (L) qRT-PCR analysis of RNA samples from primary SCC CAFs transfected with RUNX2 or scrambled siRNA for *RUNX2* relative to *RPL27* (n=3). Expression in one control sample was set to 1. (M) Representative immunofluorescence stainings for collagen type I (green) in siRUNX2-transfected primary SCC CAFs compared to scrambled siRNA controls; nuclei are counterstained with Hoechst (blue). (N) Quantification of collagen type I staining in siRUNX2-transfected primary SCC CAFs compared to controls (n=2-3). (O) Representative LifeAct fluorescence of human SCC13 cells cultured on top of decellularized ECM deposited by siRUNX2-transfected primary SCC CAFs compared to scrambled siRNA controls. (P) Quantification of colony sizes of SCC13 cells cultured on ECM derived from siRUNX2-transfected SCC CAFs compared to scrambled siRNA controls (n=122 colonies per group). (Q) Representative LifeAct fluorescence of human SCC13 cells cultured in conditioned medium (CM) derived from siRUNX2-transfected primary SCC CAFs compared to scrambled siRNA controls. (R) Quantification of colony sizes of SCC13 cells cultured in CM derived from siRUNX2-transfected SCC CAFs compared to scrambled siRNA controls (n=173 colonies per group). Scale bars = 100 μm, Bars show mean ± SEM; *p<0.05, **p<0.01, ***p<0.001, ****p<0.0001 (Student’s t-test (B, F-H, J, P, R), one-way ANOVA followed by Bonferroni’s post-tests (A, L, N)).

To analyze the function of Runx2 in skin fibroblasts, we generated immortalized mouse fibroblast cell lines and human primary dermal fibroblasts with stable shRNA-mediated knock-down of Runx2/RUNX2 (Figs. S7H-S7K, Table S12). The knock-down significantly reduced their proliferation and migration (Figures 6F, 6G, S8A and S8B), as well as their collagen gel contraction capability (Figs. 6H and S8C), which correlated with a decrease in α-SMA expression and stress fiber formation (Figs. 6I and S8D). Runx2 knock-down also led to a significant reduction in *Col1a1* gene and collagen type I protein expression in the immortalized fibroblasts (Figs. 6J and 6K, respectively).

To study RUNX2 function in primary human CAFs, we used well-characterized patient-derived skin SCC CAFs [36] and induced transient RUNX2 knock-down with siRNA (Fig. 6L). RUNX2 knock-down CAFs produced significantly less type I collagen (Figs. 6M-6N) and showed reduced expression of CAF markers associated with the production of a tumorigenic matrix, including *INHBA* and *FN1* (Fig. S8E) [33]. To determine a possible role of RUNX2 in the production of a pro-tumorigenic ECM by CAFs, we seeded skin SCC cells (SCC13 cell line) on decellularized ECM deposited by CAFs, which had been transfected with non-target, negative control siRNA or RUNX2 siRNA. Colonies that developed on matrix deposited by RUNX2 knock-down fibroblasts were significantly smaller compared to colonies that developed on matrix from control cells (Figs. 6O and 6P). A similar effect of *RUNX2* knock-down in CAFs was seen when SCC13 cells were exposed to CAF-conditioned medium (Figs. 6Q and 6R).

These results point to an important role of RUNX2 in regulating major early wound- and cancer-associated fibroblast functions. The pro-tumorigenic potential of RUNX2-controlled CAF ECM confirmed the association of RUNX2 with the early CAF/collagen-enriched subtype signatures and also highlights its functional importance.

### ECM imaging of human wounds and tumors reveals spatially distinct collagen and elastin architectures

Besides *RUNX2*, *POSTN* and *MFAP4* were among the top marker genes, which were specifically up-regulated in the early wound collagen-rich CAF subtype (*RUNX2, POSTN*) or in the late wound elastin-rich CAF subtype (*MFAP4*) in mice (Figs. 2H, 2L, 3I and 3M) and humans (Figs. 4G, 4H, 4O, and 4P). We therefore performed global gene co-expression analyses on pan-cancer datasets and identified the genes that were strongly co-expressed with *RUNX2*, *POSTN* or *MFAP4* (Table S13). There was a large overlap between the genes that were co-expressed with *RUNX2* or *POSTN*, and these genes enriched for the “collagen formation” pathway (Fig. 6A, left). In contrast, there was minimal overlap between genes co-expressed with *MFAP4 vs RUNX2*/*POSTN*, and the non-overlapping genes co-expressing with *MFAP4* enriched for “elastin formation” (Fig. 7A).

**Fig. 7.**
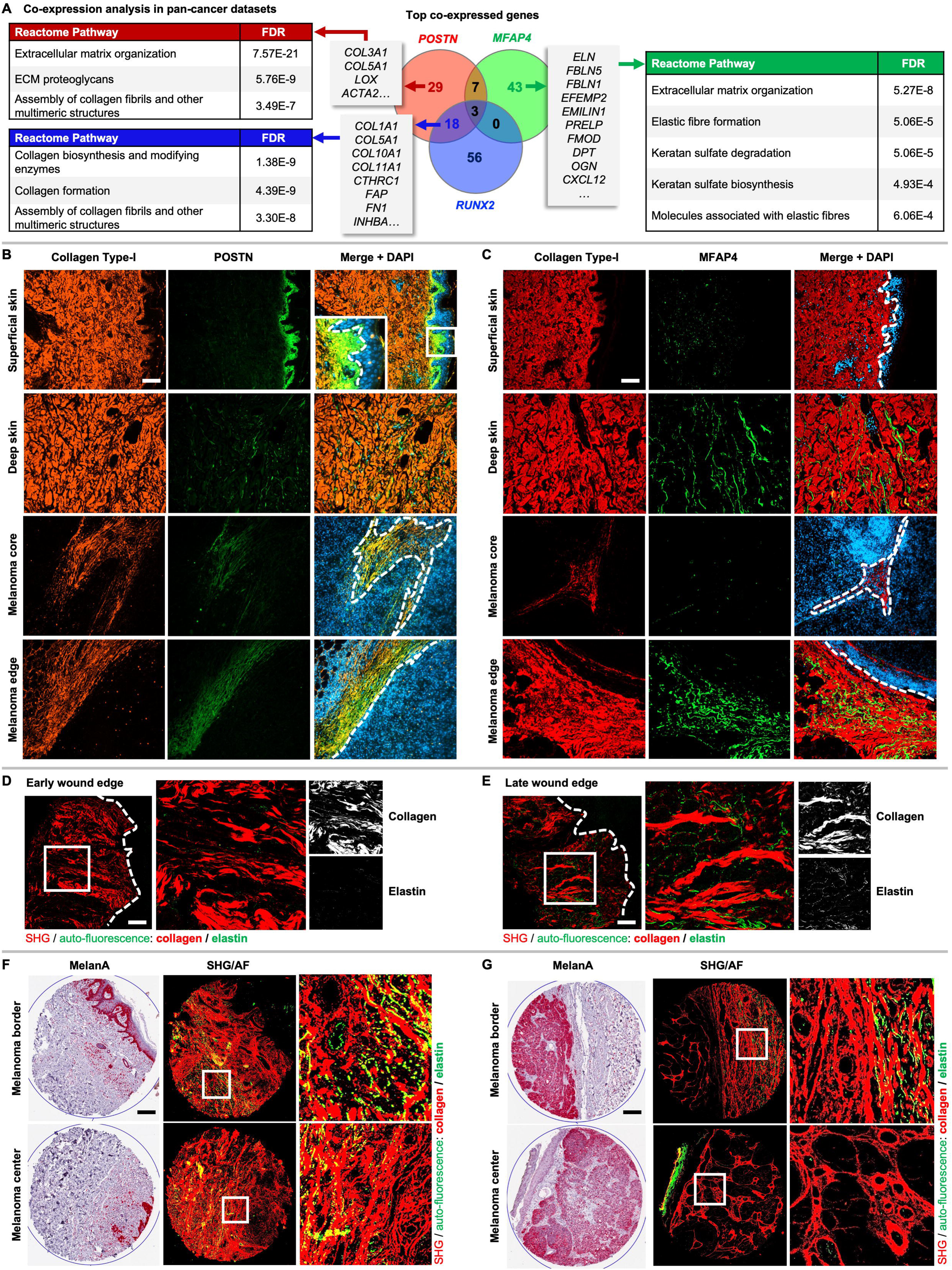
Imaging of matrix proteins in human skin wounds and tumors. (A) Global gene co-expression analysis for *RUNX2*, *POSTN* and *MFAP4* using pan-cancer datasets. A comparison of the top co-expressed genes (center) and functional enrichment analysis using Reactome pathways for genes uniquely or jointly co-expressed with *POSTN* and/or *RUNX2* (left) or uniquely co-expressed with *MFAP4* (right) are shown. (B, C) Representative photomicrographs showing immunofluorescence co-staining for collagen type I (red) and POSTN (B) or MFAP4 (C) (green) in sections of human superficial and deep skin and subcutaneous melanoma core and edge derived from several patients. Nuclei are counterstained with DAPI (blue). Inset in (B) shows characteristic papillary dermis staining of POSTN in healthy skin. (D, E) ECM imaging of the wound edge of early- (D) and late-stage (E) human excisional wounds using second-harmonic generation (SHG) for collagen (red) and auto-fluorescence (AF) for elastin (green). Insets show detailed organization of collagen and elastin fibers, with greater elastin deposition in late-stage wounds. (F, G) Staining of primary melanoma tumor cores for melanA to identify tumor cells (left), and ECM imaging (center, right) by SHG/AF of adjacent sections of less aggressive (F) and more aggressive (G) lesions biopsied at the melanoma border (top) and center (bottom). Insets show detailed organization of collagen and elastin fibers, with highly-aligned collagen fibers close to tumor margins and elastin fibers appearing at longer distances from tumor margins. Scale bars = 200 μm. White dashed lines represent epidermal-dermal junctions (B-E) or tumor margins (B, C).

Immunostaining of sections from normal human skin and primary and metastatic melanoma from several patients for POSTN and MFAP4, combined with co-staining for collagen type I and fibronectin, showed the expected distribution of POSTN in the collagenous papillary dermis with minimal staining in the deep dermis [37] (Fig. 7B), whereas MFAP4 showed fibrillar staining in the superficial and deep reticular dermis, which did not co-localize with collagen or fibronectin (Figs. 7C top, S9A and S9B). Subcutaneous melanoma metastases showed strong co-localization of POSTN with stromal collagen (Fig. 7B, bottom). In contrast, MFAP4 showed minimal signal within tumor stroma, but significant fibrous staining at the tumor periphery and further away in the adjacent non-tumor stroma (Figs. 7C bottom, S9C and S9D). In both cases MFAP4 did not co-localize with collagen or fibronectin.

We then stained serial sections with the collagen-specific Picrosirius Red and analyzed them under brightfield and polarized light conditions. Both POSTN-rich and MFAP4-positive adjacent tumor stroma was collagen-rich, but the collagen architecture in these areas was strikingly different. Notably, while MFAP4-positive stroma had thick, non-aligned collagen fibers with a basket-weave appearance (Fig. S9E), POSTN-rich areas of the tumor stroma showed highly aligned, fibrotic collagen fibers (Fig. S9F).

To image collagen and elastin simultaneously, we utilized multiphoton microscopy with second-harmonic generation (SHG) specific for collagen and auto-fluorescence (AF) signal specific for elastin [38, 39]. Imaging the antibody-stained non-tumorigenic skin adjacent to the tumor lesion demonstrated co-localization of MFAP4 with elastic fibers (Figs. S9G and S9H). We optimized the label-free collagen/elastin ECM imaging protocol for use with rehydrated, unstained sections to obtain unambiguous signals from both ECM types [40, 41]. This allowed for simultaneous and efficient investigation of collagen and elastin architectures as shown for primary melanoma and BCC (Figs. S9I and S9J). We therefore used it to investigate collagen and elastin deposition in human excisional skin wounds (see Experimental Procedures). Late-stage wounds showed considerable elastin deposition next to collagen fibers in areas of the scar tissue, in contrast to early-stage wounds that had minimal elastin signal in proximity to the granulation tissue (Figs. 7D, 7E, S9K and S9L). These results confirmed our transcriptomic meta-analyses of whole wounds and wound fibroblasts, which showed up-regulation of elastin-associated genes in the late-stage wounds.

### Divergent collagen and elastin fiber dynamics may predict primary melanoma outcome

Finally, we used the ECM imaging protocol to visualize the divergent ECM repertoires in primary melanoma using a well-characterized tissue microarray (TMA) (Fig. S10A). The TMA consisted of 33 patient samples, which had paired biopsies from the tumor center and tumor border, and were linked to clinico-histopathological data [42] (see Experimental Procedures) (Table S14). The images allowed for analysis of spatially-resolved features of collagen- and elastin-associated ECM architectures in a wide spectrum of relatively benign to more aggressive primary melanoma lesions (Figs. 7F and 7G, respectively).

Fiber-specific analyses using CT-FIRE/CurveAlign (see Figs. 8A and S10B for descriptive ECM parameter diagrams) revealed certain universal features for collagen and elastin fibers across tumor cores (Fig. S10C; Table S15). Collagen fibers were generally significantly thicker, longer and denser within tumor stroma than elastin fibers (Fig. 8B). We then extracted individual ECM fiber properties and correlated them with one another, demonstrating inter-fiber patterns that differed between fiber types and biopsy locations (Fig. S10D). We used this data to plot the individual fibers’ spatial distributions relative to the tumor margins (Fig. 8C), and found a marked distribution of elastin fibers further away from the tumor margins, whereas collagen fibers localized much closer to the tumor margins. These results are consistent with our analyses of the ST data.

**Fig. 8.**
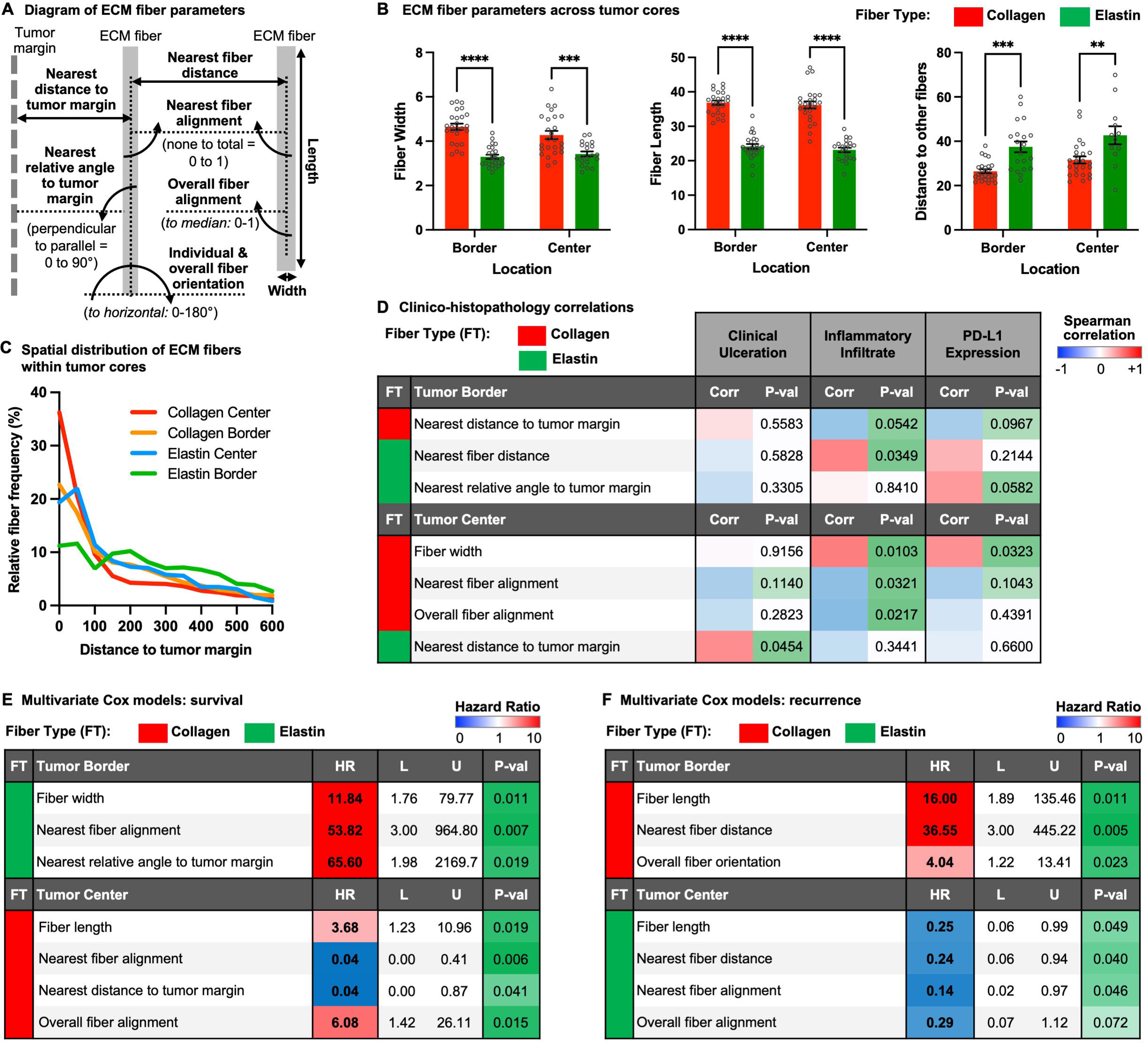
Spatial analysis of collagen and elastic fiber parameters and their prognostic potential in primary melanoma. (A) Diagram describing the measured ECM fiber parameters. (B) Overall parameters that differ between collagen and elastin fibers, including width, length and distance to other fibers, at the tumor border or center. n=21-25. (C) Spatial distribution of ECM fibers within tumor cores plotted as relative fiber frequency (%) over the distance to tumor margins of collagen and elastin fibers at the tumor border or center. (D) Spearman correlation analysis of ECM fiber parameters to melanoma clinico-histopathological features, showing the most significantly correlated collagen (red) and elastin (green) parameters at the tumor border (top) and center (bottom) to presence of clinical ulceration, extent of inflammatory infiltrate and PD-L1 expression. (E, F) Multivariate Cox regression models of ECM fiber parameters significantly impacting patient survival (E) or melanoma recurrence (F), showing the collagen (red) and elastin (green) fiber parameters jointly contributing to statistically-significant hazard ratios (HR) at the tumor border (top) and center (bottom). Bars show mean ± SEM; **p0.01, ***p<0.001, ****p<0.0001 (Two-way ANOVA followed by Bonferroni’s post-tests (B)). FT: fiber type; Corr: correlation; P-val: P-value; HR: hazard ratio; L: lower bound; U: upper bound.

We correlated ECM fiber parameters of the tumor cores with the TMA-linked clinico-histopathological variables, such as ulceration, inflammatory infiltrate, and programmed death ligand 1 (PD-L1) expression, which are important prognostic features of primary melanoma [42, 43] (Fig. 8D; Table S15). At the tumor border, larger collagen distance to the tumor margin correlated negatively with inflammatory infiltrate, while higher elastin nearest fiber distance (e.g. more interspersed matrix) correlated positively with inflammatory infiltrate. In addition, higher elastin fiber angle relative to the tumor margin (e.g. more parallel fibers) correlated positively with PD-L1 expression. At the tumor center, collagen fiber width was correlated positively with inflammatory infiltrate and PD-L1 expression, while nearest fiber alignment and overall fiber alignment were negatively correlated with inflammatory infiltrate. Interestingly, a longer distance of elastin fibers to the tumor margin was positively correlated with the presence of ulceration. Given these correlations between ECM architectures and immunity-related variables, we further showed that staining for the pan-immune cell marker CD45 around the tumor margins of primary melanoma cores coincided with the presence of adjacent elastic fibers (Fig. S10E).

We next performed multivariate Cox modelling using patient survival data (Figs. 8E and S10F; Table S16). At the tumor border, elastin fiber width, nearest fiber alignment, and higher relative angle to the tumor margin (e.g. more parallel orientation) were highly predictive of worse outcome. At the tumor center, higher collagen fiber length, higher overall fiber alignment, lower nearest fiber alignment and lower distance to tumor margin were predictive of worse outcome. These results highlight the importance of spatially-defined ECM architectures for melanoma malignancy, with highly-aligned elastin fibers being prognostic at the tumor border, while at the tumor center these parameters are more prognostic for collagen fibers localizing closer to the tumor margins. Finally, we performed multivariate Cox modelling using melanoma recurrence data (Figs. 8F and S10G; Table S16). At the tumor border, higher collagen fiber length, nearest fiber distance, and overall fiber orientation were highly predictive of tumor recurrence. At the tumor center, higher elastin fiber length, nearest fiber distance, nearest fiber alignment, and overall fiber alignment were predictive of less recurrence. These results suggest that while longer, dispersed collagen fibers at the tumor border may promote melanoma recurrence, these parameters in elastin fibers present at the tumor center may play a protective role.

In summary, our data show that divergent collagen and elastin fiber ECM repertoires are deposited by CAF subtypes with opposing wound healing phase signatures, and are likely determinants of primary melanoma progression, recurrence, and patient outcome.

## DISCUSSION

We identified phase-resolved wound healing signatures in whole wounds and wound fibroblasts and used these phase-specific genes to identify dysregulated wound-associated genes and pathways in whole tumors and CAFs (Figs. S11A and S11B). Many of the most prognostic wound-associated genes enriched for ECM pathways, especially collagen formation, which is consistent with previous studies that identified collagen-related signature genes as highly prognostic for poor outcome across several cancer types [44–50]. Analyses of bulk and single-cell sequenced wound fibroblasts and CAFs identified three main wound-associated CAF subtypes: contractile, collagen-forming, and elastin-forming CAFs. Trajectory analyses suggested a paradigm of differentiation from the early wound-associated contractile and collagen-forming subtypes to the late wound-associated elastin-forming subtypes.

Our data for the early wound-associated subtypes are consistent with the fibrotic and contractile CAF subtypes identified in several cancer types [31, 51–55], especially the myofibroblast CAF subtype (myCAF) [56]. A recent meta-analysis of fibroblasts from perturbed states identified several transcriptomic clusters that were enriched in wounds and tumors, particularly the LRRC15^+^ myofibroblast cluster that shares expression of several marker genes with our early wound-associated CAF subtype, including *LRRC15*, *COL1A1*, *POSTN,* and *RUNX2* [11].

Collagen formation *in vivo* is a complex process involving not only the assembly of major fibrillar collagens into macromolecular alloys, but also less abundant collagens as assembly organizers, intra- and extracellular enzymes acting directly on collagen molecules, non-collagenous glycoproteins as adapters, and a large repertoire of integrins orchestrating the process at the cell membrane [57–59]. The collagen-forming CAF subtype, which was highly correlated with earlier phases of healing, overexpressed many genes encoding proteins involved in this cascade of collagen formation, including subunits of major skin/wound collagens (*COL1A1*, *COL1A2* and *COL3A1*) and of less abundant collagens (*COL5A1* and *COL5A2*), enzymes (*PLOD2*, *LOX* and *LOXL2*), other ECM proteins that bind to collagens and/or regulate collagen deposition (*FN1*, *TNC*, *CTHRC1* and *POSTN*), and integrin subunits (*ITGB1*, *ITGB5*). Additional components may also be upregulated in these cells, but were possibly not detected by scRNA-seq because of the lower coverage compared to bulk RNA-seq. Furthermore, some components may only be regulated at the posttranscriptional level. Taken together, our data suggest that the early wound CAF subtype may be responsible for the fibrotic phenotype of stroma immediately adjacent to tumor margins.

One of the top markers of the early wound-associated collagen-forming CAF subtype was periostin, an extracellular mediator of collagen formation with important functions in healing and carcinogenesis [60, 61]. It interacts with fibrillar collagens, and its absence results in aberrant collagen structure [62–64]. We previously showed that periostin expression and its deposition by wound fibroblasts is associated with exacerbated scar formation, and that activin A (encoded by *INHBA*) directly controls expression of periostin and of some co-expressed ECM genes in both wound fibroblasts and skin CAFs [8, 20, 33]. This was shown to result in the formation of a stiffer ECM in healing wounds [20]. Our scRNA-seq analyses confirmed the relation of *INHBA* with fibrosis, since it was one of the top markers of the collagen-forming CAF subtype. Importantly, high expression levels of *INHBA* are associated with poor prognosis in different types of cancer [33], which correlates with a pro-tumorigenic activity of early-wound CAFs that produce high levels of collagen. Furthermore, we identified RUNX2 as a marker and putative upstream regulator of the early wound-associated CAF subtype, with an important role in the control of fibroblast proliferation, migration, contraction and pro-tumorigenic collagenous ECM deposition. These results are in line with the pro-fibrotic role of RUNX2 in keloid-derived fibroblasts [65], and the promotion of bladder carcinogenesis by RUNX2-expressing CAFs [66]. Our data link these two studies by showing the importance of RUNX2 matrix targets in the pro-tumorigenic effect of this transcription factor.

Although previous studies have not explicitly described elastin-expressing CAFs, elastin fiber formation has been evaluated in some tumor types, and it may have both pro- and anti-tumorigenic functions [67, 68]. Intriguingly, in a fibroblast meta-analysis, the NPNT^+^ non-perturbed cluster strongly resembled our late wound-associated subtype, sharing several marker genes involved in the elastic fiber formation pathway [11]. The elastin-forming CAF subtype was highly correlated with the resolution phase of healing, and one of the top markers of this subtype was MFAP4, a key component of elastic fiber formation [69]. MFAP4 is down-regulated during intrinsic skin aging along with other elastic fiber components [39], and it has a photo-protective role [70]. Previous bioinformatics data predicted better survival of breast cancer patients with high *MFAP4* expression [71], and our data suggest that MFAP4-expressing late wound-associated CAFs may be involved in controlling tumorigenesis by influencing its ECM microenvironment.

Our multi-omics analyses of wound-associated skin CAF subtypes revealed that collagen-related genes were expressed in the inner tumor stroma, while elastin-related genes were expressed in tumor-adjacent stroma. This was confirmed by ECM imaging of primary melanomas. Furthermore, certain collagen and elastic fiber parameters, such as width, length and relative distance and alignment to the tumor margins, were prognostic for survival and recurrence in primary melanoma, and this was dependent on whether the analyzed biopsy was taken from the tumor center or border. Similar location-dependent collagen fiber properties have previously been found to be prognostic in several cancer types, with thick, highly-aligned collagen bundles correlating with increased tumor aggressiveness [72–74].

The correlation of elastic fiber features with melanoma prognosis identified in this study expands on histological studies demonstrating qualitative differences in elastic fiber localization in benign, metastatic and regressing melanoma lesions, in which decreased elastic fiber presence correlated with increased tumor severity [75, 76]. These relationships of divergent ECM repertoires with cancer aggression may involve their effect on immune cells within the TME, wherein stiff and fibrotic collagen-rich ECM prevents T cells from efficiently infiltrating the tumor [77], and therefore limit the efficacy of immunotherapy [45]. In addition, these pro-fibrotic ECM alterations directly affect the cancer cells and also other cells of the TME, which is mediated at least in part by alterations in integrin-mediated signaling [78]. By contrast, elastic fibers are classically linked with higher tissue elasticity, and our results suggest that their presence surrounding the tumor margins correlates with immune cell presence and better clinical outcome. However, these data are preliminary and require validation in future studies.

Our results highlight the tremendous spatial heterogeneity of CAFs and their ECM products, revealing spatially-resolved niches of “wound phases” within the TME. It is within these niches that the classical wound healing pathways become dysregulated and promote pro-tumorigenic phenotypes as seen in “wounds that fail to heal” [6] (Fig. S11C). We demonstrate how the physiological processes of ECM formation and remodeling, normally mediated by wound fibroblasts, are mirrored by CAF subtypes. In the inner tumor stroma and nearby tumor margins, CAFs resembling early wound fibroblasts become hyper-activated by tumor-derived cytokines and growth factors, embody a fibrotic phenotype and deposit a dense, collagen-rich stroma. Just outside of the tumor capsule, CAFs resembling early wound fibroblasts are likely to become hyper-activated biomechanically by the stiff fibrotic matrix and embody a myo-fibroblastic, contractile phenotype. Further away from the tumor margins, CAFs resembling the late wound fibroblasts embody a remodeling phenotype, and they deposit elastic fibers surrounded by a collagen matrix more akin to normal tissue architecture.

Taken together, our results provide insight into the molecular parallels between wound healing and cancer and open new avenues for cancer diagnosis based on expression of wound-regulated genes in fibroblasts and the arrangement of ECM fibers. Finally, the identified wound phase-specific and phase-predictive genes in cancer are promising targets for the development of next-generation cancer therapeutics.

## EXPERIMENTAL PROCEDURES

### Whole wound and wound fibroblast time-course transcriptomes

For the analysis of whole wound (WW) time-course data, datasets from two different wound healing models were used: 1) RNA-seq data from 3-mm diameter excisional, full-thickness skin wounds comprising the time-points 0-day (non-injured back skin), 1-day, 4-day, 8-day and 14-day post-injury; and 2) microarray data from 1-mm diameter excisional, full-thickness skin wounds comprising the time-points 0-day, 12 h, 24h, 3-day, 5-day, 7-day and 10-day post-injury (Table S1). For the analysis of wound fibroblast (WFB) time-course data, two datasets were combined: 1) bulk WFB RNA-seq data from normal back skin and 5-mm diameter excisional, full-thickness wounds at 5-day post-injury, and 2) bulk WFB RNA-seq data from 2.25 cm^2^ rectangular excisional, full-thickness wounds comprising the time-points 12-day, 15-day, 21-day and 26-day post-injury (Table S1). RNA-seq data was processed using CLC Genomics Workbench version 10 (Qiagen; Hilden, Germany). Pre-processed and annotated microarray data was downloaded from GEO. The WFB datasets were combined, and batch correction was performed using the *ComBat* R method implemented in the sva R package (version 3.20.0). Batch correction was assessed by means of PCA. Gene expression in WW and WFB was normalized for each time-point relative to the average expression of the gene across all samples, and subsequent analyses were all based on log2 fold-changes. The Spearman correlation matrices comparing WW and WFB on the transcriptome level or according to wound phase-specific genes were generated using the corrplot R package (version 0.84).

### Identification of phase-specific genes via new Biopeak R package

Impulse-like gene expression changes over the time-course of wound healing were identified by means of a novel peak detection algorithm, which we published as a new R package called Biopeak (https://CRAN.R-project.org/package=Biopeak). This peak detection algorithm features high flexibility in respect to the input dataset, and any time course dataset monitoring a continuous variable can be treated as a signal and processed efficiently by the model. On a high level, the algorithm treats the gene expression of each gene as a signal that propagates along the time axis of wound healing and aims to find the peak of the signal. This is achieved by finding all local maxima (peaks) first and subsequently ordering them by their magnitude before applying a range of filtering criteria, to select only the most prominent peaks. A local maximum is found if the requirement in (1) is satisfied, where y(t) represents the gene expression as a function of the series t (e.g. time or temperature condition).

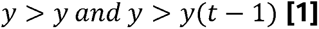

*Filtering Criteria:* Peaks with a magnitude lower than the baseline expression, which is calculated as the mean across all time-points (defined as μ_exp_), are discarded (2).

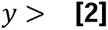

Genes that feature an activation of small magnitude can be excluded. The activation threshold is defined as shown in (3). The higher the standard deviation is relative to the mean expression, the stronger are the activation pulses.

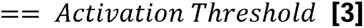

If more than one peak for a specific gene passes this filter criteria, the largest of those peaks is selected by first ordering all peaks b by magnitude and subsequently selecting genes for which the largest peak is at least a certain percentage higher than the second highest peak (peak prominence) (4).

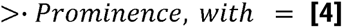

Generally, any series dataset monitoring a continuous variable can be treated as a signal and, thus, can be processed by the model.

Since the goal was to identify gene expression changes that are unique for a specific phase of healing, only genes with a single large peak were considered. In order to select such phase-specific genes, a set of filters was defined: 1) peaks with a magnitude lower than the baseline expression, which is calculated as the mean across all time-points, were discarded; 2) genes that featured an activation of small magnitude were excluded from the analysis, with higher standard deviation relative to mean expression requiring stronger activation pulses, resulting in an activation threshold of 0.2; 3) multiple peaks per gene that satisfied criteria (1) and (2) were ordered by magnitude, with the largest of those peaks selected, but only if the peak was at least 30% higher than the second highest peak; 4) sustained peak activation in the vicinity of the largest peak was assessed by selecting proximal time points that showed an activation of more than 60% of the largest peak. These parameters were optimized for our datasets based on biological reasoning. In general, the tool showed high robustness towards parameter setting changes due to the abundance of well-pronounced peaks.

In whole wounds, this led to the identification of early (Inflammatory phase), mid (Proliferative phase) and late (Resolution phase) phase-specific genes (Table S2). In wound fibroblasts, this led to the identification of Inflammatory and Resolution phase-specific genes (Table S6). Functional enrichment analyses of phase-specific genes were performed using EnrichR [79], Ingenuity Pathway Analysis (Qiagen), and the Cytoscape (version 3.8.2) plugin ClueGO (version 2.5.8) that allowed for the determination and visualization of the relative enrichment of pathways across wound phases [80] (Table S2). Gene sets were compared to one another using Venny 2.1 (https://bioinfogp.cnb.csic.es/tools/venny/).

### Comparison to human whole wound and skin graft transcriptomes

For comparisons to published whole human wound and skin graft transcriptomes, the relevant datasets were downloaded and processed for analysis (Table S1). For the human wound healing RNA-seq dataset, the pre-processed data was downloaded and the genes were filtered for normalized RPKM values >1 and those, whose log fold change compared to the respective intact skin or oral mucosa was >2. Venny 2.1 was used to compare the up-regulated genes between the 3- and 6-day post-injury time points, and the final comparisons were made to phase-specific genes based on these time-dependent genes (Fig S1E; Table S3). For the human split-thickness skin graft microarray dataset, the pre-processed data was downloaded from Geo2R, and the genes were filtered for those whose log fold change compared to respective intact skin was >1.5 and FDR<0.05. Venny 2.1 was used to compare the up-regulated genes between the 3-, 7-, 14- and 21-day post-injury time-points, and the final comparisons were made to phase-specific genes based on the time-dependent genes that only had overlap with the next adjacent time-point, with the genes at 3-, 3/7- and 7-day post-injury classified as “Inflammatory-Proliferative” and the genes at 14-, 14/21-, and 21-day post-injury classified as “Proliferative-Resolution” (Fig S1F; Table S3). Comparisons of both human datasets to healing phase-specific genes were performed using Venny 2.1; the percentages of phase-specific genes shared with the human time-dependent genes were calculated and normalized according to the size of the respective human gene sets.

### Whole tumor transcriptomes

Normalized count data from TCGA and GTEX project databases, which was obtained from the UCSC Xena Browser [81], was filtered for matching tissues of origin, primary site tumors and non-blood and non-lymphatic tissues; we excluded TARGET data, and data from the following tissue sites: “Bone Marrow”, “Blood”, “Blood Vessel”, “Lymphatic tissue”, “White blood cell” (Table S1). TCGA tumor samples were normalized by the average gene expression across all corresponding tissue samples in GTEX, i.e. fold-change relative to GTEX expression. Genes with an average expression of zero in GTEX were given an arbitrarily high fold change of 100,000 to avoid division errors. Next, log2 transformation of the GTEX-normalized data was performed. Since the log of zero is not defined, genes featuring zero expression in TCGA were given a pseudo count of 1 resulting in zero log2 fold-change.

For comparison to published whole Actinic Keratosis (AK), SCC, and BCC transcriptomes, the relevant datasets were downloaded and processed for analysis (Table S1). These data were similarly normalized by the average gene expression across corresponding normal skin samples, and log2 transformation of the skin-normalized data was performed. All normalized expression matrices were then filtered to include only the human orthologs of healing phase-specific genes.

### Wound score calculation in tumor samples

Whole wound samples were classified into four groups (phases): Intact (unwounded), Inflammatory (early), Proliferative (mid) and Resolution (late). These classifications were based on the samples’ clustering according to Spearman correlation matrices. The following samples were grouped together, for Intact: NS_3mm and NS_1mm; for Inflammatory: 1D_3mm, 4D_3mm, 12h_1mm, 24h_1mm; for Proliferative: 8D_3mm, 3D_1mm, 5D_1mm; for Resolution: 14D_3mm, 7D_1mm, 10D_1mm. Next, the Spearman correlation coefficient was calculated for all TCGA patient samples of a given tissue according to their normalized expression and the individual wound samples, using shared phase-specific genes between the two datasets. In order to derive the final wound score, the correlation coefficients between the wound samples and the individual TCGA samples were aggregated by taking the average correlation for each phase of healing. In order to maximize the difference between the scores for the three main phases of healing (Inflammatory, Proliferative and Resolution), a differential wound score was derived for each healing phase using the following formulae: differential Inflammatory score: (Inflammatory score + 1) – (Resolution score + 1) – (Proliferative score + 1); differential Proliferative score: (Proliferative score + 1) – (Resolution score + 1) – (Inflammatory score + 1); differential Resolution score: (Resolution score + 1) – (Inflammatory score + 1) – (Proliferative score + 1). By adding 1 to each score, the negative sign of each negative correlation coefficient was removed, allowing the calculation of score differentials by simple subtraction. A tumor sample could therefore be classified into one distinct phase of healing by comparing the differential wound scores. For comparison to normalized AK, SCC, and BCC transcriptomes, the Spearman correlation coefficient was calculated for all samples and the individual wound samples, using shared phase-specific genes between the two datasets. Mean wound scores across tumor types and individual wound scores for tumors of the skin, breast and lung were plotted using GraphPad Prism (version 9.1.2) (GraphPad Software, LLC, San Diego, CA).

### Robustness analyses for calculation of wound scores via Spearman correlation

In order to verify the robustness of the Spearman correlation coefficient used for the determination of the wound-score, two methods were developed. First, the Spearman correlation was compared to the pair-wise Pearson correlation between wound and TCGA samples for each cancer-type and for the genes selected by the peak detection algorithm. We plotted the Spearman correlation coefficients against the Pearson correlation coefficients and determined their consensus by means of a linear regression model. The adjusted R-Squared was then used to evaluate the goodness of the fit. The pair-wise comparison robustness analyses with their respective R-Squared values for all TCGA cancer types can be found in Table S17. For skin cancer, an adjusted R-Squared value of 0.953 was calculated, indicating a high congruence between the two correlation metrics and, thus, a high methodological robustness. Since the rank-order-based Spearman correlation captures monotonic relationships, it is generally considered a more conservative method than the Pearson correlation coefficient, which is only considering linear relationships between two variables. Therefore, we conclude that for skin cancers the correlation between wound and TCGA samples are to a great extent linear and, thus, the Spearman correlation is highly robust.

Second, we conducted an additional robustness test by removing individual genes from the peak detection gene-set and re-calculating the Spearman correlation, the base of the wound-score (leave-one-out (LOO) analysis). The peak detection gene-set consisted of 1660 individual genes shared between wound and tumor samples. For each removed gene, the Spearman correlation between all wound and TCGA samples was calculated resulting in 1660 correlation matrices. For each TCGA sample and wound sample, the standard deviation of the correlation across all possible gene-sets was calculated, resulting in a tensor with dimensions: (number of wound-samples x number of LOO gene-sets x number of TCGA samples). Finally, the distribution of the tensor was assessed by plotting a histogram for each wound sample encompassing all TCGA samples with all LOO gene-sets. The LOO robustness analyses with their respective standard deviation values for skin and uterus TCGA cancer types can be found in Table S18. For skin cancer, the standard deviation of the Spearman correlation upon removing genes individually across all wound samples was below 0.0007 and, thus, very low. The removal of an individual gene from a gene-set comprising of 1660 genes and aggregated by a rather robust rank-based correlation metric such as the Spearman correlation has only a small effect on the overall scores. These findings indicate a high robustness of the Spearman correlation as the wound-score metric.

### Identification of tumor phase-predictive genes using machine learning

In order to identify the genes in the TCGA tumor samples that were most predictive for a high wound score of a given phase of healing, a classification problem was formulated. Each tumor sample of a specific tissue was sorted by its differential wound score. Next, the top decile (10%) of samples with the highest Inflammatory, Proliferative or Resolution differential scores were labelled and used for classification. The selection of the top decile was chosen to maximize the predictive power by training the classification model on only the samples that classify unambiguously to a specific phase of healing, while also avoiding class imbalance problems. The latter are characterized by training a predictive model with two classes, (here: phases of healing) with a highly imbalanced number of observations between the two groups [82]. Our selection process fixes the number of observations in both categories and, thus, avoids that problem.

In order to isolate the expression signals of each healing phase, pairwise comparisons were performed: Inflammatory-to-Resolution, Inflammatory-to-Proliferative, Proliferative-to-Resolution. The data-frame consisting of all genes identified by the peak detection algorithm and the top scoring samples labelled by their representative phase of healing was split into training and testing datasets (75% training, 25% testing). Next, the data was scaled and centered, and linear discriminant analysis with repeated cross-validation (3 folds and 3 repeats) was performed. Indeed, K-fold cross validation is known to provide reliable accuracy estimates [83].

Both training and testing performance accuracy were monitored via Cohen’s Kappa and confusion matrix. Overall, the test-set accuracy was high with scores above 0.95 for most cancer types, with the exception of colon, esophagus and lung for the comparison Inflammatory-to-Proliferative (accuracies above 0.8). Moreover, testis and pancreas Proliferative-to-Resolution comparisons featured a test-set accuracies of 0.63 and 0.75, respectively. For uterine cancer, the test-set comprised only 2 samples and was therefore inconclusive. Specifically, skin Inflammatory-to-Resolution and Proliferative-to-Resolution showed rather high test-set accuracies of 0.96 and 1 (with 22 test-set samples and 72 training-set samples), respectively. A complete list of all models and their associated performance metrices can be found in Table S19.

The choice of LDA for the machine learning model was based on its close relationship with principal component analysis and, thus, its inherent ability to handle high dimensional datasets, i.e. high number of columns/inputs [84]. LDA aims to project the data into a new feature space that maximizes the class separability. Notably, while the application of LDA requires the classes to be linearly separable, it tends to overfit much less than other, non-linear classifiers [85].

The feature importance of the binary classification was calculated by means of the *varImp* function (R library Caret using ROC Curve analysis, version 6.0-86), representing the most phase-predictive genes. Phase-predictive genes for each tumor tissue were aggregated using the Borda count method to identify tissue-independent predictors. For each tumor tissue and phase, the top third ranking genes with the highest feature importance were selected. Next, the ordinary Borda count (R library votesys, version 0.1.1) was applied to all candidate genes resulting in a ranked list of genes. The gene with the lowest Borda count represented the highest ranked gene across all tumor tissues for a given phase. In order to combine the phase-predictive gene lists for the three healing phases into one single list, each list was filtered. The 60 percentile for Inflammatory, 55 percentile for Proliferative and 50 percentile for Resolution was selected, representing the observed distribution of phase-specific genes after the peak detection algorithm. This led to the identification of tumor Inflammatory, Proliferative and Resolution phase-predictive genes (Table S4). Wound scores of TCGA tumor samples were then recalculated using these phase-predictive genes. The prognostic potential of the wound scores was determined by first uploading the tumor wound score meta-data into the UCSC Xena Browser and then plotting Kaplan-Meier curves according to median wound score stratification of patients in primary and metastatic melanoma [81]. Functional enrichment analysis of phase-predictive genes was performed using the Cytoscape (version 3.8.2) plugin ClueGO (version 2.5.8) that allowed for the determination and visualization of the relative enrichment of pathways across wound phases [80] (Table S4).

### Identification of prognostic tumor phase-predictive genes

Tumor phase-predictive genes of the Proliferative and Resolution phases were analyzed for their prognostic potential using the PRECOG database of normalized survival scores across many different cancer types in TCGA and other compiled cancer datasets [21]. Potentially prognostic genes were considered if their TCGA or PRECOG metaZ scores were less than −1 (for better outcome) or more than 1 (for worse outcome) (Table S5). Functional enrichment analysis of the prognostic Proliferative and Resolution phase-predictive genes was performed using the Cytoscape (version 3.8.2) plugin ClueGO (version 2.5.8) that allowed for the determination and visualization of the relative enrichment of pathways across the TCGA and PRECOG databases [80]. Gene-gene interaction network with functional enrichment of the Proliferative and Resolution phase-predictive prognostic genes was generated using the StringApp plugin for Cytoscape [86] by inputting the shared worse outcome gene list and manually color-labeling enrichments in the GO and Reactome pathway databases. Cell type enrichment was determined using ARCHS4 Cell Lines Enrichment tab in EnrichR and by ImmGen MyGeneSet RNA-seq (http://rstats.immgen.org/MyGeneSet_New/index.html). Gene sets were compared to one another using Venny 2.1. For the determination of genes associated with increased severity in NMSC, the 200 top normalized up-regulated genes in SCC *vs* AK and treatment-resistant BCC *vs* treatment-sensitive BCC were determined, compared using Venny 2.1, and functional enrichment of the overlapped genes was performed via EnrichR and the Reactome pathway tab.

### Animals

PDGFRα-eGFP transgenic mice [35] and transgenic mice expressing the HPV8 oncogenes (F1 progeny of CD 1 and C57BL/6 background mice) [87, 88] were used. HPV8 transgenic mice were monitored regularly for development of spontaneous back skin papillomas. We used the PDGFRα eGFP transgenic mice and C57BL/6 wild-type mice to analyze wound healing using the excisional wound model, wherein female mice (9–12 weeks old) were anaesthetized by ketamine/xylazine intraperitoneal injection. Their back skin was shaved and cleaned with 70% ethanol, and four full thickness excisional wounds (5□mm diameter) were generated on either side of the back midline. The mice were housed under specific pathogen-free conditions and received food and water *ad libitum*. Mouse maintenance and all animal experiments had been approved by the veterinary authorities of the Swiss Canton of Zurich (Kantonales Veterinäramt Zürich).

### Genotyping

Animals were genotyped using DNA extracted from toe or ear biopsies. DNA was amplified by PCR using KAPA2G FAST Genotyping Mix (Kapa Biosystems, Wilmington, MA) using primers listed in Table 1, and analyzed by agarose gel electrophoresis followed by ethidium bromide staining of the gels.

**Table 1.**
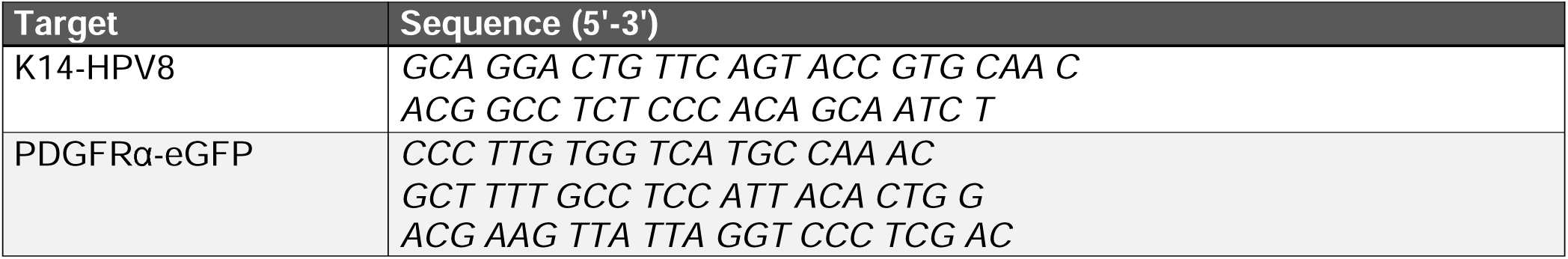
Primers used for genotyping

### Intracellular flow cytometry analysis

Wound samples collected 3, 5, 7, and 9 days post-injury and back skin tumors were harvested and processed for flow cytometry according to the protocol described in [89]. Intracellular staining was performed using the Transcription Factor Phospho Buffer Set (BD Pharmingen, San Diego, CA) according to the manufacturer’s protocol. The staining panel used is listed in Table 2. Compensation of fluorescence emission was performed using compensation beads (BD Biosciences, Allschwil, Switzerland). Stained cells were analyzed using a BD LSRII Fortessa equipped with FACSDiva software (Version 6) (BD Pharmingen). Samples were acquired using Fortessa’s HTS plate reader option at an event rate below 20,000 events/s. Isotype and fluorescence minus one (FMO) control samples were used for staining and gating verification. Compensation adjustment, gating, and data analysis were performed using FlowJo software (Version X, Tree Star, Inc., Ashland, OR). The gating strategies for analysis of Runx2^+^ fibroblasts in wounds and tumors are diagrammed in Fig. S7C-G. Data was combined from several experiments to generate the final figures.

**Table 2.**
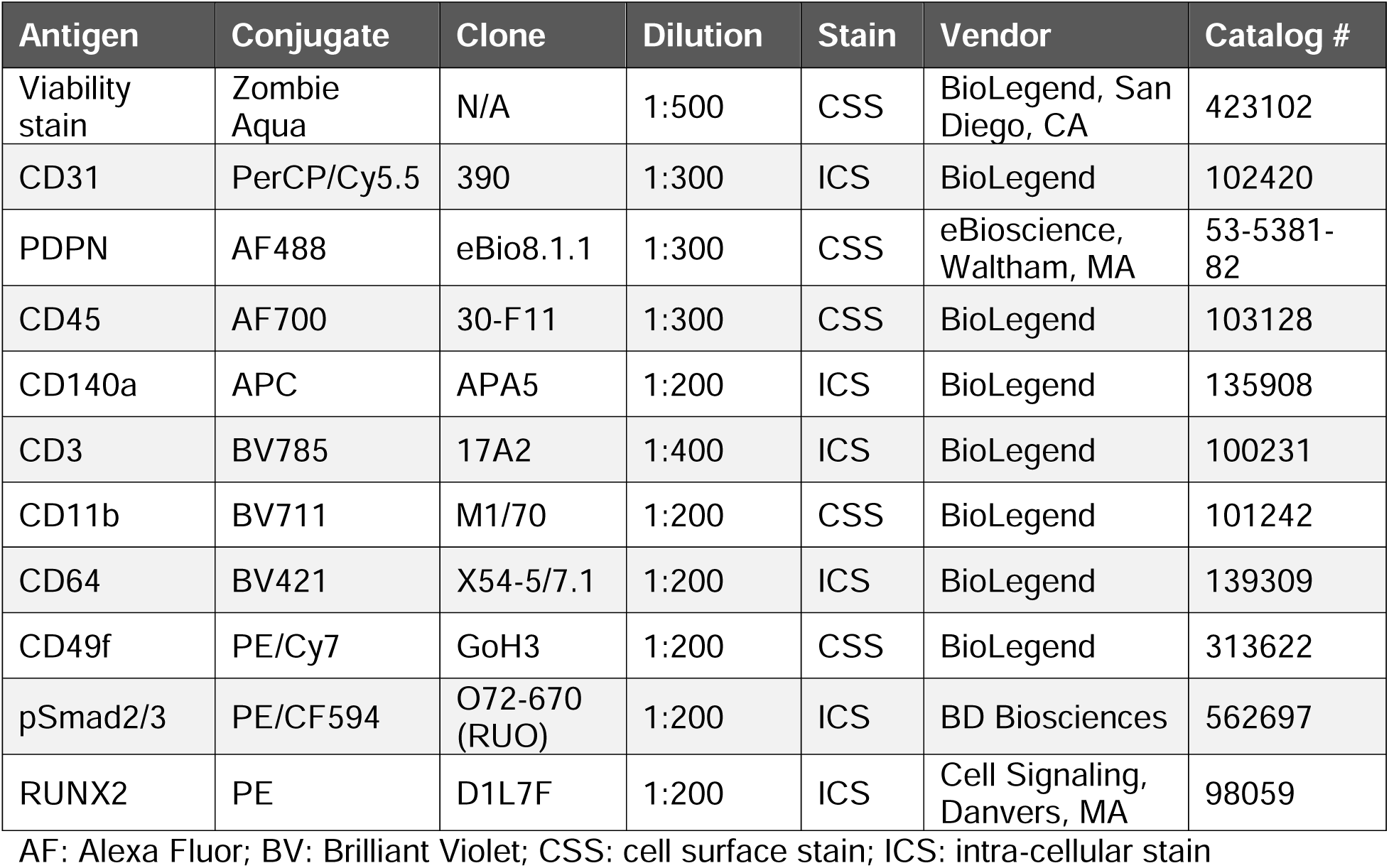
Dyes and antibodies used for flow cytometry

| Antigen | Conjugate | Clone | Dilution | Stain | Vendor | Catalog # |
| --- | --- | --- | --- | --- | --- | --- |
| Viability stain | Zombie Aqua | N/A | 1:500 | CSS | BioLegend, San Diego, CA | 423102 |
| CD31 | PerCP/Cy5.5 | 390 | 1:300 | ICS | BioLegend | 102420 |
| PDPN | AF488 | eBio8.1.1 | 1:300 | CSS | eBioscience, Waltham, MA | 53-5381-82 |
| CD45 | AF700 | 30-F11 | 1:300 | CSS | BioLegend | 103128 |
| CD140a | APC | APA5 | 1:200 | ICS | BioLegend | 135908 |
| CD3 | BV785 | 17A2 | 1:400 | ICS | BioLegend | 100231 |
| CD11b | BV711 | M1/70 | 1:200 | CSS | BioLegend | 101242 |
| CD64 | BV421 | X54-5/7.1 | 1:200 | ICS | BioLegend | 139309 |
| CD49f | PE/Cy7 | GoH3 | 1:200 | CSS | BioLegend | 313622 |
| pSmad2/3 | PE/CF594 | O72-670 (RUO) | 1:200 | ICS | BD Biosciences | 562697 |
| RUNX2 | PE | D1L7F | 1:200 | ICS | Cell Signaling, Danvers, MA | 98059 |

AF: Alexa Fluor; BV: Brilliant Violet; CSS: cell surface stain; ICS: intra-cellular stain

### Fluorescence-activated cell sorting (FACS) and RNA-seq of mouse skin tumor-associated fibroblasts

Papillomas that formed spontaneously the back skin of HPV8-transgenic animals were harvested, along with the normal back skin (NS) of littermates. Two tumors of 70- and 78-week-old animals were analyzed, along with six NS samples of 68-, 69- and 70-week-old animals. The tissues were processed for FACS of fibroblasts as described [20]. For cell analysis and sorting, fibroblasts were defined as CD140a^+^ CD45^−^ CD11b^−^ F4/80^−^ live cells. From tumors, 100,000-110,000 fibroblasts were isolated per sample; from NS, 20,000-120,000 fibroblasts were isolated per sample; several NS samples were pooled to achieve adequate cell numbers. RNA was isolated via RNeasy Micro Kit (Qiagen), and RNA quality was analyzed using a TapeStation (Agilent Technologies, Santa Clara, CA). RNA samples with RIN□>□7.0 were adjusted to 3 ng/μl and were subjected to the RNA-seq protocol via poly-A enrichment, True-Seq library preparation, and single-end 100□bp sequencing on an Illumina HiSeq 2500 v3 instrument at the Functional Genomics Center Zurich (FGCZ). Sequence alignment to the mouse reference genome (build GRCm38), quality control, and initial bioinformatics analyses were performed in the CLC Genomics Workbench (version 12) (Qiagen). All samples generated approximately 16 million reads, showed high (98.5-98.8%) mapping rates to the mouse reference genome, high (96.0-97.1%) mapping rates to genes, and low (0.15–0.39%) rRNA mapping rates. Numbers of RNA transcripts were determined on the gene level and normalized to reads per kilobase of transcript, per million mapped reads (RPKM).

The tumor fibroblast (CAF) RNA-seq data was combined with that obtained from fibroblasts isolated from 5-day wounds (wFB) and age-matched normal skin from mice of the same genetic background and using previously described FACS protocols [20] (Table S1). The datasets were analyzed together in CLC Genomics Workbench (version 12) (Qiagen), and PCA was performed to evaluate transcriptome grouping of all samples. Genes were ranked according to absolute expression in all fibroblasts, and functional enrichment of the shared genes among 100 top-expressed genes was determined using ARCHS4 Cell Lines Enrichment tab in EnrichR and by ImmGen MyGeneSet. Differential expression analysis was performed across all group comparisons using the edgeR exact test as implemented in the CLC Genomics Workbench (Qiagen), and differentially expressed genes (DEGs) for wound fibroblasts (5DW *vs* respective NS) and CAF *vs* respective NS were determined and compared at the statistical significance level of FDR□<□0.05 (Table S10). Venny 2.1 was used to compare the statistically significant (FDR□<□0.05 DEGs across 5dw and Tumor *vs* respective NS comparisons. Shared up-regulated DEGs were subjected to functional enrichment analysis using Gene Ontology (GO) biological processes and Reactome Pathways in EnrichR. Analysis of upstream regulators of the WFB and CAF up-regulated genes was performed using Chip-seq Enrichment Analysis (ChEA3) [90] and the Upstream Regulators tab within IPA (Qiagen).

### Gene Set Enrichment Analysis (GSEA)

Sets of significantly up- and down-regulated genes were generated by mining of various relevant databases, including GEO for published transcriptomic data of wound-derived (myo)fibroblasts, treated skin fibroblasts, tumor stroma, and published scRNA-seq data (Table S1). Original gene sets were uploaded to GSEA (version 4.1.0) and filtered to those mapped by gene symbol and present in the tested datasets. The gene sets were tested against the wound fibroblast signature (wFB *v*s NS) and the cancer fibroblast signature (CAF *vs* NS). GSEA results were organized in tables, and the normalized enrichment scores (NES) and FDR values were color coded for visualization. Leading Edge analyses were extracted to identify commonly enriched genes between gene sets. The previously published datasets and samples analyzed, original and filtered gene sets, GSEA settings used for each analysis, ranked gene lists, and original GSEA output data, including leading edge analyses for all experiments, are provided in Table S11.

### Primary cells and cell lines

Mouse primary cells and immortalized fibroblast lines were established as previously described [91]. Immortalized skin fibroblasts were initially isolated from PDGFRα-eGFP transgenic mice and immortalized via serial passaging.

Primary foreskin fibroblasts and CAFs were established from human foreskin and skin SCC biopsies, respectively. CAFs were directly isolated from skin SCC biopsies alongside their paired normal skin fibroblasts as described previously, with some modifications [92]. Briefly, skin samples were digested with trypsin, followed by digestion with collagenase. Cells were cultivated with J2 feeder cells. Fibroblasts and CAFs were separated from SCC cells upon short incubation with trypsin and then cultivated without feeder cells in Dulbecco’s modified Eagle’s medium (DMEM), 10% fetal bovine serum (FBS), 1% penicillin/streptomycin (P/S). The CAF phenotype was confirmed by expression of skin CAF markers and enhanced mitogenic and motogenic properties of these cells [93], and fibroblast cultures isolated from SCCs that did not have CAF properties were excluded.

Human primary foreskin fibroblasts were kindly provided by Dr. Hans-Dietmar Beer, University of Zurich, Switzerland. The foreskin had been collected with informed written consent of the parents in the context of the Biobank project and approved by the local and cantonal Research Ethics Committees. The SCC13 cells (human cutaneous squamous cell carcinoma cell line; [94]) were kindly provided by Dr. Petra Boukamp, Leibniz Institute for Environmental Research, Düsseldorf, Germany. All cells were cultured in DMEM supplemented with 10% FCS and P/S and regularly tested for mycoplasma contamination.

### Lentiviral production, transduction and stable transfection of fibroblasts with shRNA

Immortalized mouse and primary human dermal fibroblasts were infected with lentiviral shRNA vectors designed by The RNAi Consortium (TRC), as described previously [33]. Five shRNAs, each directed against mouse and human Runx2/RUNX2 in the pLKO.1 lentiviral vector, were tested (Table 3). Immortalized mouse fibroblasts were infected with TRCN0000095589-93, and primary human dermal fibroblasts were infected with TRCN0000013653-57. Briefly, HEK293T cells at 30% confluency were transfected overnight in DMEM/10% FBS without P/S using jetPEI Transfection Reagent (Polyplus-Transfection SA, Illkirch, Graffenstaden, France; #101-40N). For each 10 cm dish, 5 μg DNA was incubated with1.75 μg pMD2.G and 3.25 μg pCMV-dR8.91 (Addgene, Watertown, MA; #12259). On the next day, the medium was replaced by DMEM/10%FBS/P/S. Cells were cultured for at least 48 h, and the supernatant was filtered through a Filtropur S 0.2-μm filter (Sarstedt, Nürnbrecht, Germany) and stored at −80°C. Fibroblasts were grown to 30% confluency and incubated with 3 ml/dish of viral supernatant supplemented with 8 μg/ml of polybrene for 6 h at 37°C. Afterwards, cells were washed twice with 1x PBS, and fresh medium was added. Two days later, the selection process was started by addition of 0.5 μg/ml puromycin (Sigma, Munich, Germany; #P8833) for 10-15 days, with medium changes every 2-3 days, to achieve stable transfection. As controls, either scrambled shRNA (SC) or empty vector (EV) transfected fibroblasts were generated. All five shRunx2 immortalized fibroblast cell lines and human primary fibroblasts were characterized for efficiency of Runx2 knock-down via qRT-PCR and western blotting, and one shRunx2 per cell line (B8 for immortalized fibroblasts and C9 for human fibroblasts) was chosen based on its significant Runx2 down-regulation for downstream functional analysis compared to their respective SC or EV controls (Fig. S7H-I; Table S12).

**Table 3.**
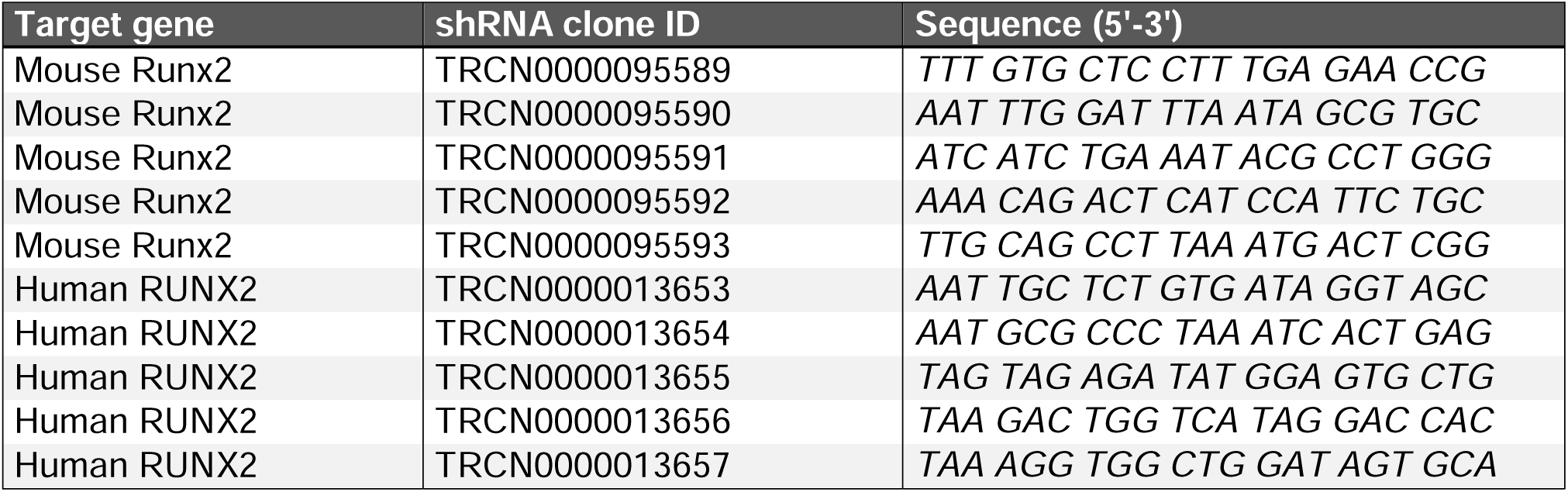
shRNA sequences

### Transient transfection of human fibroblasts with siRNA

Human CAFs and normal fibroblasts were transfected with siRNA using the Lipofectamine® RNAiMAX Reagent kit (Invitrogen) according to the manufacturer’s instructions. The RUNX2 siRNAs were previously validated: siRUNX2.1 was used in human keloid fibroblasts [95]; siRUNX2.2 was used in human renal cell carcinoma cells [96]; a validated non-targeting negative control sequence was used as siRNA control (Microsynth AG, Balgach, Switzerland) (Table 4). *RUNX2* knock-down was confirmed by qRT-PCR, as described below.

**Table 4.**
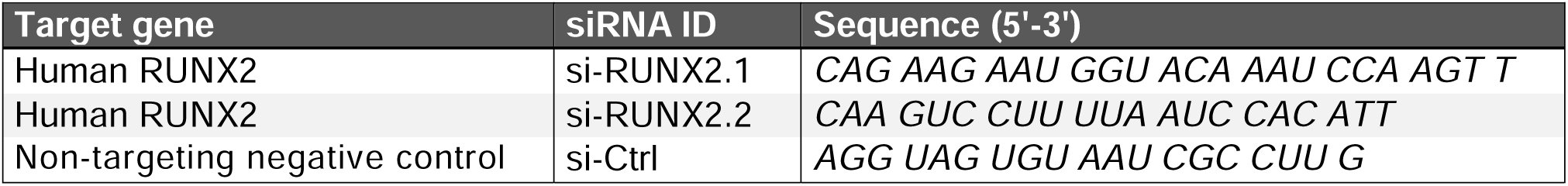
siRNA sequences

### Proliferation assay

To analyze cell proliferation, cells were seeded at 20,000 cells/well into a 6-well plate and counted at 24 h, 48 h, 72 h, and 96 h post-seeding after Trypan Blue staining using a hemocytometer.

### Migration assay

To analyze cell migration, a scratch assay was performed. The cells were seeded at 50,000 cells/well in a 6-well plate and grown to 100% confluency. They were cultured in DMEM/10% FBS/1% P/S and treated with mitomycin C (1:100) for 2 h at 37°C. A scratch was made vertically from the top to the bottom of each well using a sterile P200 pipette tip. The medium was aspirated and the cells washed carefully with PBS. Normal culture medium was added and the scratched area was photographed and analyzed after 6 h, 12 h, 24 h, and 48 h.

### Collagen gel contraction assay

To study collagen gel contraction by fibroblasts, 24-well plates were coated with 1% BSA at 37°C for 1 h. For each gel, 120 μl HEPES, 26 μl 10× DMEM, 16 μl Milli-Q water, and 240 μl of cell suspension (420,000 cells/ml) were combined. 198 μl of cold TeloCol Type I Bovine Collagen Solution (Advanced BioMatrix, San Diego, CA; #5026) was added to the mixture and immediately applied to each well. The 24-well plate containing the gels was placed at 37°C and incubated for 30 min to allow the gel to solidify. Afterwards, 800 μl of DMEM was added to the wells containing the gels. Using a sterile P200 pipette tip, the sides of each well were scraped gently, to ensure that the gels did not stick to the walls. Gels were photographed next to a ruler in view at 0 h and after 2 h, 12 h, and 24 h.

### Analysis of the pro-tumorigenic potential of fibroblast-derived secretomes and matrisomes

Human CAFs and normal fibroblasts were transiently transfected with siRNA and plated at 70-90% confluency in DMEM/10% FBS/P/S. On the following day, they were pre-treated with 2 μg/ml mitomycin C and cultured in starvation medium (DMEM/1% FBS/P/S) for an additional 3 days. Conditioned media and ECM were prepared as previously described [91]. SCC13 cells were seeded in CM or on ECM in 24-well plates. The cells were co-cultured in complete DMEM for 3 days and subsequently measured for the SCC13-LifeAct (rLV-Ubi-LifeAct Lentiviral Vectors RFP-Tag, #60142, Vitaris, Baar, Switzerland) colony area.

### Western blot analysis

Preparation of protein lysates and Western blot were performed using standard procedures. Protein concentration of the samples was measured using the bicinchoninic acid (BCA) kit (Thermo Fisher Scientific, Waltham, MA) according to the manufacturer’s instructions. After blocking in 5% BSA in TBS-T for 1 h, primary antibodies, including anti-β-actin (Abcam, Cambridge, UK; #ab8229), anti-collagen I (Southern Biotech, Birmingham, AL; #1310-01), anti-lamin A (Santa Cruz, Santa Cruz, CA; #sc-6214), anti-mouse Runx2 (Abcam; #ab114133), and anti-human RUNX2 (Sigma; #HPA022040), were diluted at 1:1000 in 5% BSA in TBS-T and incubated overnight at 4°C, followed by incubation with horsradish peroxidase (HRP)-conjugated secondary antibodies (Promega, Madison, WI). The chemiluminescence signals were detected using a Western blot developing machine (Fusion Solo S chemiluminescence imager) using EvolutionCapt Solo 6S software.

### RNA isolation and qRT-PCR

RNA was isolated from cultured cells using Trizol (Life Technologies, Carlsbad, CA) or the Mini Total RNA kit (IBI Scientific, Dubuque, IA) and reverse transcribed using the iScript cDNA synthesis kit (BioRad, Hercules, CA), all according to the manufacturers’ instructions. Quantitative PCR was performed using LightCycler 480 SYBR Green I Master reaction mix (Roche, Rotkreuz, Switzerland) and the primers listed in Table 5. Data were quantified using second derivative maximum analysis and gene expression represented as relative to the internal housekeeping gene *Rps2*9 (for mouse samples) or *RPL27* (for human samples).

**Table 5.**
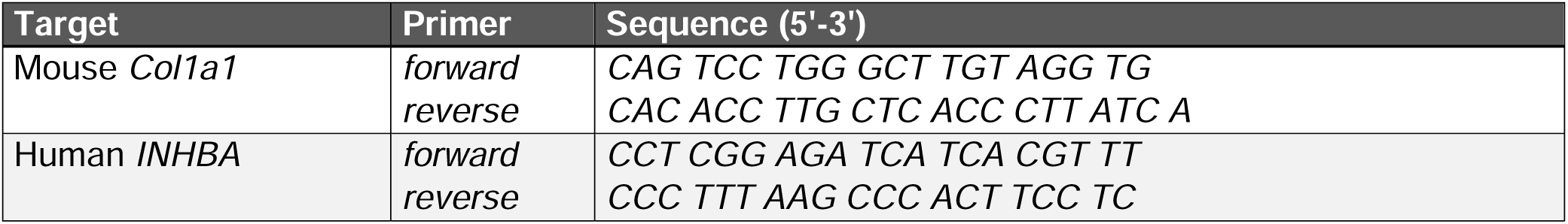
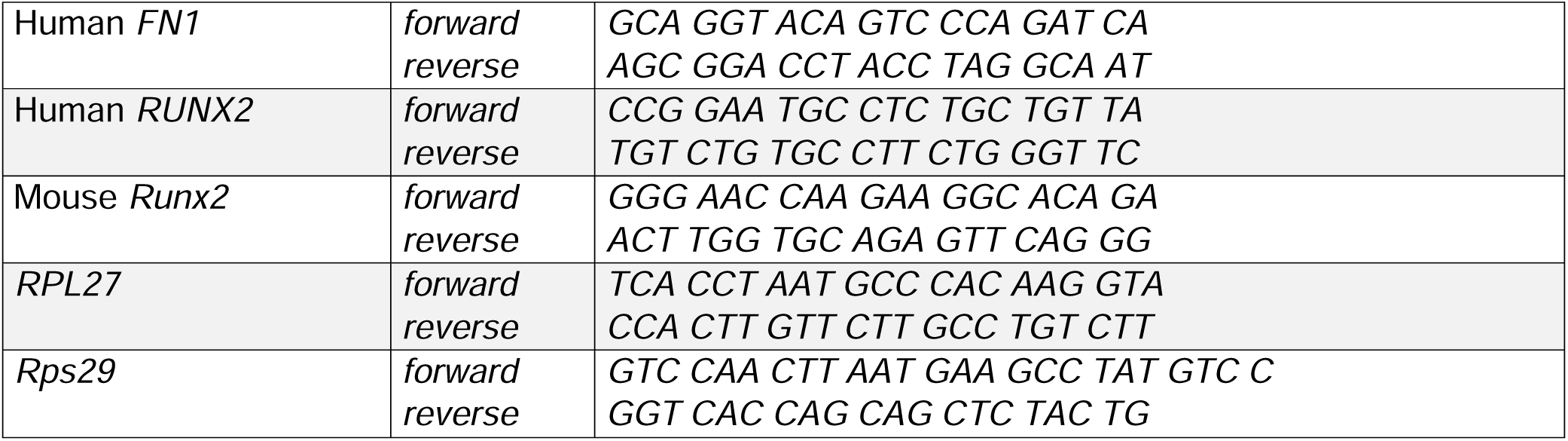
Primers used for qRT-PCR

### Meta-analysis of bulk CAF datasets

Microarray and RNA-seq gene expression profiles of CAFs from different tissues were acquired from Gene Expression Omnibus (GEO) using the GEOquery R package (version 2.38.4) (Table S1). RNA-seq data was processed using CLC Genomics Workbench version 10 (Qiagen). Affymetrix microarray data based on Human Genome U133 Plus 2.0 and Affymetrix Human Exon 1.0 ST arrays were normalized in batches of CEL files by means of the robust multi-array average (RMA) method in the affy R package (version 1.50.0). Probe-set mapping to Gene Symbols was retrieved from GEO for all arrays. Probes that map to the same gene were averaged. Log2 fold-changes of the expression in CAF samples relative to the mean expression across the respective normal fibroblast samples was calculated for each gene and CAF sample. When paired samples of cancer and normal samples from the same patient were available, log2 fold-changes were calculated for the pairs instead of the mean across the normal samples.

Individual gene expression matrices were harmonized by combining them into a single matrix comprising genes shared across all datasets. The combined matrix was filtered based on the wound fibroblast phase-specific genes or on CAF phase-predictive genes, and CAFs were correlated to wound fibroblast samples using Spearman correlation via the *cor* function in the stats R package (version 4.0.2). Annotated heatmaps of WFB-CAF correlations and CAF gene-gene correlations were generated using the ComplexHeatmap R package (version 2.6.2). The CAF gene-gene correlation heatmap was clustered according to Ward, allowing for identification of major gene clusters in the dataset, two of which were mutually exclusive to one another, leading to the identification of CAF early and late phase-predictive genes (Table S6). Functional enrichment analysis of CAF phase-predictive genes was performed using the Cytoscape (version 3.8.2) plugin ClueGO (version 2.5.8) [80], and gene-gene interaction networks with functional enrichment were generated using the Cytoscape plugin StringApp (version 1.6.0) [86].

### Analysis of scRNA-seq datasets

ScRNA-seq datasets were downloaded (Table S1) and filtered based on original author-generated cell type annotations to contain only fibroblasts and CAFs using the Matrix R package (version 1.3-2). Filtered fibroblast datasets were analyzed using the Seurat R package (version 4.0.2), according to the published data analysis and integration protocol [97]. Individual Seurat objects were created from each dataset and the following quality control measures were taken. For each cell in a dataset, mitochondrial and ribosomal RNA percentages were calculated via *PercentageFeatureSet* for patterns “^MT-”, “^RPL” and “^RPS”, respectively; RNA feature counts (nFeature_RNA) were plotted against mitochondrial and ribosomal RNA percentages via *FeatureScatter*; the datasets were subset to include only cells expressing >500 RNA features and relatively low percentages of mitochondrial and ribosomal RNA (see Table S20 for dataset-specific settings). The datasets were normalized according to the *SCTransform* protocol [98] using standard settings and regressing out the mitochondrial and ribosomal RNA features via the *vars.to.regress* modifier. The normalized datasets were analyzed via the *RunPCA* function, followed by *ElbowPlot* to visualize the standard deviation of the principal components and to select the most significant dimensions for further analyses (see Table S20 for dataset-specific settings). The following functions were then run using standard settings and the dataset-specific number of principal comments via the *dims* modifier: *RunUMAP*, *FindNeighbors*.

Scoring of gene expression signatures in datasets was done via the *AddModuleScore* function using standard settings and gene sets that included the wound fibroblast early and late phase-specific genes, early and late phase-predictive genes, and genes comprising the Reactome pathways “collagen formation”, “elastic fibre formation”, and “smooth muscle contraction”. The signature scores were visualized on the UMAP plot via the *FeaturePlot* function. The custom red/blue color gradients were added to the plots with the RColorBrewer R package (version 1.3) and the *scale_colour_gradientn* function.

Trajectory analyses were performed on the datasets using the Monocle3 R package (version 1.2.9) [99]. The standard protocol was followed, as outlined on the Monocle3 website (https://cole-trapnell-lab.github.io/monocle3/docs/trajectories/), except that partitions were set to 1 in order to connect separated clusters of cells. Selected wFB phase-specific and CAF phase-predictive gene expressions were normalized via z-score, clustered via K-means using 3 groups, and plotted along the pseudotime using the ComplexHeatmap R package (version 2.12) and the *Heatmap* function. As validation, the CytoTrace web tool (https://cytotrace.stanford.edu/) was also used to analyze the wound fibroblast dataset in order to obtain a predicted differentiation trajectory [100], which largely matched the Monocle-predicted trajectory.

Cluster-based analyses were performed on the datasets via the *FindClusters* function to group cells that clearly clustered together according to their UMAP projections while minimizing over-clustering of the data (see Table S20 for dataset-specific settings). In all CAF datasets, there were three major clusters identified with unique transcriptional profiles enriching for the early (including collagen and contractile) and late (including elastin) wound signatures. In the wFB dataset, four major clusters were identified, since there was a sizeable population of normal or unwounded skin fibroblasts with a unique signature separate from the collagen, elastin and contractile populations. Differential expression analyses were done to identify statistically-significant differentially expressed genes between the cell clusters via the *FindAllMarkers* function, using parameters: min.pct = 0.25, logfc.threshold = 0.25. Visualizations were generated by the *VlnPlot* and *DotPlot* functions for violin plots and dot plots, respectively, using selected wFB phase-specific or CAF phase-predictive genes as input. Dataset-specific clusters and DEGs are provided in Table S7 for mouse and Table S9 for human datasets.

### Analysis of spatial RNA-seq datasets

Spatial RNA-seq datasets were downloaded (Table S1) and re-analyzed using the Seurat R package (version 4.0.2). 10x Genomics Visium data were uploaded using the *Load10X_Spatial* function and normalized via *SCTransform* function using standard settings. ST data were first manually uploaded as separate sample, spot and image files, Seurat objects were created via the *InputFromTable* function, and the data were normalized via *SCTransform* function using standard settings. Scoring of gene expression signatures in datasets was done via the *AddModuleScore* function using standard settings and gene sets that included the wound fibroblast early and late phase-specific genes, early and late phase-predictive genes, and the genes comprising the Reactome pathways “collagen formation”, “elastic fibre formation”, and “smooth muscle contraction”. The signature scores were overlaid on the histological sections via the *SpatialDimPlot* function.

### Global co-expression analysis of *POSTN* and *MFAP4*

Co-expression analyses of *RUNX2*, *POSTN*, and *MFAP4* across all cancer types was performed using the Search-Based Exploration of Expression Compendium (SEEK) [101]. The top co-expressed genes (P-value<0.01 for all; co-expression score >1 for *POSTN* and *MFAP4*; co-expression score >0.6 for *RUNX2*) were compared using Venny 2.1 and functionally enriched using EnrichR. The co-expressed gene lists and full functional enrichment lists are provided (Table S13).

### Human skin tumor and wound biopsies

Normal human skin and skin cancer samples were obtained anonymously from the Department of Dermatology, University Hospital of Zurich, in the context of the Biobank project and approved by the local and cantonal Research Ethics Committees. Primary and metastatic melanoma was diagnosed by experienced pathologists. Written informed consent was obtained from all subjects and the experiments conformed to the principles set out in the WMA Declaration of Helsinki and the Department of Health and Human Services Belmont Report. Biopsies were fixed and paraffin embedded, and sections were prepared for histological analysis.

We selected 33 patient samples from a previously published primary melanoma tissue micro-array (TMA) [42], which had paired biopsies from the tumor center and tumor border (66 total tumor cores), for ECM imaging. The selected samples were linked to clinico-histopathological metadata, including immune cell staining, clinical ulceration, recurrence and survival. We selected samples containing undamaged stromal areas in paired tumor center/border biopsy tumor cores to enable comparisons of ECM between tumor locations. Patient metadata for the analyzed set of tumor cores is provided (Table S14). Adjacent sections of the TMA cores were previously stained for melan A (tumor cells) and CD45 (immune cells), and the scanned images were compared to the ECM imaging data.

Human wound samples were obtained anonymously from a healing kinetics study with healthy volunteers (trial number 002WH99) undertaken by SWITCH BIOTECH GmbH, Neuried, Germany, at the Clinic and Polyclinic for Dermatology and Allergology at Biederstein of the Technical University of Munich, Germany, after approval by the local Research Ethics Committee. Briefly, 6-mm diameter punch biopsies were taken from the margins of acute skin wounds at 5-, 8-, 12-, and 15-days post-injury, fixed, and paraffin embedded, and 5 μm sections were prepared for histological analysis.

### Histology, immunostaining, and image analysis

For immunofluorescence staining of cultured cells, coverslips were fixed in 4% PFA, permeabilized with 0.1% Triton X100 in 1x PBS for 30min, blocked with 1% BSA, and incubated overnight with the anti-Ki67 (1:1000; Abcam; #ab15580), anti-α-SMA (1:500; Sigma; #A2547), anti-α-SMA-FITC (1:1000; Sigma; #F3777) primary antibodies, or with phalloidin-AlexaFluor 555 (1:200; Thermo Fisher Scientific; #A34055), followed by donkey anti-rabbit AlexaFluor 488 (1:1000; Jackson ImmunoResearch, West Grove, PA) or AlexaFluor 594 (1:1000; Jackson ImmunoResearch) secondary antibodies, and counterstained with Hoechst 33342 (Sigma).

For immunofluorescence staining of mouse skin wounds, fresh tissue samples were embedded in optimal cutting temperature (OCT) compound, frozen, and 7 μm sections were cut at the cryostat. The wound sections were air-dried for 20 min in the dark and rehydrated in 1x PBS for 10 min. The sections were fixed in pre-cooled acetone at −20°C for 5 min. The sections were blocked with 10% normal donkey serum (Jackson ImmunoResearch; #017-000-121) in PBS-T for 90 min, incubated overnight in the dark with anti-Runx2 primary antibody (1:100; Abcam; #ab114133), followed by donkey anti-rabbit AlexaFluor 647 secondary antibody (1:1000; Jackson ImmunoResearch; #711-605-152), and counterstained with Hoechst 33342 (Sigma).

For immunofluorescence staining of human tumors, paraffin sections were dewaxed and rehydrated using a xylene/ethanol gradient followed by antigen retrieval using pre-heated citrate buffer at 98°C for 20 min and washing in PBS-T. The sections were blocked with 10% normal donkey serum (Jackson ImmunoResearch; #017-000-121) and incubated overnight with 1) goat anti-collagen I (1:200; SouthernBiotech; #1310-01) and rabbit anti-MFAP4 (1:100; Invitrogen, Waltham, MA; # PA5-62917), 2) mouse anti-periostin (1:100; Santa Cruz; #sc-398631) and rabbit anti-MFAP4 (1:100; Invitrogen; # PA5-62917), or 3) goat anti-fibronectin (1:200; Santa Cruz; #sc-6953) and rabbit anti-MFAP4 (1:100; Invitrogen; # PA5-62917), followed by donkey anti-goat AlexaFluor 594 (#705-585-003), donkey anti-goat AlexaFluor 647 (#705-605-14), donkey anti-rabbit AlexaFluor 647 (#711-605-152) or donkey anti-rabbit AlexaFluor 488 (#711-547-003) secondary antibodies (1:2000; Jackson ImmunoResearch), and counterstained with DAPI (Sigma).

Fluorescence images were acquired with a Zeiss AxioImager.M2 microscope with a motorized stage at ×10, ×20, and ×40 magnification. Zen Pro software (Carl Zeiss AG, Oberkochen, Germany) was used to control the camera and to stitch together individual photomicrographs into wound- or coverslip-spanning images.

### Human wound and tumor ECM imaging and analysis

Second-harmonic generation (SHG) imaging was performed on dewaxed and rehydrated sections, as previously described [40, 41]. SHG and auto-fluorescence (AF) signals were collected to specifically visualize collagen via SHG and elastin via AF, as previously described [38, 39]. Imaging was carried out using a Leica TCS SP8 microscope equipped with a ×25 0.95□numerical aperture (NA) L Water HCX IRAPO objective and a Mai Tai XF (Spectra-Physics) MP laser tunable from 720–950□nm. For ECM imaging, sequential scan SHG was generated using a 900 nm excitation laser: the SHG collagen signal was detected at 450 nm, and the AF elastin signal was detected at 505-550 nm. Wounds and tumor cores were imaged one at a time over several sessions, using 30-50 tiles and 2-4 z-planes per core that were subsequently merged into single images, to account for tissue thickness and sample angling. Laser power was monitored and kept constant throughout each experiment, as were photomultiplier tube (PMT) voltage, gain and offset. Leica LAS AF SP8 (version 4.0) software was used to control the instrument, for image acquisition, tile merging and file export.

Exported whole tumor core stacks were edited in Fiji (version 2.1.0). The z-stacks were collapsed via *ZProjection* type “Max Intensity,” the SHG (collagen) and AF (elastin) channels were split and separately thresholded to maximize ECM fiber signal and minimize noise. At this step, obvious artifacts were manually removed from the images. The separate channels were exported for downstream analysis or merged in RGB color, with SHG as red and AF as green, and exported for figure preparation. The exported tumor core images were combined in TMA format using Adobe Illustrator 2020 (version 24.0.1), though the ECM fiber analysis was performed on individual tumor core images. To delineate tumor margins within tumor cores, the previously melan A-stained adjacent cores were visualized in QuPath (version 0.2.3), opened side-by-side with the ECM images and used to manually draw tumor margin masks.

CT-FIRE (version 3.0) [102] was used to analyze the divergent ECM architectures of collagen and elastin images at the individual fiber level in batch using standard settings. CurveAlign (version 5.0) [102] was then used to analyze the ECM fiber parameters in relation to the tumor margins in batch using standard settings (diagrammed in Fig. S10B). The ECM fiber parameters, which were considered for downstream analysis, are diagrammed in Fig. 8A. All mean ECM fiber parameters including clinico-histopathological data were combined into a single data matrix and imported into GraphPad Prism (version 9) for Spearman correlation analyses. Individual fiber level data per core were exported and combined into individual data matrices according to biopsy location and fiber type and analyzed in GraphPad Prism using Histogram (binned to 50 pixel units) and Spearman correlation analyses. The correlation matrices were exported along with their respective P-values (Table S15).

Multivariate Cox proportional hazards regression modelling via the survival R package (version 3.2-11) was used to evaluate the prognostic potential of tumor core collagen and elastic fiber parameters in terms of survival and recurrence. The combined data matrix was imported into R, center-normalized and scaled using the *scale* function, and split according to biopsy location (tumor center and tumor border) and ECM fiber type (collagen and elastin), and four types of analyses were performed, separately for survival and recurrence. All ECM fiber parameters were added to the model using the *coxph* function, including: fiber length, fiber width, nearest fiber distance, nearest fiber alignment, overall fiber alignment, overall fiber orientation, nearest distance to tumor margin, and nearest relative angle to tumor margin. The model was evaluated using the *summary* and *anova* functions, and features were removed from the model sequentially via the *drop1* function until a model remained with universally significant or near-significant features. Potential confounders such as gender and age did not significantly change the results of the survival and recurrence analyses when they were added to the multivariate Cox models. The final model tables were generated via the *prettify* function of the papeR package (version 1.0-5). Full model details are available (Table S16).

### Statistical Analysis

Statistical analyses were performed using Prism for MacOS (ver. 9) and R (ver. 4.0.2) in RStudio (ver. 1.3.959). For comparison of two groups, the two-sided Student’s t-test was used; for comparison of more than two groups, one way or two-way ANOVA and Bonferroni’s multiple comparisons post hoc tests were used. For Spearman correlation analyses, two-sided P values were computed for each pair of variables tested. For multivariate Cox models, co-variates were removed stepwise by single term deletions of highest Chi-square p-values (via *drop1* function) until co-variates remained with p-values < 0.10 (via *summary* function).

## ACKNOWLEDGEMENTS

We thank Chelsea Chen, ETH Zurich, for experimental help, Dr. Patrick Turko (University Hospital Zurich) for help with statistical modeling, Sol Taguinod (ETH Zurich Phenomics Center) for help with the mouse maintenance, Dr. Malgorzata Kisielow and Anette Schütz (ETH Zurich Flow Cytometry Core Facility) for help with the flow cytometry experiments, Catharine Aquino (Functional Genomics Center Zurich) for performing the RNA-sequencing experiments, Justine Kusch (Scientific Center for Optical and Electron Microscopy at ETH) for microscopy assistance, Dr. Hans-Dietmar Beer (University of Zurich) for providing human primary fibroblasts, and Dr. Petra Boukamp (Leibniz Institute for Environmental Research, Düsseldorf, Germany) for providing SCC13 cells.

This work was supported by the Swiss National Science Foundation, grants 31003A_169204 and 31003B-189364 (SW), Cancer Research Switzerland, grant KFS-4510-08-2018 (SW), and the ETH Zurich Open ETH Project SKINTEGRITY.CH (SW). MC, MPL, RD and SW are members of the SKINTEGRITY.CH collaborative research program.

## AUTHOR CONTRIBUTIONS

Conceptualization: MSW, SW

Methodology: MSW, DL, MCa, MCl, MPL, RD

Investigation: MSW, DL, MCa, SS, JJ, AG

Visualization: MSW, DL

Funding acquisition: SW

Project administration: MSW, SW

Supervision: MSW, SW

Writing – original draft: MSW, SW

Writing – review & editing: all authors

## COMPETING INTERESTS

RD has intermittent, project focused consulting and/or advisory relationships with Novartis, Merck Sharp & Dhome (MSD), Bristol-Myers Squibb (BMS), Roche, Amgen, Takeda, Pierre Fabre, Sun Pharma, Sanofi, Catalym, Second Genome, Regeneron, Alligator, T3 Pharma, MaxiVAX SA, Pfizer and touchIME, all outside of the submitted work. No other authors have potential competing interests to declare.

## DATA AVAILABILITY

The authors declare that all data supporting the findings of this study are available within the article and its supplementary materials, or from the corresponding authors upon request. RNA-seq data from bulk-sorted skin papilloma CAFs have been deposited in the Gene Expression Omnibus database under accession code: GSE182880 (reviewer token: krcpocqydbutbkv). All other transcriptomics datasets analyzed in this manuscript are listed in Table S1. The Biopeak R package introduced in this manuscript has been deposited and documented in CRAN at the following URL: https://CRAN.R-project.org/package=Biopeak.

## SUPPLEMENTAL INFORMATION TITLES AND LEGENDS

### Supplementary Figures

**Fig. S1. Wound healing phase-specific genes, tumor phase-predictive genes, and their prognostic analysis.**

(A) Comparison of genes across time-points from a published repair meta-analysis [12], showing many overlaps between time-points. True time-resolved genes were extracted and grouped according to Inflammatory-Proliferative (red), Proliferative-Resolution (green) and Non-specific (blue) categories. (B) Comparison of healing phase-specific genes to time-resolved genes from (A), showing an input-normalized percentage (%) of shared phase-specific genes with gene sets from Inflammation (I) and/or Early repair (ER) and/or Late repair (LR) and/or Remodeling (R), or Non-specific (Non-Sp) genes.

(C) Functional enrichment analysis of healing phase-specific genes, showing top five Gene Ontology (GO) Biological Process pathways and their statistical significance.

(D) Ingenuity Pathways Analysis (IPA) of healing phase-specific genes via the Biological Functions analysis, showing the Activation Z-score of selected functions for each phase.

(E) Comparison of genes across time-points in transcriptomic datasets of human excisional wound biopsies from skin and oral buccal mucosa [15]. Only relatively highly abundant transcripts (RPKM > 1) that were more abundant in wounds compared to respective intact, unwounded tissues (Log_2_ fold change (FC) > 2) were considered.

(F) Comparison of genes across time-points in transcriptomic datasets of human split-thickness skin graft biopsies [16]. Only genes that were over-expressed in wounds compared to intact, unwounded skin (Log_2_ fold change (FC) > 1.5, false discovery rate (FDR) < 0.05) were considered. True time-resolved genes were extracted and grouped according to Inflammatory-Proliferative (red) and Proliferative-Resolution (green) categories.

(G) Examples of shared genes from the comparisons of phase-predictive genes and time-resolved human split-thickness skin grafts.

(H) Comparison of genes associated with increased severity in non-melanoma skin cancer and precursor lesions, showing the overlap of 200 top up-regulated genes, relative to respective intact skin samples, in squamous cell carcinoma (SCC) compared to actinic keratosis (AK) and in treatment-resistant basal cell carcinoma (rBCC) compared to treatment-sensitive BCC (sBCC). Example overlap genes are shown.

(I) Plot of individual Proliferative and Resolution wound scores in skin (melanoma), breast and lung tumor samples according to the phase-predictive genes. Large labeled circles and boxes indicate mean wound score for each tumor type for phase-specific and phase-predictive genes, respectively.

(J) Kaplan-Meyer plots of TCGA patient samples median-stratified according to resolution wound scores (top) and proliferative wound scores (bottom) in primary melanoma (left) and melanoma metastases (right). The log-rank P-value is shown for each plot comparing the survival of patients stratified according to the top and bottom median wound scores.

(K) Comparison of the Core Serum Response (CSR) gene set [7] to inflammatory, proliferative and resolution wound phase-specific and tumor phase-predictive genes, showing the percentage (%) of shared CSR genes with time-resolved gene sets.

(L) Comparison of pan-cancer prognostic proliferative and resolution phase-predictive genes according to the TCGA and PRECOG databases.

(M) Functional enrichment analysis of genes associated with worse (left) or better (right) patient outcome in skin malignant melanoma using both TCGA and PRECOG databases. FDR: false discovery rate.

**Fig. S2. Early and late wound fibroblast signatures enrich for divergent matrix pathways.**

(A) Prediction of cell type according to the prognostic (worse patient outcome) proliferative and resolution phase-predictive genes via the Immunological Genome Project, showing high enrichment in fibroblastic reticular cells within the stromal cells category.

(B) Functional enrichment networks generated by the ClueGO Cytoscape plugin, showing the wound fibroblast (wFB) “Early” (red) and “Late” (green) phase-specific genes co-enriched in ECM-related Reactome pathways.

(C) Comparison of wound fibroblast Early and Late phase-specific genes with whole wound (WW) Early, Proliferative and Late phase-specific genes, showing example overlap genes.

(D) UMAP plots showing the distributions of Monocle trajectory pseudotimes (left) and CytoTrace trajectory differentiation scores (right) of wFBs from small excisional skin wounds.

(E) UMAP plot showing mouse wFB cells from large excisional wounds at 12- and 18-days post-wounding colored by dataset of origin.

(F) UMAP plot showing distribution of early (left) and late (right) wFB phase-specific gene signature scores.

(G) UMAP plot showing distribution of selected Reactome pathway gene signature scores.

(H) UMAP plot showing Monocle-based trajectory path and distribution of pseudotime values.

(I) Clustered heatmap showing normalized expression of selected wFB phase-specific genes along the trajectory pseudotime.

**Fig. S3. Single-cell human melanoma CAF analyses reveal divergent early *vs* late CAF subtype signatures.**

(A) UMAP plot showing distribution of early (left) and late (right) CAF phase-predictive gene signature scores in human melanoma-derived CAFs.

(B) UMAP plot showing distribution of selected Reactome pathway gene signature scores in melanoma CAFs.

(C) UMAP plot showing melanoma CAFs colored by transcriptional cluster number.

(D) Violin plots showing the distribution of early and late CAF phase-predictive and selected Reactome pathway gene signature scores in melanoma CAF clusters identified in (D).

(E) Dot plot showing the average normalized expression (node color) and percent expression (node size) of selected CAF phase-predictive genes in melanoma CAF clusters identified in (D).

**Fig. S4. Additional single-cell analyses in non-skin tumor types.**

(A-E) Early and late CAF signatures and selected Reactome pathway signatures were scored in scRNA-seq analyzed CAFs derived from human breast (BC, A), lung (LC, B), colorectal (CRC, C), head and neck squamous cell (HNSC, D), and ovarian (OC, E) tumors and visualized via UMAP plots showing their distribution across the datasets.

(F) Datasets of non-skin CAFs (A-E) and skin-derived CAFs from BCC and SCC were integrated, analyzed, and visualized in a UMAP plot colored by tumor type of origin.

(G) UMAP plots showing showing the distributions of early and late CAF signature and selected Reactome pathway signature scores across the pan-CAF landscape.

(H) UMAP plot showing the integrated CAF dataset in cluster-based analysis resulting in 3 main CAF clusters.

(I) Violin plots showing the distribution of early and late CAF phase-predictive and selected Reactome pathway gene signature scores in pan-CAF clusters identified in (H).

(J) Dot plot showing the average normalized expression (node color) and percent expression (node size) of selected CAF phase-predictive genes in pan-CAF clusters identified in (H).

**Fig. S5. Additional spatial RNA-seq analyses of skin SCC sections.**

(A-C) Spatial RNA-seq analyses of skin SCC sections, analyzed via Visium (A) and via early Spatial Transcriptomics (ST; B and C). Left: H&E-stained section diagrams indicating locations of the tumor edges with black dashed lines. Middle: Feature plots showing the spatial distribution of early (left) and late (right) CAF phase-predictive gene signature scores overlaying the SCC sections. Right: Feature plots showing the spatial distribution of selected Reactome pathway gene signature scores overlaying the SCC sections. Scale bars = 500 μm.

**Fig. S6. RNA-seq analysis of bulk-sorted skin wound- and cancer-derived fibroblasts.**

(A) Flow cytometry gating strategy for sorting of fibroblasts from mouse back skin, 5-day wounds, and papilloma tumors.

(B) Representative H&E staining of a piece of papilloma from which fibroblasts were isolated. Scale bar = 100 µm.

(C) Comparison of 100 top expressed genes from each RNA-seq category, showing example universally-overlap genes, which include major fibroblast markers.

(D) Prediction of cell type according to the 54 top-expressed overlapped genes from (C) via the Immunological Genome Project, showing high enrichment in fibroblastic reticular cells within the stromal cells category.

(E) Principal component analysis (PCA) plot of RNA-seq data from wound fibroblasts (wFB), cancer-associated fibroblasts (CAF), and respective fibroblasts from normal, intact skin (NS).

(F) Gene set enrichment analysis (GSEA) comparing genes up-regulated in early wFBs and CAFs with published datasets of wFB scRNA-seq clusters [27], showing normalized enrichment scores (NES) and statistical significance; FDR: false discovery rate.

(G) Comparison of the genes enriched by leading edge analysis in at least 3 datasets by GSEA of wFB and CAF up-regulated genes with all wFB datasets, showing examples of the top enriched genes; bolded shared enriched genes are common myofibroblast and/or CAF markers.

(H) Table showing top 25 GSEA co-enriched genes of wFB and CAF up-regulated genes with the listed published wFB datasets, ranked by the CAF leading edge combined enrichment score; bolded shared enriched genes are common myofibroblast and/or CAF markers; dataset details are listed in Table S1.

**Fig. S7. RUNX2 as an upstream regulator of major wound fibroblast and CAF genes and generation of RUNX2-knockdown fibroblast cell lines.**

(A) Ingenuity Pathway Analysis (IPA) Upstream Regulator analysis of shared early wound fibroblast (wFB) and CAF up-regulated genes, showing the most significant predicted Upstream Regulators and their P-values, ranked by their Activation z-scores (>2, <-2).

(B) Staining of sections from normal (unwounded) back skin and 5-day excisional skin wounds of PDGFRa-eGFP mice for Runx2 (red); fibroblasts express nuclear eGFP (green) and nuclei are counterstained with Hoechst (blue). Top row represents the “no primary antibody” control, showing non-specific staining in the upper epidermal layers and hair follicles of healthy skin and wounds. Inset is the zoomed-in area within the wound bed indicated by the white rectangle, showing co-expression of Runx2 and eGFP in the nuclei of stromal cells. Scale bars = 50 μm.

(C) Representative flow cytometric plots from back skin tumors showing the initial gating strategy for live cell singlets.

(D-G) Representative flow cytometric plots showing the gating strategy for Runx2^high^ fibroblasts (CD49f^-^, CD45^-^, CD140a^+^, Runx2^high^ cells) in back skin tumors (G), intact back skin (D), 3-day wounds (E), and 7-day wounds (F).

(H) Representative live-cell brightfield photographs showing the morphology of mouse immortalized fibroblast (mImmFB) cell lines #1 & #3 and human primary fibroblasts (hpFB) stably transduced with shRUNX2 lentiviruses (shRUNX2) compared to controls. Scale bar = 100 μm.

(I) Western blot analyses showing Runx2 expression in shRunx2-transduced mImmFB cell lines #1-3 and pFB compared to those transduced with viruses carrying an empty vector (EV) or scrambled shRNA (SC) controls; β-actin or lamin A were used as loading controls for whole cell or nuclear extracts, respectively.

(J) Quantification of Runx2 protein expression (normalized to loading controls) in shRunx2 mImmFB and human pFB relative to EV or SC controls.

Bars show mean ± SEM; n=3, except n=2 for human pFB; shRunx2 *vs* control. *p<0.05, **p<0.01 (unpaired Student *t*-test).

(K) qRT-PCR analysis showing relative *Runx2* expression in shRunx2-transduced mImmFB and human pFB relative to EV or SC controls; expression data are normalized to *Rps29* for mImmFB or *RPL27* for human pFB. Bars show mean ± SEM; n=3 for all, except n=1 for human pFB; shRunx2 *vs* control *p<0.05, **p<0.01, ****p<0.0001 (unpaired Student *t*-test).

**Fig. S8. RUNX2 regulates major fibroblast functions.**

(A) Cell numbers over time of shRunx2 (blue) mouse immortalized fibroblasts (mImmFB) and human primary fibroblasts (hpFB) compared to controls (grey) (n=4).

(B) Representative immunofluorescence staining for the proliferation marker Ki-67 (red or green) in shRunx2 mouse immortalized fibroblast (mImmFB) cell lines #2-3 and human primary fibroblasts compared to controls; nuclei are counterstained with Hoechst (blue). Quantification of these data is shown in Fig 5H.

(C) Representative photographs of collagen gels after 20 h of contraction by shRunx2 mImmFB cell lines #1-3 compared to controls. Quantification of these data is shown in in Fig 6H.

(D) Representative immunofluorescence stainings for α-smooth muscle actin (red) and phalloidin fluorescence (green) in shRunx2 immFB cell line #2 compared to controls; nuclei are counterstained with Hoechst (blue).

(E) qRT-PCR for *INHBA* and *FN1* relative to *RPL27* using RNA from siRUNX2-transfected primary SCC CAFs compared to scrambled siRNA controls (n=2-3). Scale bars = 100 μm. Bars show mean ± SEM; *p<0.05, **p0.01, ****p<0.0001 (Student’s t-test (E), two-way ANOVA followed by Bonferroni’s post-tests (A)).

**Fig. S9. Collagen type I, fibronectin, periostin, MFAP4, collagen and elastin fibers in normal human skin, tumors and wounds.**

(A-D) Immunofluorescence co-staining for fibronectin (A-C, red) or collagen type I (D, red) and MFAP4 (green) in sections of human superficial (A) and deep skin (B), subcutaneous melanoma edge (C-D); nuclei are counterstained with DAPI (blue). Yellow dashed lines represent epidermal-dermal junctions (A-B) or tumor margins (C-D). Individual channels are shown in black and white. (E, F) Picrosirius red staining for collagen of human superficial (E, left and middle) and deep skin (E, right), subcutaneous melanoma core (F), visualized in brightfield (left) or polarized light (middle, right). Scale bars = 200 μm.

(G) Visualization via multiphoton microscopy by second-harmonic generation (SHG) of collagen (red) and auto-fluorescence (AF) for elastin (green) in a section of human superficial and deep skin stained with an antibody against MFAP4 (cyan) and counter-stained with DAPI (blue), showing individual channels in black and white.

(H) Inset detail of the area marked by the white square in (G), showing significant fibrous co-localization of elastin AF signal (middle) and MFAP4 staining (right) in the dermis. Scale bars = 200 μm.

(I-L) Visualization via multiphoton microscopy by second-harmonic generation (SHG) for collagen (red) and auto-fluorescence (AF) for elastin (green) of subcutaneous melanoma (I), BCC (J), and human excisional wound sections from 5-, 8-, 12-, and 15-days post-injury (K-L). Insets (white boxes in zoomed out images) show detailed organization of collagen and elastin fibers at the tumor margins (I-J) and at the edges of wounds (K, insets in L); individual channels are shown in black and white. Scale bars = 200 μm.

Black or white dashed lines represent epidermal-dermal junctions (A, B, G, I) or tumor margins (C, D, I, J).

**Fig. S10. Collagen/elastin imaging and ECM fiber analysis of primary melanoma tissue microarray.**

(A) Composite image showing the primary melanoma tissue micro-array (TMA) of tumor cores individually imaged via multiphoton microscopy by second-harmonic generation (SHG) for collagen (red) and auto-fluorescence (AF) for elastin (green).

(B) Diagram showing the major features of (ECM fiber analysis of tumor cores by CT-FIRE/CurveAlign, taking into account the points of reference for calculation of ECM fiber parameters, especially those that are relative to the manually-drawn borders of the tumor margin within each tumor core.

(C) Spearman correlation matrix of mean ECM fiber parameters across tumor cores for collagen and elastin fibers at the tumor border (top) or the tumor center (bottom).

(D) Spearman correlation matrix of individual ECM fiber parameters within individual tumor cores for collagen (left) or elastin (right) fibers at the tumor border (top) or the tumor center (bottom).

(E) Representative melanoma tumor border core, showing sections stained for tumor cells (melanA, left), immune cells (CD45, center) and ECM fibers (collagen/elastin, right). Inset shows the spatial relationship between the tumor margin, immune cell infiltrate, and high density of elastin fibers (green).

(F, G) Additional Multivariate Cox regression models of ECM fiber parameters significantly impacting patient survival (F) or melanoma recurrence (G), showing the collagen (red) and elastin (green) fiber parameters contributing to statistically-significant hazard ratios (HR) at the tumor border (top) and center (bottom).

Black or white dashed lines represent tumor margins. FT: fiber type; P-val: P-value; L: lower bound; U: upper bound.

**Fig S11. Summary figure.**

(A) Diagram of the whole tissue bioinformatics pipeline identifying healing phase-specific genes from analysis of the wound healing spectrum and tumor phase-predictive genes from comparisons to pan-cancer datasets, and focusing on the most prognostic genes and pathways within the mid and late phases of healing, which enriched for collagen and elastin formation.

(B) Diagram of the fibroblast-centric bioinformatics pipeline identifying wound fibroblast phase-specific genes and CAF phase-predictive genes from comparisons to pan-CAF datasets, which enriched for contraction and collagen formation in the “early” group and for elastin formation in the “late” group. ScRNA-seq meta-analyses of wound fibroblast and CAF datasets confirmed the existence of these functionally distinct “early” and “late” CAF subtypes, and re-analyses of spatial transcriptomics cancer datasets confirmed the presence of these CAF subtypes in distinct niches of the TME.

(C) Conceptual summary of the manuscript: During wound healing, sequential healing phases return the injured tissue to homeostasis. As a “wound that fails to heal”, cancer stalls in the early phases of repair, with cancer cells and TME cells continually releasing cytokines and growth factors to exacerbate the early wound phenotypes, including ECM formation, resulting in fibrosis. Typical wound fibroblast functions of the early healing phase become hyper-activated in “early-wound” CAF subtypes in close proximity to the tumor margins, contributing to the accumulation of pro-tumorigenic collagen-forming and contractile CAFs that exacerbate the fibrotic ECM microenvironment. Further away from the tumor margins, there is a gradient toward the normalization of the tumor-adjacent microenvironment mediated by “late-wound” CAF subtypes that deposit an elastin-enriched ECM network similar to that found in a resolving wound.

### Supplementary Tables

Table S1. List and description of datasets analyzed in this manuscript.

Table S2. Peak detected wound phase-specific genes, functional enrichment analysis.

Table S3. Analysis of human wound-related datasets.

Table S4. Phase-predictive genes in TCGA tumors, functional enrichment analysis.

Table S5. Survival analysis of wound-associated genes in all tumor types and in skin melanoma, functional enrichment of Proliferative- and Resolution-phase prognostic genes.

Table S6. Wound fibroblast phase-specific genes, functional enrichment analysis.

Table S7. Differentially-expressed genes in single cell RNA-seq analyses of mouse wound fibroblast and CAF datasets.

Table S8. Pan-CAF phase-predictive genes, functional enrichment analysis.

Table S9. Differentially-expressed genes in single cell RNA-seq analyses of human skin and pan CAF datasets.

Table S10. Differentially-expressed genes in wound fibroblast and CAF bulk RNA-seq analyses, functional enrichment analysis.

Table S11. Gene set enrichment analysis of early wound- and tumor-derived fibroblasts, leading edge analysis.

Table S12. Description of fibroblast cell lines used in experiments.

Table S13. Global gene co-expression analyses of RUNX2, POSTN and MFAP4 in cancer.

Table S14. Patient demographics of analyzed samples from primary melanoma tissue microarray.

Table S15. Details of ECM parameter correlations and clinico-histopathological correlations.

Table S16. Multi-variate Cox proportional hazards regression model analyses of ECM parameters.

Table S17. Wound score robustness analysis: Spearman vs Pearson correlation pair-wise comparison values.

Table S18. Wound score robustness analysis: leave-one-out standard deviation values.

Table S19. List of all machine learning models and their associated performance metrices.

Table S20. List of bioinformatics analysis settings for analyzed single-cell datasets.

