## Supplementary Information for "Phase-specific signatures of wound fibroblasts and matrix patterns define cancer-associated fibroblast subtypes"

**Supplementary Figures S1-S11**

**Supplementary Tables S1-S20**

**Tables S1, S12, S14 are shown below**

**Tables S2-S11, S13, S15-S20 are uploaded as separate Excel files**

**Supplementary Figure S1**

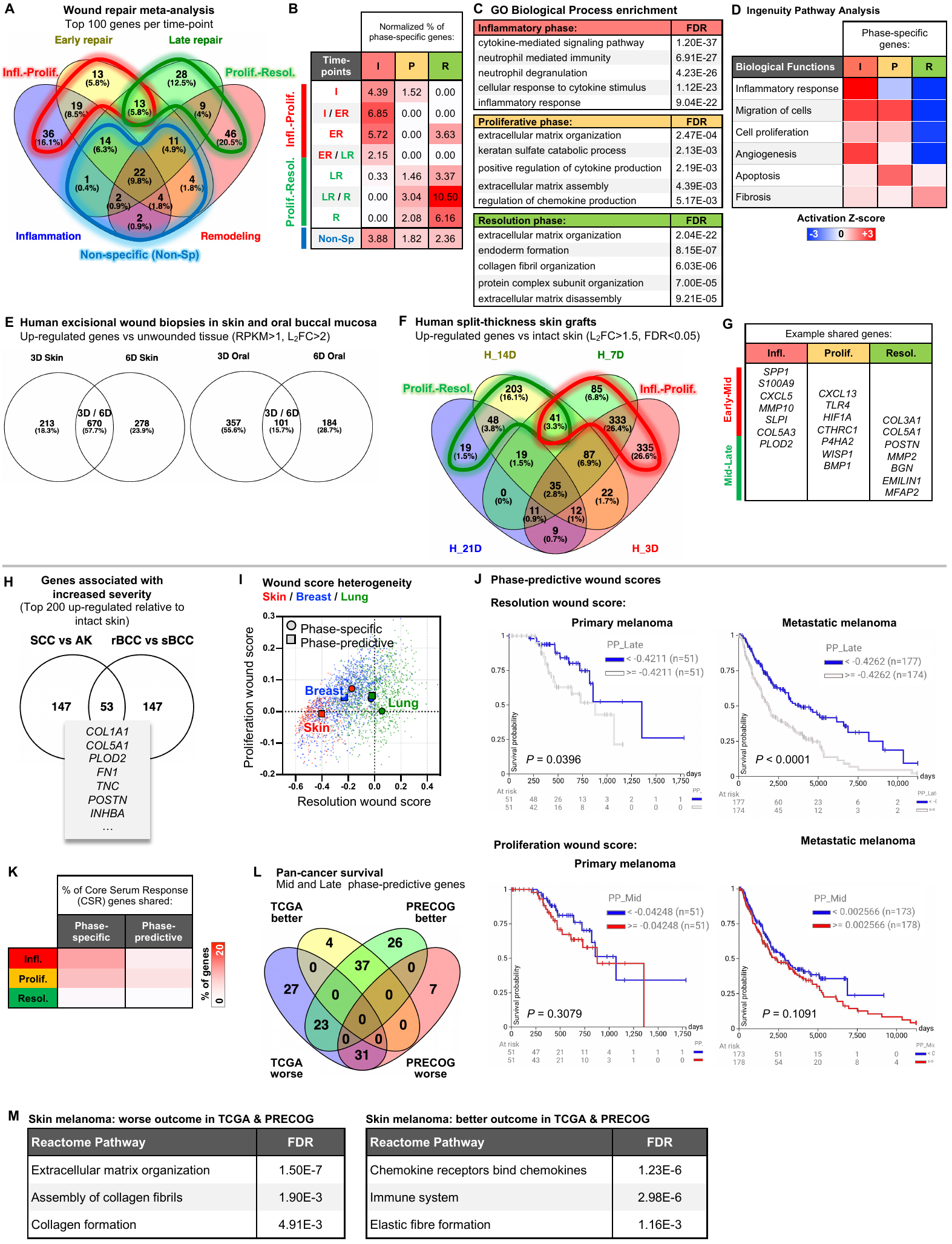

**Fig. S1. Wound healing phase-specific genes, tumor phase-predictive genes, and their prognostic analysis.**

(A) Comparison of genes across time-points from a published repair meta-analysis [12], showing many overlaps between time-points. True time-resolved genes were extracted and grouped according to Inflammatory-Proliferative (red), Proliferative-Resolution (green) and Non-specific (blue) categories. (B) Comparison of healing phase-specific genes to time-resolved genes from (A), showing an input-normalized percentage (%) of shared phase-specific genes with gene sets from Inflammation (I) and/or Early repair (ER) and/or Late repair (LR) and/or Remodeling (R), or Non-specific (Non-Sp) genes.

(C) Functional enrichment analysis of healing phase-specific genes, showing top five Gene Ontology (GO) Biological Process pathways and their statistical significance.

(D) Ingenuity Pathways Analysis (IPA) of healing phase-specific genes via the Biological Functions analysis, showing the Activation Z-score of selected functions for each phase.

(E) Comparison of genes across time-points in transcriptomic datasets of human excisional wound biopsies from skin and oral buccal mucosa [15]. Only relatively highly abundant transcripts (RPKM > 1) that were more abundant in wounds compared to respective intact, unwounded tissues (Log_2_ fold change (FC) > 2) were considered.

(F) Comparison of genes across time-points in transcriptomic datasets of human split-thickness skin graft biopsies [16]. Only genes that were over-expressed in wounds compared to intact, unwounded skin (Log_2_ fold change (FC) > 1.5, false discovery rate (FDR) < 0.05) were considered. True time-resolved genes were extracted and grouped according to Inflammatory-Proliferative (red) and Proliferative-Resolution (green) categories.

(G) Examples of shared genes from the comparisons of phase-predictive genes and time-resolved human split-thickness skin grafts.

(H) Comparison of genes associated with increased severity in non-melanoma skin cancer and precursor lesions, showing the overlap of 200 top up-regulated genes, relative to respective intact skin samples, in squamous cell carcinoma (SCC) compared to actinic keratosis (AK) and in treatment-resistant basal cell carcinoma (rBCC) compared to treatment-sensitive BCC (sBCC). Example overlap genes are shown.

(I) Plot of individual Proliferative and Resolution wound scores in skin (melanoma), breast and lung tumor samples according to the phase-predictive genes. Large labeled circles and boxes indicate mean wound score for each tumor type for phase-specific and phase-predictive genes, respectively.

(J) Kaplan-Meyer plots of TCGA patient samples median-stratified according to resolution wound scores (top) and proliferative wound scores (bottom) in primary melanoma (left) and melanoma metastases (right). The log-rank P-value is shown for each plot comparing the survival of patients stratified according to the top and bottom median wound scores.

(K) Comparison of the Core Serum Response (CSR) gene set [7] to inflammatory, proliferative and resolution wound phase-specific and tumor phase-predictive genes, showing the percentage (%) of shared CSR genes with time-resolved gene sets.

(L) Comparison of pan-cancer prognostic proliferative and resolution phase-predictive genes according to the TCGA and PRECOG databases.

(M) Functional enrichment analysis of genes associated with worse (left) or better (right) patient outcome in skin malignant melanoma using both TCGA and PRECOG databases. FDR: false discovery rate.

**Supplementary Figure S2**

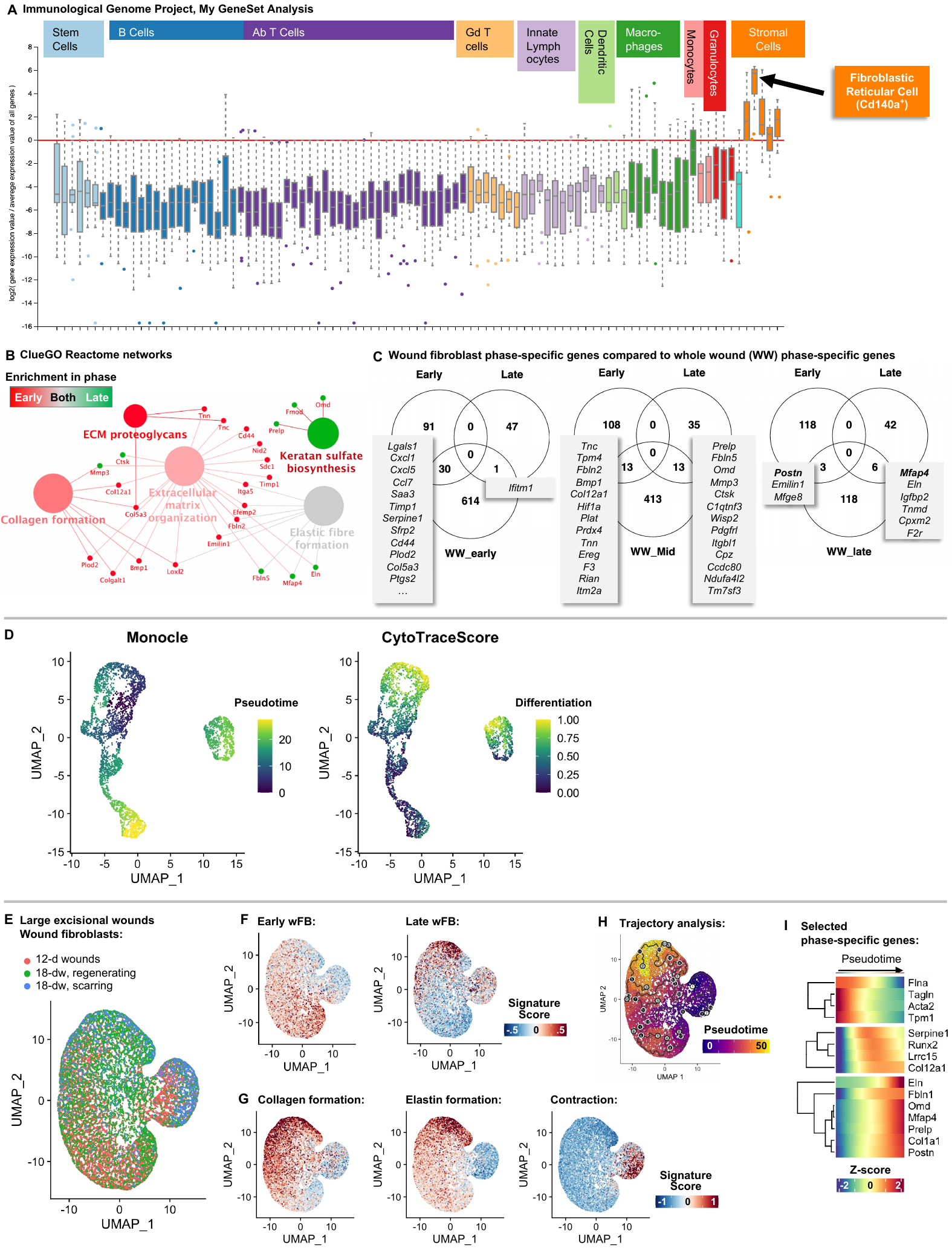

**Fig. S2. Early and late wound fibroblast signatures enrich for divergent matrix pathways.**

(A) Prediction of cell type according to the prognostic (worse patient outcome) proliferative and resolution phase-predictive genes via the Immunological Genome Project, showing high enrichment in fibroblastic reticular cells within the stromal cells category.

(B) Functional enrichment networks generated by the ClueGO Cytoscape plugin, showing the wound fibroblast (wFB) “Early” (red) and “Late” (green) phase-specific genes co-enriched in ECM-related Reactome pathways.

(C) Comparison of wound fibroblast Early and Late phase-specific genes with whole wound (WW) Early, Proliferative and Late phase-specific genes, showing example overlap genes.

(D) UMAP plots showing the distributions of Monocle trajectory pseudotimes (left) and CytoTrace trajectory differentiation scores (right) of wFBs from small excisional skin wounds.

(E) UMAP plot showing mouse wFB cells from large excisional wounds at 12- and 18-days post-wounding colored by dataset of origin.

(F) UMAP plot showing distribution of early (left) and late (right) wFB phase-specific gene signature scores.

(G) UMAP plot showing distribution of selected Reactome pathway gene signature scores.

(H) UMAP plot showing Monocle-based trajectory path and distribution of pseudotime values.

(I) Clustered heatmap showing normalized expression of selected wFB phase-specific genes along the trajectory pseudotime.

**Supplementary Figure S3**

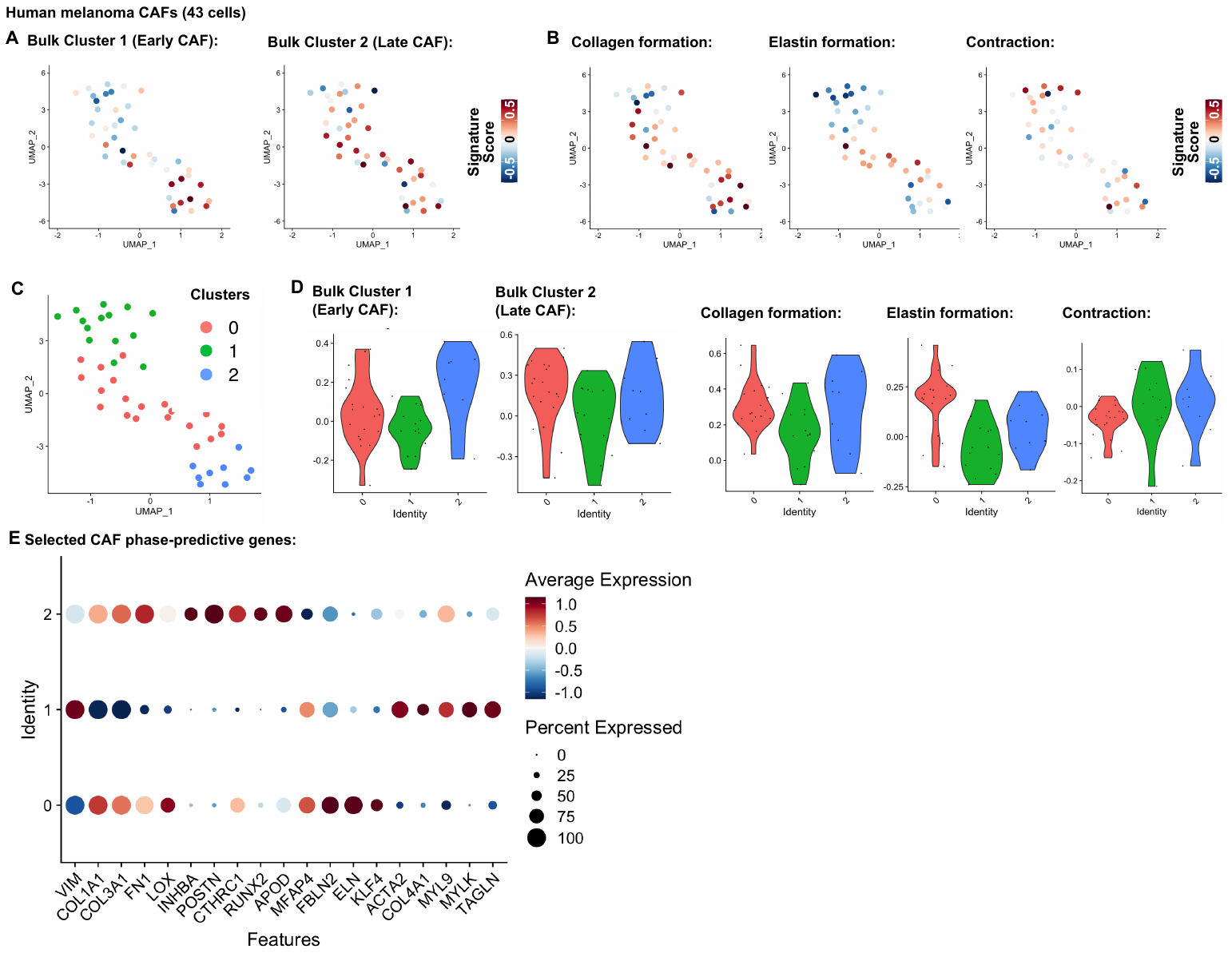

**Fig. S3. Single-cell human melanoma CAF analyses reveal divergent early *vs* late CAF subtype signatures.**

(A) UMAP plot showing distribution of early (left) and late (right) CAF phase-predictive gene signature scores in human melanoma-derived CAFs.

(B) UMAP plot showing distribution of selected Reactome pathway gene signature scores in melanoma CAFs.

(C) UMAP plot showing melanoma CAFs colored by transcriptional cluster number.

(D) Violin plots showing the distribution of early and late CAF phase-predictive and selected Reactome pathway gene signature scores in melanoma CAF clusters identified in (D).

(E) Dot plot showing the average normalized expression (node color) and percent expression (node size) of selected CAF phase-predictive genes in melanoma CAF clusters identified in (D).

**Supplementary Figure S4**

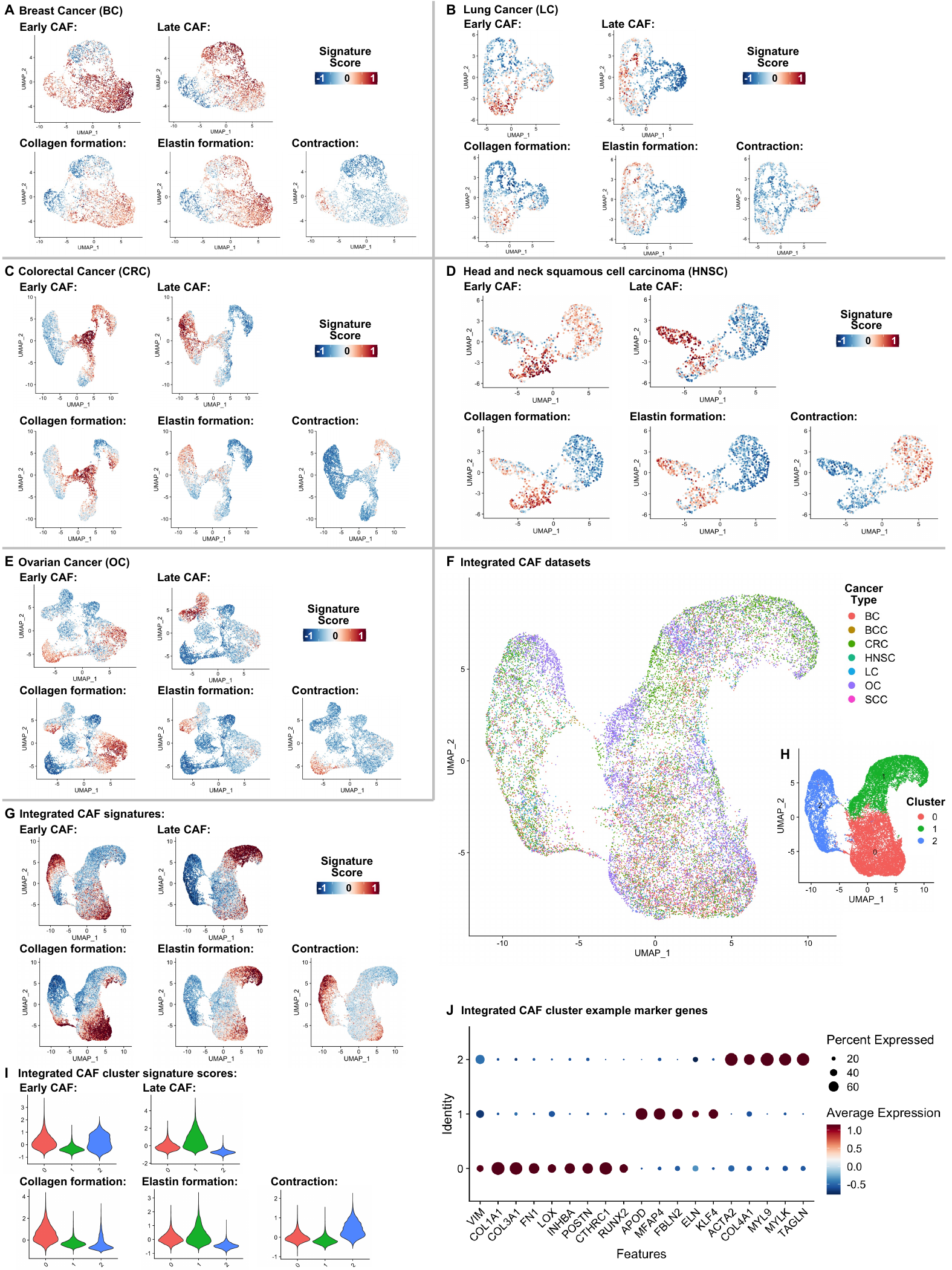

**Fig. S4. Additional single-cell analyses in non-skin tumor types.**

(A-E) Early and late CAF signatures and selected Reactome pathway signatures were scored in scRNA-seq analyzed CAFs derived from human breast (BC, A), lung (LC, B), colorectal (CRC, C), head and neck squamous cell (HNSC, D), and ovarian (OC, E) tumors and visualized via UMAP plots showing their distribution across the datasets.

(F) Datasets of non-skin CAFs (A-E) and skin-derived CAFs from BCC and SCC were integrated, analyzed, and visualized in a UMAP plot colored by tumor type of origin.

(G) UMAP plots showing showing the distributions of early and late CAF signature and selected Reactome pathway signature scores across the pan-CAF landscape.

(H) UMAP plot showing the integrated CAF dataset in cluster-based analysis resulting in 3 main CAF clusters.

(I) Violin plots showing the distribution of early and late CAF phase-predictive and selected Reactome pathway gene signature scores in pan-CAF clusters identified in (H).

(J) Dot plot showing the average normalized expression (node color) and percent expression (node size) of selected CAF phase-predictive genes in pan-CAF clusters identified in (H).

**Supplementary Figure S5**

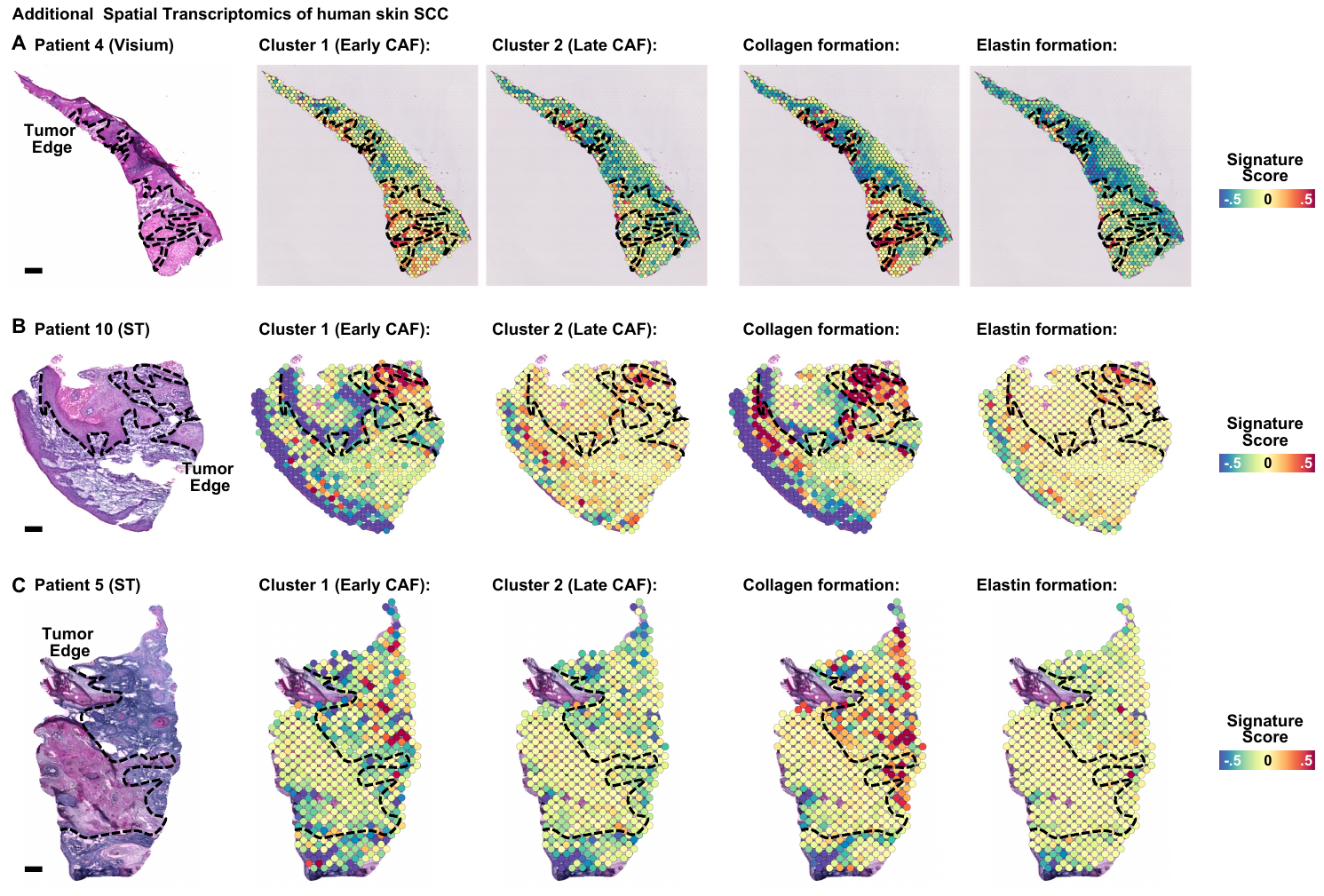

**Fig. S5. Additional spatial RNA-seq analyses of skin SCC sections.**

(A-C) Spatial RNA-seq analyses of skin SCC sections, analyzed via Visium (A) and via early Spatial Transcriptomics (ST; B and C). Left: H&E-stained section diagrams indicating locations of the tumor edges with black dashed lines. Middle: Feature plots showing the spatial distribution of early (left) and late (right) CAF phase-predictive gene signature scores overlaying the SCC sections. Right: Feature plots showing the spatial distribution of selected Reactome pathway gene signature scores overlaying the SCC sections. Scale bars = 500 μm.

**Supplementary Figure S6**

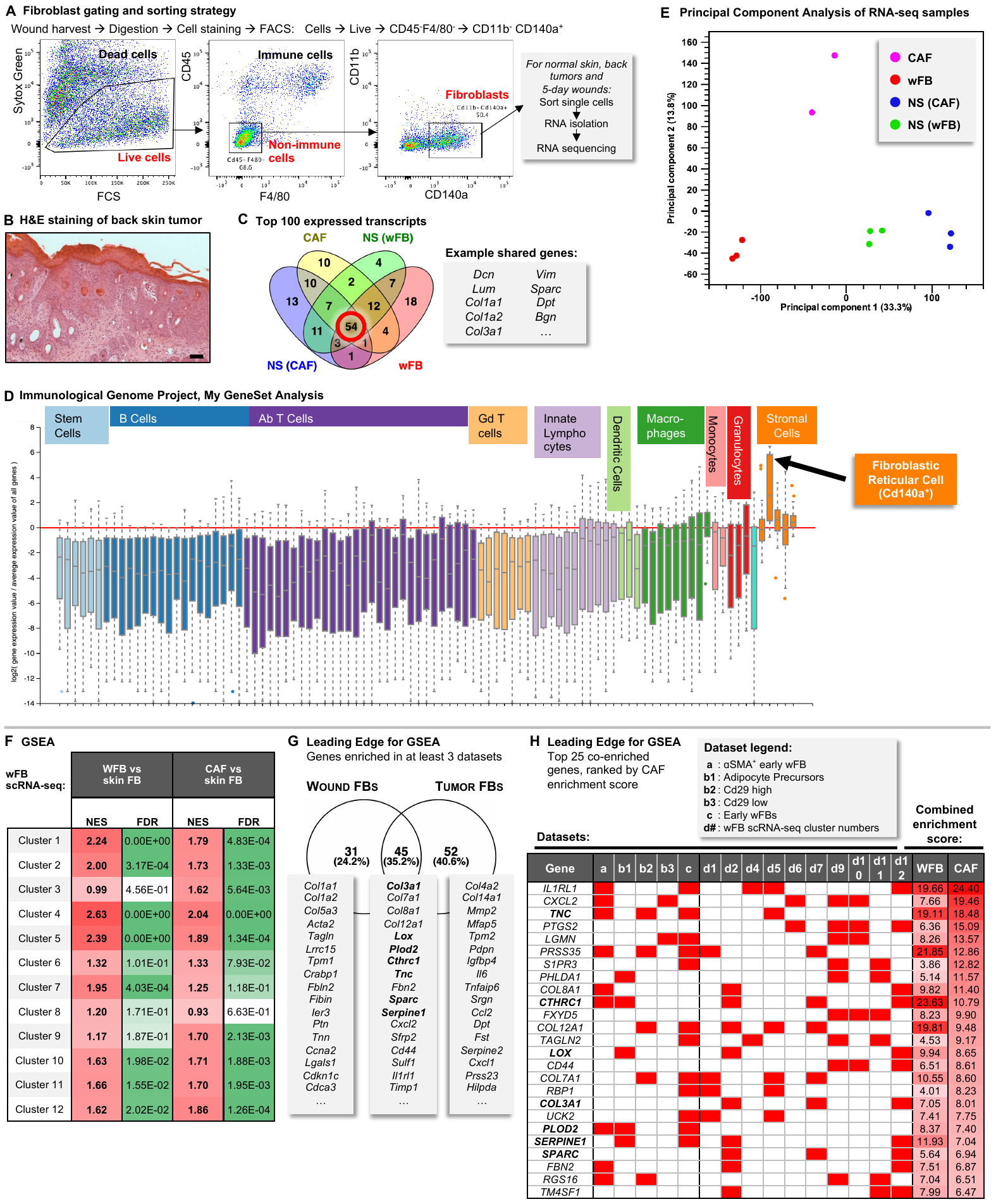

**Fig. S6. RNA-seq analysis of bulk-sorted skin wound- and cancer-derived fibroblasts.**

(A) Flow cytometry gating strategy for sorting of fibroblasts from mouse back skin, 5-day wounds, and papilloma tumors.

(B) Representative H&E staining of a piece of papilloma from which fibroblasts were isolated. Scale bar = 100 µm.

(C) Comparison of 100 top expressed genes from each RNA-seq category, showing example universally-overlap genes, which include major fibroblast markers.

(D) Prediction of cell type according to the 54 top-expressed overlapped genes from (C) via the Immunological Genome Project, showing high enrichment in fibroblastic reticular cells within the stromal cells category.

(E) Principal component analysis (PCA) plot of RNA-seq data from wound fibroblasts (wFB), cancer-associated fibroblasts (CAF), and respective fibroblasts from normal, intact skin (NS).

(F) Gene set enrichment analysis (GSEA) comparing genes up-regulated in early wFBs and CAFs with published datasets of wFB scRNA-seq clusters [27], showing normalized enrichment scores (NES) and statistical significance; FDR: false discovery rate.

(G) Comparison of the genes enriched by leading edge analysis in at least 3 datasets by GSEA of wFB and CAF up-regulated genes with all wFB datasets, showing examples of the top enriched genes; bolded shared enriched genes are common myofibroblast and/or CAF markers.

(H) Table showing top 25 GSEA co-enriched genes of wFB and CAF up-regulated genes with the listed published wFB datasets, ranked by the CAF leading edge combined enrichment score; bolded shared enriched genes are common myofibroblast and/or CAF markers; dataset details are listed in Table S1.

**Supplementary Figure S7**

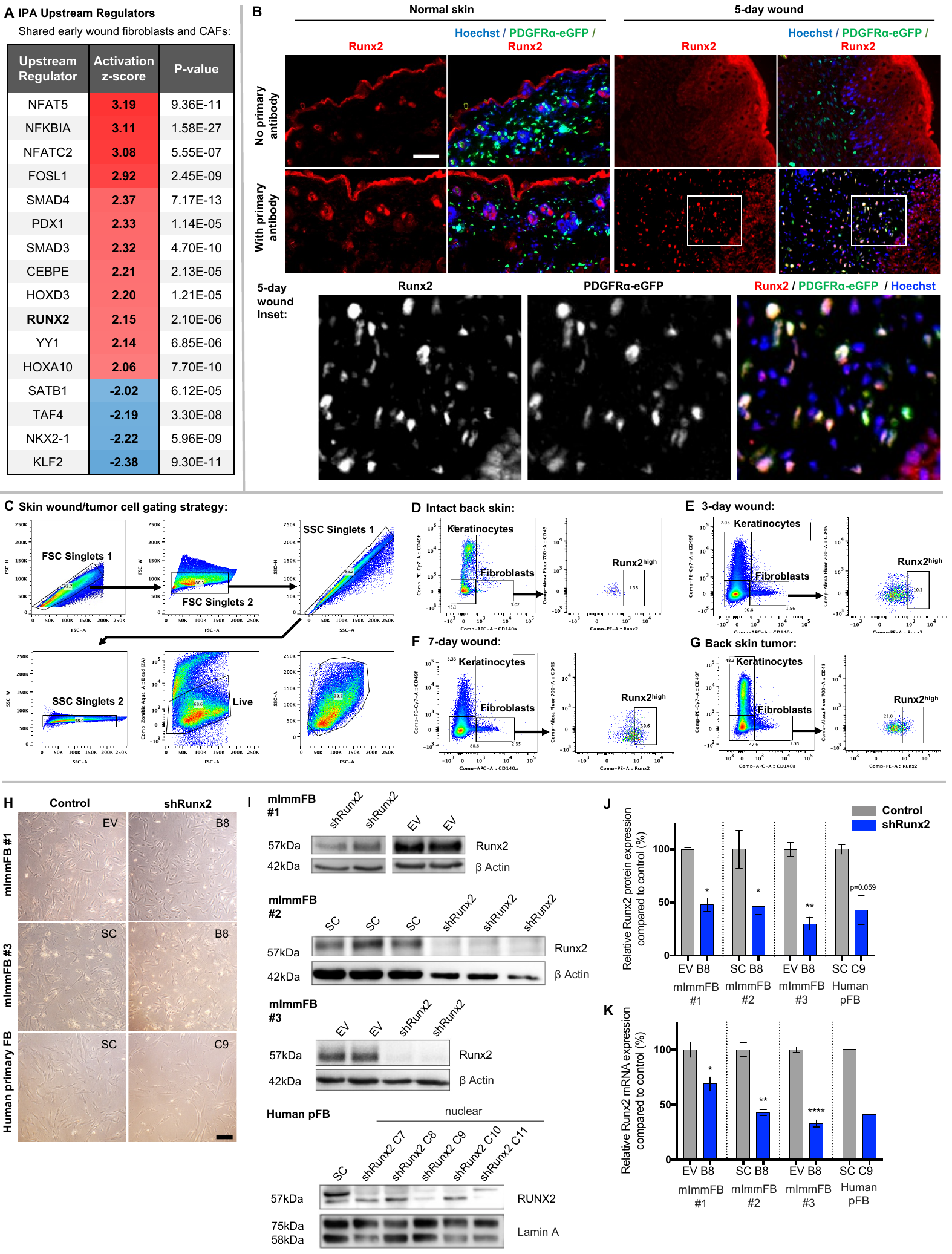

**Fig. S7. RUNX2 as an upstream regulator of major wound fibroblast and CAF genes and generation of RUNX2-knockdown fibroblast cell lines.**

(A) Ingenuity Pathway Analysis (IPA) Upstream Regulator analysis of shared early wound fibroblast (wFB) and CAF up-regulated genes, showing the most significant predicted Upstream Regulators and their P-values, ranked by their Activation z-scores (>2, <-2).

(B) Staining of sections from normal (unwounded) back skin and 5-day excisional skin wounds of PDGFRa-eGFP mice for Runx2 (red); fibroblasts express nuclear eGFP (green) and nuclei are counterstained with Hoechst (blue). Top row represents the "no primary antibody" control, showing non-specific staining in the upper epidermal layers and hair follicles of healthy skin and wounds. Inset is the zoomed-in area within the wound bed indicated by the white rectangle, showing co-expression of Runx2 and eGFP in the nuclei of stromal cells. Scale bars = 50 μm.

(C) Representative flow cytometric plots from back skin tumors showing the initial gating strategy for live cell singlets.

(D-G) Representative flow cytometric plots showing the gating strategy for Runx2^high^ fibroblasts (CD49f^-^, CD45^-^, CD140a^+^, Runx2^high^ cells) in back skin tumors (G), intact back skin (D), 3-day wounds (E), and 7-day wounds (F).

(H) Representative live-cell brightfield photographs showing the morphology of mouse immortalized fibroblast (mImmFB) cell lines #1 & #3 and human primary fibroblasts (hpFB) stably transduced with shRUNX2 lentiviruses (shRUNX2) compared to controls. Scale bar = 100 μm.

(I) Western blot analyses showing Runx2 expression in shRunx2-transduced mImmFB cell lines #1-3 and pFB compared to those transduced with viruses carrying an empty vector (EV) or scrambled shRNA (SC) controls; β-actin or lamin A were used as loading controls for whole cell or nuclear extracts, respectively.

(J) Quantification of Runx2 protein expression (normalized to loading controls) in shRunx2 mImmFB and human pFB relative to EV or SC controls.

Bars show mean ± SEM; n=3, except n=2 for human pFB; shRunx2 *vs* control. *p<0.05, **p<0.01 (unpaired Student *t*-test).

(K) qRT-PCR analysis showing relative *Runx2* expression in shRunx2-transduced mImmFB and human pFB relative to EV or SC controls; expression data are normalized to *Rps29* for mImmFB or *RPL27* for human pFB. Bars show mean ± SEM; n=3 for all, except n=1 for human pFB; shRunx2 *vs* control *p<0.05, **p<0.01, ****p<0.0001 (unpaired Student *t*-test).

**Supplementary Figure S8**

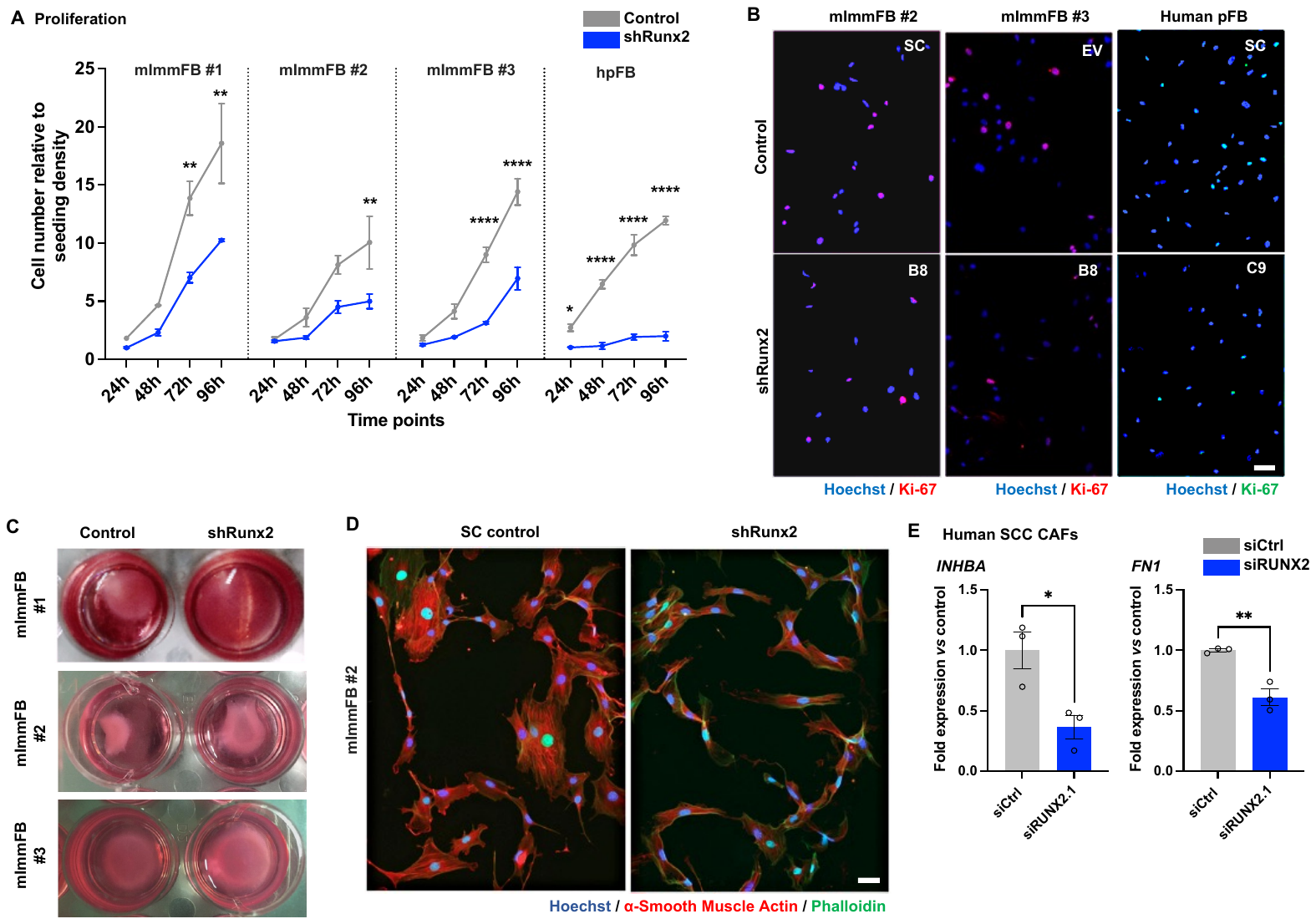

**Fig. S8. RUNX2 regulates major fibroblast functions.**

(A) Cell numbers over time of shRunx2 (blue) mouse immortalized fibroblasts (mImmFB) and human primary fibroblasts (hpFB) compared to controls (grey) (n=4).

(B) Representative immunofluorescence staining for the proliferation marker Ki-67 (red or green) in shRunx2 mouse immortalized fibroblast (mImmFB) cell lines #2-3 and human primary fibroblasts compared to controls; nuclei are counterstained with Hoechst (blue). Quantification of these data is shown in Fig 5H.

(C) Representative photographs of collagen gels after 20 h of contraction by shRunx2 mImmFB cell lines #1-3 compared to controls. Quantification of these data is shown in in Fig 6H.

(D) Representative immunofluorescence stainings for α-smooth muscle actin (red) and phalloidin fluorescence (green) in shRunx2 immFB cell line #2 compared to controls; nuclei are counterstained with Hoechst (blue).

(E) qRT-PCR for *INHBA* and *FN1* relative to *RPL27* using RNA from siRUNX2-transfected primary SCC CAFs compared to scrambled siRNA controls (n=2-3). Scale bars = 100 μm. Bars show mean ± SEM; *p<0.05, **p0.01, ****p<0.0001 (Student’s t-test (E), two-way ANOVA followed by Bonferroni’s post-tests (A)).

**Supplementary Figure S9**

**Fig. S9. Collagen type I, fibronectin, periostin, MFAP4, collagen and elastin fibers in normal human skin, tumors and wounds.**

(A-D) Immunofluorescence co-staining for fibronectin (A-C, red) or collagen type I (D, red) and MFAP4 (green) in sections of human superficial (A) and deep skin (B), subcutaneous melanoma edge (C-D); nuclei are counterstained with DAPI (blue). Yellow dashed lines represent epidermal-dermal junctions (A-B) or tumor margins (C-D). Individual channels are shown in black and white. (E, F) Picrosirius red staining for collagen of human superficial (E, left and middle) and deep skin (E, right), subcutaneous melanoma core (F), visualized in brightfield (left) or polarized light (middle, right). Scale bars = 200 μm.

(G) Visualization via multiphoton microscopy by second-harmonic generation (SHG) of collagen (red) and auto-fluorescence (AF) for elastin (green) in a section of human superficial and deep skin stained with an antibody against MFAP4 (cyan) and counter-stained with DAPI (blue), showing individual channels in black and white.

(H) Inset detail of the area marked by the white square in (G), showing significant fibrous co-localization of elastin AF signal (middle) and MFAP4 staining (right) in the dermis. Scale bars = 200 μm.

(I-L) Visualization via multiphoton microscopy by second-harmonic generation (SHG) for collagen (red) and auto-fluorescence (AF) for elastin (green) of subcutaneous melanoma (I), BCC (J), and human excisional wound sections from 5-, 8-, 12-, and 15-days post-injury (K-L). Insets (white boxes in zoomed out images) show detailed organization of collagen and elastin fibers at the tumor margins (I-J) and at the edges of wounds (K, insets in L); individual channels are shown in black and white. Scale bars = 200 μm. Black or white dashed lines represent epidermal-dermal junctions (A, B, G, I) or tumor margins (C, D, I, J).

**Supplementary Figure S10**

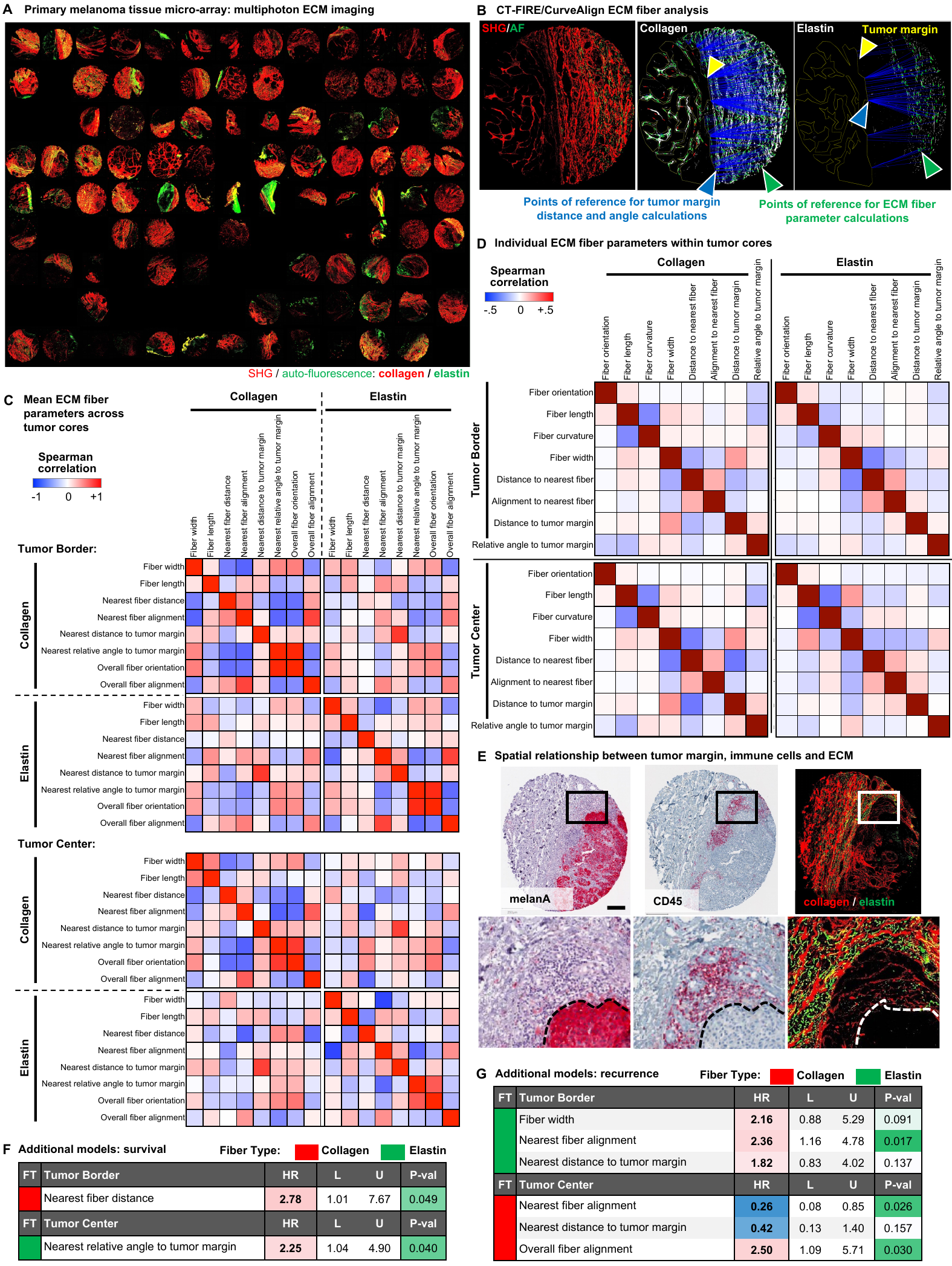

**Fig. S10. Collagen/elastin imaging and ECM fiber analysis of primary melanoma tissue microarray.**

(A) Composite image showing the primary melanoma tissue micro-array (TMA) of tumor cores individually imaged via multiphoton microscopy by second-harmonic generation (SHG) for collagen (red) and auto-fluorescence (AF) for elastin (green).

(B) Diagram showing the major features of (ECM fiber analysis of tumor cores by CT-FIRE/CurveAlign, taking into account the points of reference for calculation of ECM fiber parameters, especially those that are relative to the manually-drawn borders of the tumor margin within each tumor core.

(C) Spearman correlation matrix of mean ECM fiber parameters across tumor cores for collagen and elastin fibers at the tumor border (top) or the tumor center (bottom).

(D) Spearman correlation matrix of individual ECM fiber parameters within individual tumor cores for collagen (left) or elastin (right) fibers at the tumor border (top) or the tumor center (bottom).

(E) Representative melanoma tumor border core, showing sections stained for tumor cells (melanA, left), immune cells (CD45, center) and ECM fibers (collagen/elastin, right). Inset shows the spatial relationship between the tumor margin, immune cell infiltrate, and high density of elastin fibers (green).

(F, G) Additional Multivariate Cox regression models of ECM fiber parameters significantly impacting patient survival (F) or melanoma recurrence (G), showing the collagen (red) and elastin (green) fiber parameters contributing to statistically-significant hazard ratios (HR) at the tumor border (top) and center (bottom).

Black or white dashed lines represent tumor margins. FT: fiber type; P-val: P-value; L: lower bound; U: upper bound.

**Supplementary Figure S11 (Summary Figure)**

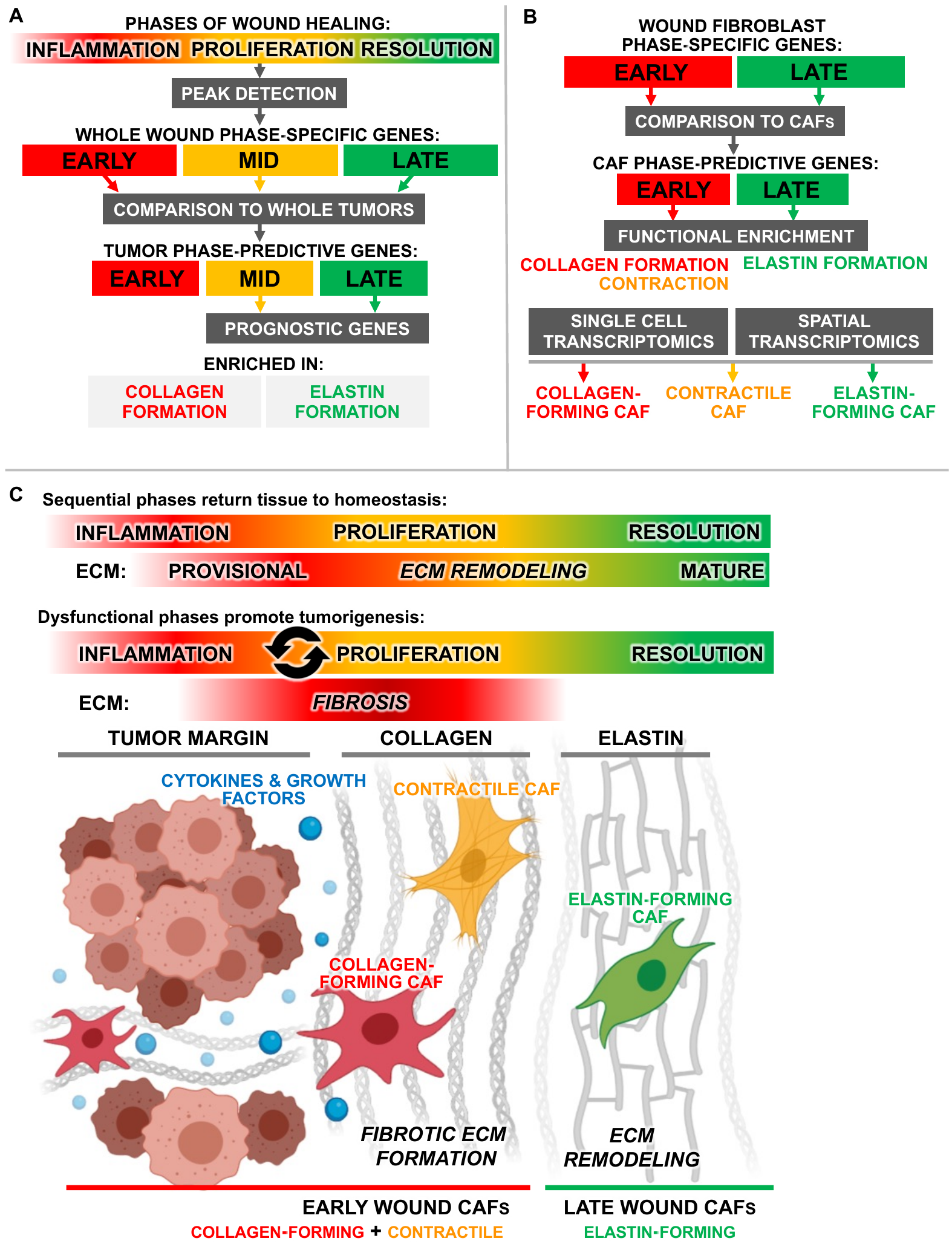

**Fig S11. Summary figure.**

(A) Diagram of the whole tissue bioinformatics pipeline identifying healing phase-specific genes from analysis of the wound healing spectrum and tumor phase-predictive genes from comparisons to pan-cancer datasets, and focusing on the most prognostic genes and pathways within the mid and late phases of healing, which enriched for collagen and elastin formation.

(B) Diagram of the fibroblast-centric bioinformatics pipeline identifying wound fibroblast phase-specific genes and CAF phase-predictive genes from comparisons to pan-CAF datasets, which enriched for contraction and collagen formation in the “early” group and for elastin formation in the “late” group. ScRNA-seq meta-analyses of wound fibroblast and CAF datasets confirmed the existence of these functionally distinct “early” and “late” CAF subtypes, and re-analyses of spatial transcriptomics cancer datasets confirmed the presence of these CAF subtypes in distinct niches of the TME.

(C) Conceptual summary of the manuscript: During wound healing, sequential healing phases return the injured tissue to homeostasis. As a “wound that fails to heal”, cancer stalls in the early phases of repair, with cancer cells and TME cells continually releasing cytokines and growth factors to exacerbate the early wound phenotypes, including ECM formation, resulting in fibrosis. Typical wound fibroblast functions of the early healing phase become hyper-activated in “early-wound” CAF subtypes in close proximity to the tumor margins, contributing to the accumulation of pro-tumorigenic collagen-forming and contractile CAFs that exacerbate the fibrotic ECM microenvironment. Further away from the tumor margins, there is a gradient toward the normalization of the tumor-adjacent microenvironment mediated by “late-wound” CAF subtypes that deposit an elastin-enriched ECM network similar to that found in a resolving wound.

**Table S1. List and description of datasets analyzed in this manuscript.**

| **Bulk tissue meta-analysis of wounds and tumors** | | | | | |
| --- | --- | --- | --- | --- | --- |
| Data category | Data description | Organism | Sample type | Accession number | Reference (PMID) |
| Meta-analysis of various wounded tissues and models | Meta-analysis comparing gene expression profiles in response to wounding in heart, liver, skin, bones, and spinal cord of rat, mouse and human | Various | Microarray, bulk tissue | N/A | 29115927 |
| Whole wound transcriptome | 3-mm diameter excisional, full-thickness skin wounds at 0-day (intact back skin), 1-day, 4-day, 8-day and 14-day post-injury | Mouse | RNA-seq, bulk tissue | GSE72964 | 27462873 |
| Whole wound transcriptome | 1-mm diameter excisional, full-thickness skin wounds at 0-day (intact back skin), 12h, 24h, 3-day, 5-day, 7-day and 10-day post-injury | Mouse | Microarray, bulk tissue | GSE23006 | 32340108 |
| Whole wound transcriptome | 3-mm diameter excisional, paired and sequential oral and skin wound biopsies at 1-day (unwounded tissue), 3-day and 6-day post-injury | Human | RNA-seq, bulk tissue | GSE97615 | 30045979 |
| Whole skin graft transcriptome | Biopsies of split-thickness skin graft donor site wounds at 0-day (intact skin), 3-day and 7-days after implantation | Human | Microarray, bulk tissue | GSE28914 | 23082929 |
| Whole skin graft transcriptome | Biopsies of split-thickness skin graft donor site wounds at 0-day (intact skin), 14-day and 21-day after implantation | Human | Microarray, bulk tissue | GSE50425 | N/A |
| Whole tumor transcriptome (various solid cancer types) | Normalized count data from the The Cancer Genome Atlas (TCGA) and Genome-Tissue Expression Project (GTEX) databases (TARGET data was removed), filtered for matching tissues of origin, primary site tumors, non-blood and non-lymphatic tissues | Human | RNA-seq, bulk tissue | TcgaTargetGtex_RSEM_Hugo_norm_count | N/A |
| Whole tumor transcriptome (skin basal cell carcinoma) | Biopsies of intact skin, treatment-resistant basal cell cancers and treatment-sensitive basal cell cancers | Human | RNA-seq, bulk tissue | GSE58375 | 25759020 |
| Whole tumor transcriptome (actinic keratosis and skin squamous cell carcinoma) | Biopsies of intact skin, actinic keratosis and cutaneous squamous cell carcinomas | Human | RNA-seq, bulk tissue | GSE84293 | 27574101 |
| **Gene set enrichment analysis of wound fibroblasts, treated fibroblasts, and tumor stroma** | | | | | |
| Data category | Data description | Organism | Sample type | Accession number | Reference (PMID) |
| Wound fibroblast transcriptome | FACS-sorted fibroblasts from 5-mm diameter excisional, full-thickness skin wounds at 0-day (normal skin) and 5-day post-injury | Mouse | RNA-seq, bulk sorted cells | GSE134789 | 32451392 |
| CAF transcriptome (skin cancer) | FACS-sorted fibroblasts from established back skin tumors of K14-HPV8 mice and age-matched back skin controls | Mouse | RNA-seq, bulk sorted cells | GSE182880 | This manuscript |
| Wound fibroblast transcriptome | FACS-sorted fibroblasts from 2.25 cm^2^ excisional, full-thickness skin wounds at 12-day and 26-day post-injury | Mouse | RNA-seq, bulk sorted cells | GSE84256 | 28059714 |
| Wound fibroblast transcriptome | FACS-sorted ɑ-SMA^+^ myofibroblasts from small excisional wounds at 7-day post-injury | Mouse | RNA-seq, bulk sorted cells | GSE95786 | 28797711 |
| Wound fibroblast transcriptome | FACS-sorted CD29^high^, CD29^low^, or adipocyte precursor myofibroblasts from small excisional wounds at 5-day post-injury | Mouse | RNA-seq, bulk sorted cells | GSE105790 | 30467144 |
| Wound fibroblast transcriptome | Single cells, including fibroblasts, isolated from 1.5 x 1.5 cm^2^ excisional, full-thickness skin wounds at 12-day post-injury | Mouse | RNA-seq, single cells | GSE113854 | 30737373 |
| Treated skin fibroblast transcriptome | Skin fibroblasts treated with serum for 6 hours | Human | Microarray, bulk cultured cells | GSE21648 | 21098492 |
| Treated skin fibroblast transcriptome | Skin fibroblasts treated with TGF-β for 24 hours | Human | Microarray, bulk cultured cells | GSE79621 | 27256716 |
| Treated skin fibroblast transcriptome | Skin fibroblasts treated with LPS for 24 hours | Human | Microarray, bulk cultured cells | GSE79621 | 27256716 |
| Cancer stroma transcriptome (breast cancer) | Laser micro-dissected tumor stroma from invasive breast carcinoma biopsies | Human | Microarray, laser micro-dissected tissue | GSE26910 | 21611158 |
| Cancer stroma transcriptome (prostate cancer) | Laser micro-dissected tumor stroma from invasive prostate carcinoma biopsies | Human | Microarray, laser micro-dissected tissue | GSE26910 | 21611158 |
| Cancer stroma transcriptome (breast cancer) | Laser micro-dissected tumor stroma from pregnancy associated breast cancer biopsies | Human | Microarray, laser micro-dissected tissue | GSE31192 | 23479404 |
| Cancer stroma transcriptome (liver cancer) | Laser micro-dissected tumor stroma from intrahepatic cholangiocarcinoma biopsies | Human | Microarray, laser micro-dissected tissue | GSE45001 | 23775819 |
| Cancer stroma transcriptome (lung cancer) | Laser micro-dissected tumor stroma from non-small cell lung carcinoma biopsies | Human | Microarray, laser micro-dissected tissue | GSE22874 | 21474781 |
| **Bulk cell meta-analysis of wound- and cancer-associated fibroblasts** | | | | | |
| Data category | Data description | Organism | Sample type | Accession number | Reference (PMID) |
| Wound fibroblast transcriptome | FACS-sorted fibroblasts from 5-mm diameter excisional, full-thickness skin wounds at 0-day (normal skin) and 5-day post-injury | Mouse | RNA-seq, bulk sorted cells | GSE134789 | 32451392 |
| Wound fibroblast transcriptome | FACS-sorted fibroblasts from 2.25 cm^2^ excisional, full-thickness skin wounds at 12-day, 15-day, 21-day and 26-day post-injury | Mouse | RNA-seq, bulk sorted cells | GSE84256 | 28059714 |
| CAF transcriptome (skin cancer) | FACS-sorted fibroblasts from established back skin tumors of K14-HPV8 mice and age-matched back skin controls | Mouse | RNA-seq, bulk sorted cells | GSE182880 | This manuscript |
| CAF transcriptome (skin cancer) | FACS-sorted fibroblasts from dysplastic skin tissue of K14-HPV16 mice, and from age-matched non-transgenic controls | Mouse | Microarray, bulk sorted cells | GSE17817 | 20138012 |
| CAF transcriptome (skin cancer) | Cultured fibroblasts isolated from individuals without cutaneous squamous cell carcinoma and directly from tumor matrix in recessive dystrophic epidermolysis bullosa (RDEB) and non-RDEB samples | Human | Microarray, bulk cultured cells | GSE37738 | 22564523 |
| CAF transcriptome (breast cancer) | Cultured fibroblasts isolated from primary breast cancer, metastatic brain lesions, and normal breast tissue | Human | RNA-seq, bulk cultured cells | GSE80333 | 27671678 |
| CAF transcriptome (breast cancer) | Cultured fibroblasts isolated from breast carcinoma and matched normal breast tissue | Human | Microarray, bulk cultured cells | GSE20086 | 20062080 |
| CAF transcriptome (head and neck cancer) | Cultured fibroblasts isolated from head and neck squamous cell carcinoma (HNSSC) and matched normal skin tissue | Human | RNA-seq, bulk cultured cells | GSE83314 | 27488962 |
| CAF transcriptome (lung cancer) | Cultured fibroblasts isolated from non-small cell lung carcinoma (NSCLC) and matched normal lung tissue | Human | Microarray, bulk cultured cells | GSE22862 | 21474781 |
| CAF transcriptome (colorectal cancer) | Cultured fibroblasts isolated from primary colorectal carcinoma and matched normal colonic mucosa | Human | Microarray, bulk cultured cells | GSE46824 | N/A |
| CAF transcriptome (prostate cancer) | Cultured fibroblasts isolated from primary prostate cancer and normal prostate tissue | Human | Microarray, bulk cultured cells | GSE68164 | 26375444 |
| CAF transcriptome (pancreatic cancer) | Cultured fibroblasts isolated from spontaneous pancreatic ductal adenocarcinomas and normal pancreatic tissue of transgenic animals | Mouse | RNA-seq, bulk cultured cells | GSE55870 | 24856586 |
| **Single cell meta-analysis of wound- and cancer-associated fibroblasts** | | | | | |
| Data category | Data description | Organism | Sample type | Accession number | Reference (PMID) |
| Wound fibroblast transcriptome | Single cells, including fibroblasts, isolated from 6 mm diameter excisional, full-thickness skin wounds at 4-day post-injury | Mouse | RNA-seq, single cells | GSE142471 | 32187560 |
| Wound fibroblast transcriptome | Single cells, including fibroblasts, isolated from 1.5 x 1.5 cm_2_ excisional, full-thickness skin wounds at 12-day post-injury | Mouse | RNA-seq, single cells | GSE113854 | 30737373 |
| Wound fibroblast transcriptome | Single cells, including fibroblasts, isolated from 1.2 x 1.2 cm^2^ excisional, full-thickness skin wounds at 18-day post-injury; wounds had two potential outcomes: fibrotic and regenerative | Mouse | RNA-seq, single cells | GSE141814 | 32219160 |
| CAF transcriptome (melanoma) | Isolated cells, including fibroblasts, from primary tumors and lymph nodes during subcutaneous melanoma formation in animals injected with B16-F10 melanoma cells | Mouse | RNA-seq, single cells | E-MTAB-7427 | 32433953 |
| CAF transcriptome (skin BCC) | Isolated cells, including fibroblasts, from basal cell carcinoma tumors | Human | RNA-seq, single cells | GSE123813 | 31359002 |
| CAF transcriptome (skin SCC) | Isolated cells, including fibroblasts, from primary cutaneous squamous cell carcinoma tumors and normal skin | Human | RNA-seq, single cells | GSE144236 | 32579974 |
| CAF transcriptome (head and neck cancer) | Isolated cells, including fibroblasts, from primary head and neck squamous cell carcinoma (HNSSC) tumors | Human | RNA-seq, single cells | GSE103322 | 29198524 |
| CAF transcriptome (lung cancer) | Isolated cells, including fibroblasts, from primary lung cancer tumors and normal lung tissue | Human | RNA-seq, single cells | E-MTAB-6149 | 32561858 |
| CAF transcriptome (breast cancer) | Isolated cells, including fibroblasts, from primary breast cancer tumors and normal breast tissue | Human | RNA-seq, single cells | E-MTAB-8107 | 32561858 |
| CAF transcriptome (colorectal cancer) | Isolated cells, including fibroblasts, from primary colorectal cancer tumors and normal colonic mucosa | Human | RNA-seq, single cells | E-MTAB-8107 | 32561858 |
| CAF transcriptome (ovarian cancer) | Isolated cells, including fibroblasts, from primary ovarian cancer tumors and normal ovarian tissue | Human | RNA-seq, single cells | E-MTAB-8107 | 32561858 |
| CAF transcriptome (skin melanoma) | Isolated cells, including fibroblasts, from primary skin melanoma lesions and metastatic lymph nodes | Human | RNA-seq, single cells | GSE72056 | 27124452 |
| **Spatial transcriptomics of tumor stroma** | | | | | |
| Data category | Data description | Organism | Sample type | Accession number | Reference (PMID) |
| Skin cancer | Sections of cutaneous squamous cell carcinoma | Human | RNA-seq, spatial | GSE144239 | 32579974 |

**Table S12. Description of fibroblast cell lines used in experiments.**

| **Full Name** | **Species** | **Primary / Immortalized** | **Cell Line Name** | **Condition** | **Serves as…** |
| --- | --- | --- | --- | --- | --- |
| mImmFB #1 EV | mouse | immortalized | mImmFB #1 | Empty Vector | control |
| mImmFB #1 shRunx2 B8 | mouse | immortalized | mImmFB #1 | shRunx2 | knock-down cells |
| mImmFb #2 SC | mouse | immortalized | mImmFB #2 | Scrambled shRNA | control |
| mImmFB #2 shRunx2 B8 | mouse | immortalized | mImmFB #2 | shRunx2 | knock-down cells |
| mImmFB #3 EV | mouse | immortalized | mImmFB #3 | Empty Vector | control |
| mImmFB #3 shRunx2 B8 | mouse | immortalized | mImmFB #3 | shRunx2 | knock-down cells |
| hpFB SC | human | primary | hpFB | Scrambled shRNA | control |
| hpFB shRunx2 C9 | human | primary | hpFB | shRunx2 | knock-down cells |

**Table S14. Patient demographics of analyzed samples from primary melanoma tissue microarray.**

| **Variable** | **Median or Number (range or %)** |
| --- | --- |
| **Age at diagnosis** | 65 (29-79) |
| **Gender** |  |
| Female | 12 (36.36%) |
| Male | 21 (63.64%) |
| **Histological subtype** |  |
| Superficial spreading | 10 (30.30%) |
| Nodular | 16 (48.48%) |
| Other | 7 (21.21%) |
| **Months follow-up** | 51 (1-112) |
| **Breslow thickness** | 3.20 (0.80-12.50) |
| **Ulceration** |  |
| Yes | 22 (66.67%) |
| No | 11 (33.33%) |
| **Recurrence** |  |
| Yes | 10 (30.30%) |
| No | 23 (69.70%) |
| **Death** |  |
| Yes | 9 (27.27%) |
| No | 24 (72.73%) |
